# *In silico* drug repurposing and in vitro validation of cestode fatty acid binding proteins

**DOI:** 10.64898/2026.03.04.709508

**Authors:** Santiago Rodríguez, Lucas N. Alberca, Luciana Gavernet, Gisela Franchini, Alan Talevi

## Abstract

Echinococcosis is a Neglected Tropical Disease (NTD) caused by *Echinococcus granulosus* and *Echinococcus multilocularis*, the etiological agents of cystic and alveolar echinococcosis, respectively. These infections pose a significant public health burden, particularly in endemic regions. Cestodes lack key enzymes involved in lipid metabolism and must acquire lipids from their hosts. Fatty Acid Binding Proteins (FABPs), which mediate lipid trafficking and intracellular transport, have therefore emerged as essential and potentially druggable targets.

In this study, we implemented an integrated virtual screening strategy combining ligand-based and structure-based approaches to identify novel FABP binders as potential therapeutic agents against Echinococcus spp. High-specificity screening of approximately 435,000 compounds yielded a limited number of prioritized *in silico* hits. Four compounds—hydrochlorothiazide, naratriptan, fenticonazole, and montelukast—were selected for experimental validation, prioritizing repurposing candidates.

Fluorescence displacement assays confirmed that hydrochlorothiazide binds to three cestode FABPs (EgFABP1, EmFABP1, and EmFABP3), validating the predictive performance of the computational workflow. These findings support the value of parallel *in silico* screening strategies and drug repurposing approaches for the discovery of new therapeutic candidates against neglected tropical diseases.

## 1 Introduction

Echinococcosis is listed by the World Health Organization (WHO) among the top twenty neglected tropical diseases (NTDs). In particular, cystic echinococcosis is a serious zoonotic disease in South America [1,2]. Infected dogs harbor adult worms in their intestines, where they lay eggs that are subsequently released in the feces. The eggs can then be ingested by herbivores (e.g., livestock) or humans, who act as potential intermediate hosts. The larval stage develops through asexual reproduction in the internal organs of the intermediate host, progressively increasing intracystic pressure.

To date, benzimidazoles have been the primary medical treatment for echinococcosis, although combination therapy with praziquantel has shown higher scolicidal activity than benzimidazoles alone [3]. However, these medications are often poorly tolerated [4], and the efficacy of benzimidazoles depends on the type, size, and location of the cyst [5]. Therefore, the development of new anthelmintic agents remains a priority.

Tsai and coworkers reported that genes encoding enzymes for *de novo* fatty acid and cholesterol biosynthesis are absent in tapeworms, whereas genes involved in lipid trafficking, such as fatty acid binding proteins (FABPs), are highly expressed [6]. FABPs are small intracellular proteins (14–15 kDa) that bind fatty acids as well as other hydrophobic ligands, including sterols [7]. They are widely distributed throughout the animal kingdom and are highly expressed in cells with active lipid metabolism [8]. A previous study identified six FABP-coding genes in *Echinococcus spp*. In *E. multilocularis*, two of these genes encode identical protein sequences, whereas in *E. granulosus* all coding sequences are distinct [9]. Functional characterization of EgFABP1, a FABP isoform from *E. granulosus*, demonstrated its ability to bind and transport fatty acids between artificial membranes [10]. EgFABP1 is currently the only cestode FABP isoform with a resolved tertiary structure available in the Protein Data Bank (PDB ID: 1O8V). Furthermore, FABPs have been proposed as potential drug targets for the development of new therapeutic strategies against platyhelminths [11].

Virtual screening (VS) encompasses a range of computational techniques used to screen large chemical libraries and identify potential drug candidates [12,13]. VS approaches are generally classified as ligand-based or structure-based methods, which can be applied in a complementary manner to offset their respective limitations [12,13]. In this study, we combined ligand- and structure-based models to identify potential inhibitors of cestode FABPs through VS. The candidate compounds identified in both campaigns were subsequently subjected to experimental validation to confirm the computational predictions.

## 2 Results and discussion

### 2.1 Generation and Validation of Linear Models

4,000 ligand-based linear models were generated using a random subspace approach applied to Mordred descriptors, yielding linear classifiers capable of discriminating between ACTIVE and INACTIVE compounds against cestode FABPs. Equation 1 presents the best-performing model in RL1 (model 3963):

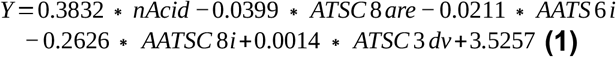

where *nAcid* represents the number of acidic groups in the molecule; *ATSC*8*are* corresponds to the centered Broto–Moreau autocorrelation coefficient of order 8 weighted by Allred–Rochow electronegativity; *AATS*6*i* is the averaged Broto–Moreau autocorrelation coefficient of order 6 weighted by ionization potential; *AATSC*8*i* refers to the centered and averaged Broto–Moreau autocorrelation coefficient of order 8 weighted by ionization potential; and *ATSC*3*dv* corresponds to the centered Broto– Moreau autocorrelation coefficient of order 3 weighted by valence electrons [14,15]. It can be observed that most of the features included in the model correspond to 2D autocorrelation descriptors, which are intuitively related to the pharmacophore concept [12,16], as they reflect the molecular distribution of atomic properties relevant for ligand–target interactions, such as ionization potential and electronegativity. Furthermore, the presence of acidic groups is directly correlated with the ability of a ligand to bind FABPs. This observation may be related to the fact that the physiological substrates of these proteins are fatty acids containing free acidic functions [17].

Ensemble models were constructed using the operators MIN, PROD, AVE, VOTE, and RANK. The BEDROC metric was employed to evaluate early enrichment performance, and ensembles were selected by balancing BEDROC values against the number of combined models. This evaluation was conducted using the RL1-A dataset, whereas RL1-B was reserved for independent validation (see section 5.1 for the details of dataset composition).

As shown in **Figure 1**, the MIN operator achieved the best BEDROC performance, leading to the selection of a 32-model ensemble (MIN32) for use in the prospective VS campaign. MIN32 was preferred over other similarly performing ensembles because it achieved comparable performance while requiring fewer constituent models, in agreement with the principle of parsimony. In addition, MIN32 significantly outperformed the best individual model in terms of BEDROC in both RL1-A (0.632 ± 0.041 vs. 0.255 ± 0.032) and RL1-B (0.603 ± 0.034 vs. 0.273 ± 0.038). The detailed results are summarized in **Table 1**.

**Table 1.** AUC-ROC and BEDROC values (𝛼=100) for the best-performing individual model (model 3963) and the MIN32 ensemble.

| MODEL | RL1-A<br>AUC-ROC<br>(+/- SD) | RL1-A<br>BEDROC<br>(+/- SD) | RL1-B<br>AUC-ROC<br>(+/- SD) | RL1-B<br>BEDROC<br>(+/- SD) |
| --- | --- | --- | --- | --- |
| 3963 | 0.932<br>(+/- 0.015) | 0.255<br>(+/- 0.032) | 0.954<br>(+/- 0.005) | 0.273<br>(+/- 0.038) |
| MIN32 | 0.945<br>(+/- 0.015) | 0.632<br>(+/- 0.041) | 0.935<br>(+/- 0.013) | 0.603<br>(+/- 0.034) |

**Figure 1.**
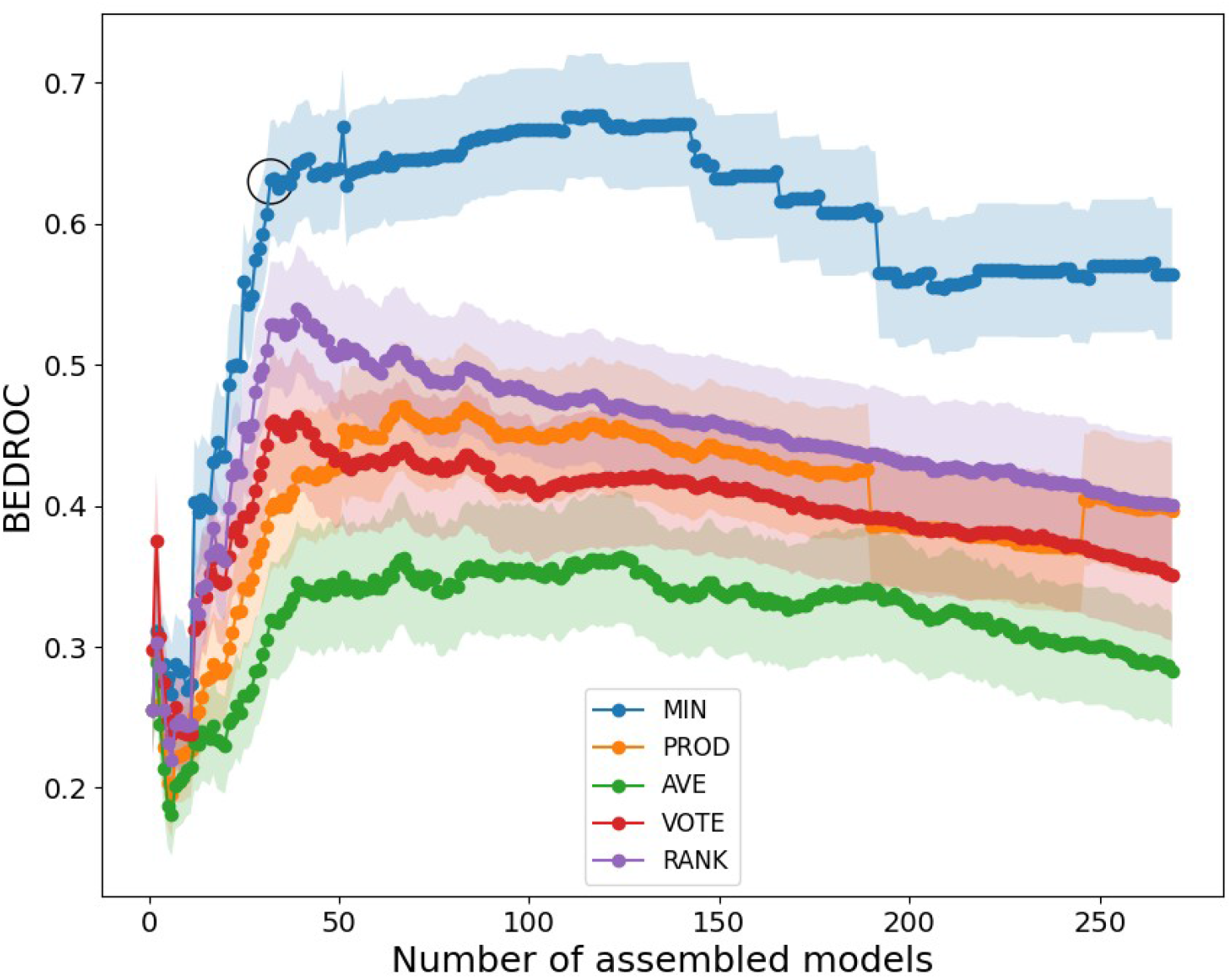
Variation of the BEDROC metric (𝛼 = 100) as a function of the number of combined models for each of the evaluated operators. The shaded area around each curve represents the standard deviation at each point, estimated through 1,000 rounds of bootstrapping. The MIN32 ensemble, selected for the prospective virtual screening campaign, is indicated by a circle.

To validate the robustness and predictive power of the models included in MIN32, Y-randomization and LGO cross-validation were performed. In the first case, the average accuracy of the randomly generated models was significantly lower than that of the original models, suggesting a low probability of spurious correlations. In the LGO procedure, the mean accuracies of the re-trained models were very similar to those obtained with the original models, with differences falling within ± one standard deviation of the mean accuracy estimated across the LGO rounds. The results of both Y-randomization and LGO validations confirmed the robustness and non-spurious nature of the models. Model details and validation metrics are provided in the Supplementary Information (**List S1, Table S1**).

A PPV (Predicted Positive Value) surface was constructed using the RL1-B dataset to define a score threshold for prospective VS. PPV estimates the proportion of compounds predicted as ACTIVE by the models (*in silico* hits) that are expected to be experimentally confirmed. A threshold of 0.400 was selected, yielding Sp = 0.994, Se = 0.455, and PPV = 0.455 (proportion of actives (Ya) = 0.01). This implies approximately a 50% probability of true positives among the selected compounds, assuming a 1% hit rate, and about 30% if a Ya of 0.5% is assumed. Similar results were obtained for the RL2 dataset (Sp = 0.995, Se = 0.455, PPV = 0.496). **Table 2** summarizes these results, while the PPV surfaces for both libraries are presented in the Supplementary Information (**Figures S1** and **S2**).

**Table 2.** Se, Sp, and PPV values corresponding to the selected cutoff in the retrospective ligand-based VS. Values in parentheses indicate the proportion of ACTIVE compounds assumed for the PPV calculation.

| LIBRARY | CUTOFF VALUE | Se | Sp | PPV (0.001) | PPV (0.005) | PPV (0.01) |
| --- | --- | --- | --- | --- | --- | --- |
| RL1-B | 0.400 | 0.455 | 0.994 | 0.076 | 0.294 | 0.455 |
| RL2 | 0.400 | 0.455 | 0.995 | 0.089 | 0.328 | 0.496 |

### 2.2 Prospective Ligand-Based VS

The chemical databases listed in the computational methods section were screened. As a result, 56 out of approximately 435,000 compounds were classified as ACTIVE (Supplementary Information, **List S2**) and **fell within the applicability domain of the selected models**. Among these hit compounds, 14 are currently undergoing clinical trials for the treatment of other pathologies. Based on **commercial availability and cost considerations**, naratriptan HCl, fenticonazole nitrate, and hydrochlorothiazide were acquired for subsequent experimental validation.

### 2.3 Molecular Dynamics and WATCLUST Algorithm

Molecular dynamics (MD) simulations were performed on Apo-EgFABP1 to identify water sites (WS). Analysis of the trajectory using WATCLUST revealed three distinct water clusters located near the binding triad of cestode FABPs (57), in proximity to the crystallographic water molecules observed in the target structure (**Figure 2**). The Water Finding Probability (WFP) calculated using WATCLUST indicated that the probability of water occupancy in each of these clusters was approximately fivefold higher than in bulk solvent (5.62, 4.83, and 4.95 for each WS). The coordinates of the identified WS were subsequently used to guide docking calculations using the Autodock (AD) Bias algorithm, as described in the computational methods (section 5.5).

**Figure 2.**
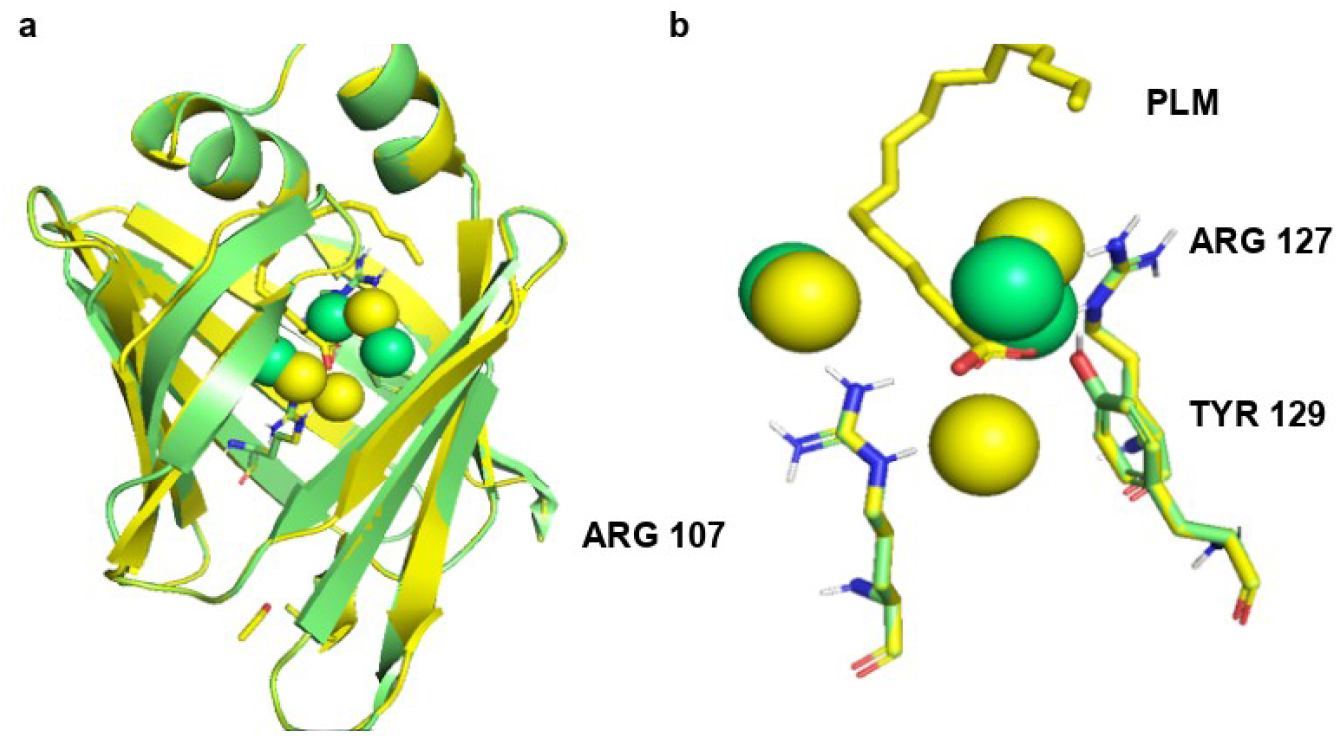
(a) Crystal structure of EgFABP1 (shown in yellow) superimposed with the structure obtained from MD simulation (shown in green). The binding triad and the ligand are highlighted as sticks, while water molecules are represented as spheres. The predicted WS are depicted in yellow. (b) Superimposition of the crystallographic water molecules and the FABP binding triad with the WS predicted by WATCLUST. PLM: Palmitic acid.

### 2.4 Molecular Docking, Score Threshold Selection, and Prospective VS Campaign

The screening performance of different docking methods was evaluated using EgFABP1 as the receptor. In the specific case of AD Bias, the three WS identified by MD were incorporated into the protocol. These were modeled as hydrogen-bond donor (D) or hydrogen-bond acceptor (A) biases, considering all possible combinations. The scores of the top-ranked poses obtained for compounds in the validation sets were used to calculate the AUC-ROC for each docking protocol. AD Bias with three donor biases (DDD) demonstrated the best performance, yielding higher AUCROC values than the remaining algorithms in both the dRL1 and dRL2 libraries (0.7670 ± 0.0080 and 0.9030 ± 0.0059, respectively; Supplementary Information; **Table S2**). Moreover, switching the bias class from donor to acceptor negatively affected the AUCROC values, particularly when the substitution involved WS2 and, to a lesser extent, WS1 and WS3. Conversely, converting the bias class from acceptor to donor resulted in improved AUCROC values (**Table 3**).

**Table 3.**
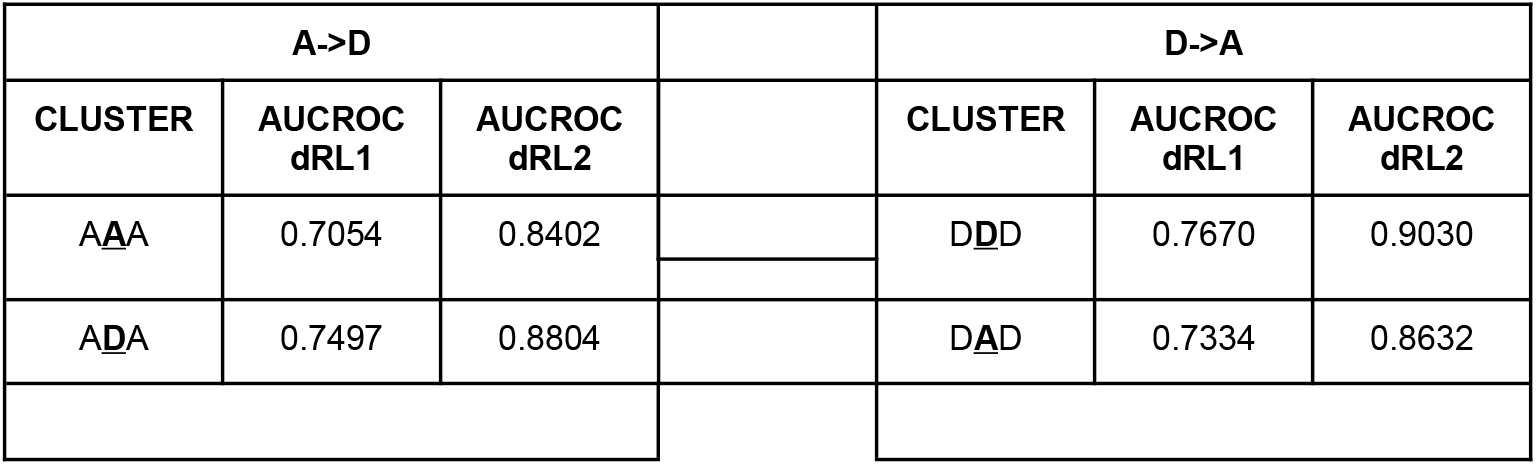
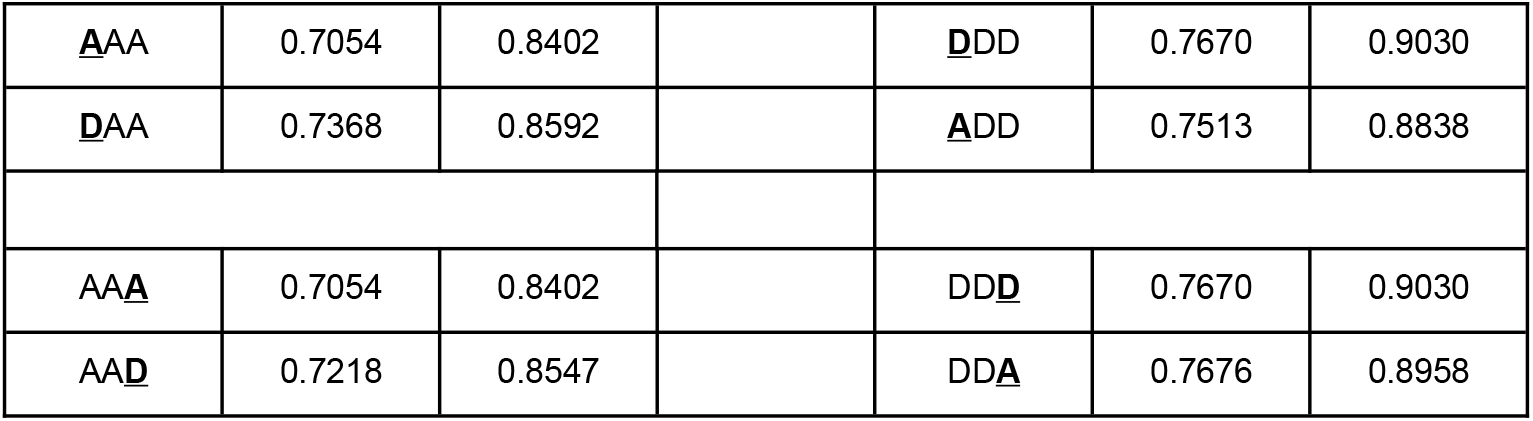
Variation in AUC-ROC values upon switching acceptor/donor bias according to WS position.

Based on these results, the third WS donor bias was removed (DD-scheme), followed sequentially by the removal of the first one (-D-), and the effects on AUCROC values were compared. Docking performance improved in both the dRL1 and dRL2 datasets when a single donor bias was retained at WS2 (-D-), as reflected by increased AUCROC values (0.7794 ± 0.0077 and 0.9052, respectively; see **Table 4**). Therefore, this restriction was incorporated into the prospective structure-based virtual screening campaign. The location of the donor bias within the binding site is shown in **Figure 3**.

**Table 4.** AUCROC values in the dRL1 and dRL2 libraries obtained using one, two, or three donor biases. *Significant differences between AD Bias (-D-) and the different docking algorithms in the dRL1 library (Tukey’s test, α = 0.05).

|  | dRL1<br>(+/- SD) | dRL2<br>(+/- SD) |
| --- | --- | --- |
| **DDD** | 0.7670<br>(+/- 0.0080) | 0.9030<br>(+/- 0.0059) |
| **DD-** | 0.7696*<br>(+/- 0.0079) | 0.9011<br>(+/- 0.0058) |
| **-D-** | 0.7794<br>(+/- 0.0077) | 0.9052<br>(+/- 0.0056) |

**Figure 3.**
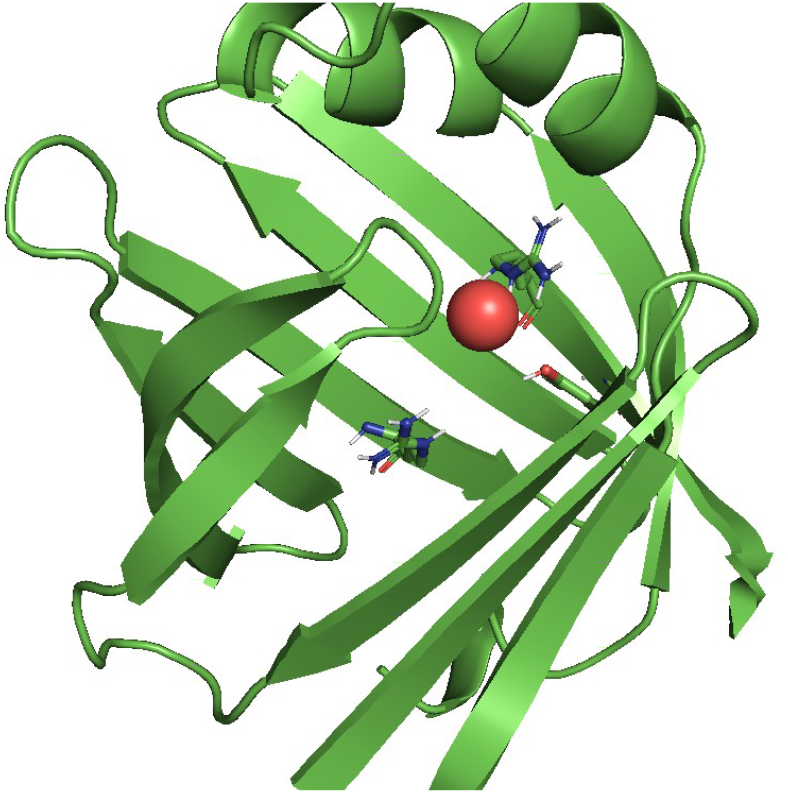
Docking bias (-D-) shown in the active site as a red sphere located **between residues Arg127 and Tyr129**.

PPV surface analysis of the AD Bias (-D-) protocol was performed using the dRL1 and dRL2 datasets. A threshold of –12.65 was selected by prioritizing Sp over Se, following the same criterion applied in the ligand-based approach. Compounds scoring equal to or more negative than this threshold were classified as ACTIVE. The resulting metrics were Sp = 0.999, Se = 0.043, and PPV = 0.292 for dRL1, and Sp = 0.999, Se = 0.043, and PPV = 0.709 for dRL2. These results indicate a 30– 70% probability of correctly identifying ACTIVE compounds under a 1% hit-rate scenario. The results are summarized in **Table 5**, and the PPV surfaces are provided in Supplementary **Figures S3** and **S4**.

**Table 5.** Se, Sp, and PPV values corresponding to the selected cutoff score. Values in parentheses indicate the proportion of ACTIVE compounds assumed for the PPV calculation.

| LIBRARY | CUTOFF VALUE | Se | Sp | PPV (0.001) | PPV (0.005) | PPV (0.01) |
| --- | --- | --- | --- | --- | --- | --- |
| dRL1 | -12.65 | 0.043 | 0.999 | 0.039 | 0.170 | 0.292 |
| dRL2 | -12.65 | 0.043 | 0.999 | 0.194 | 0.548 | 0.709 |

Using the -D-docking model, we screened the same chemical databases described for the ligand-based VS, getting 98 out of 435,000 compounds predicted as ACTIVE (Supplementary Information, **List S3**). None of these ACTIVE compounds had been found by the ligand-based models, suggesting both approaches may lead to complementary lists of *in silico* hits. Among these docking hits, two have been in clinical trials for other pathologies. Based on the cost and availability for acquisition, we selected montelukast for experimental evaluation.

The chemical structures of the four molecules submitted to experimental validation can be shown in **Figure 4**.

**Figure 4.**
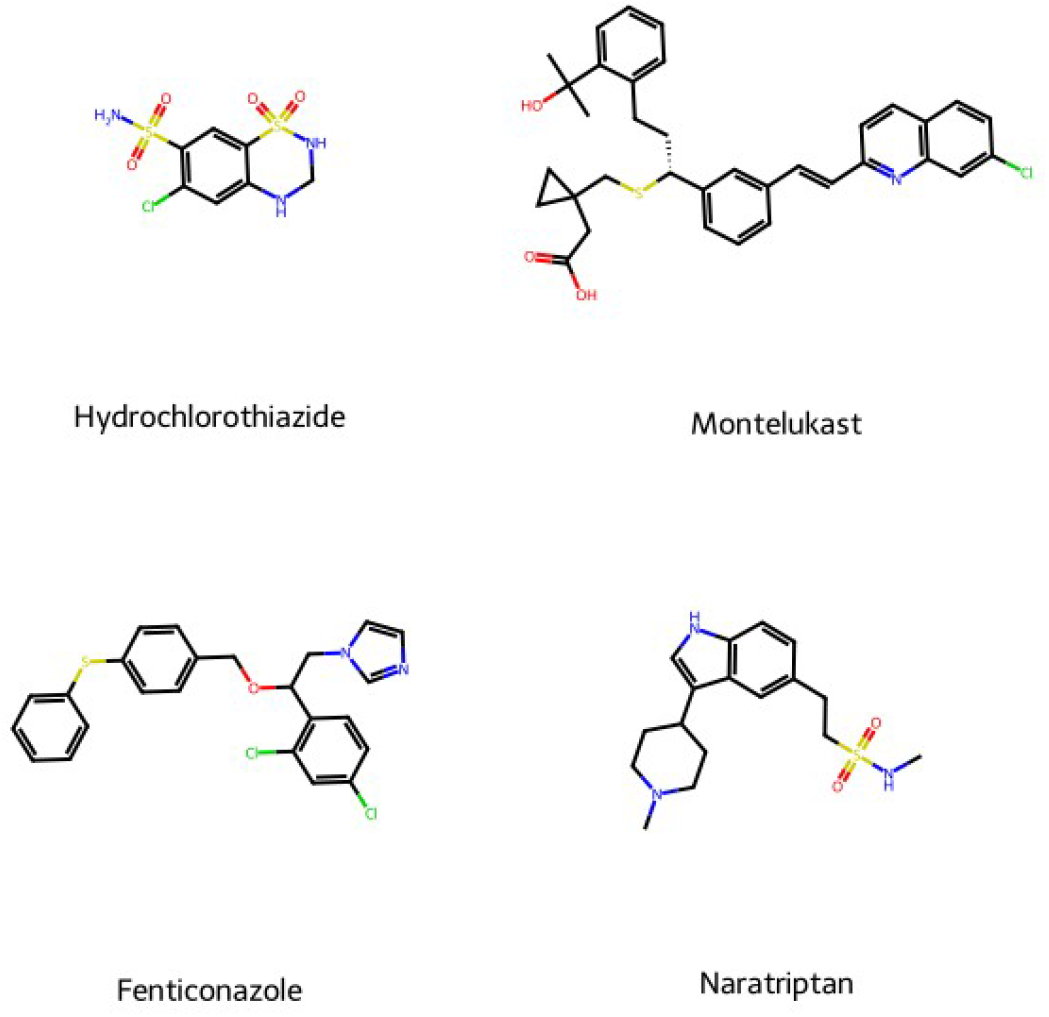
Chemical structures of the compounds submitted to experimental validation.

### 2.5 Competition Experiments

The four selected candidates were evaluated using the ANS displacement assay, with oleic acid as a positive control. As expected, increasing concentrations of oleic acid resulted in a reduction of ANS fluorescence (∼500 nm) in EgFABP1, EmFABP1, and EmFABP3, confirming ligand displacement (Supplementary **Figures S5–S7**). A similar decrease in fluorescence intensity was observed for hydrochlorothiazide, supporting the predictions of the ligand-based model ensemble (Supplementary **Figures S8–S10**).

The results (**Table 6**) showed that EmFABP3 exhibits a lower apparent dissociation constant (K_d,app_) for oleic acid compared to EgFABP1 and EmFABP1 isoforms. Conversely, EgFABP1 appears to display higher affinity for hydrochlorothiazide relative to EmFABP1 and EmFABP3. Supplementary **Figures S11** and **S12** present representative binding curves derived from the three independent replicates obtained for each isoform–compound pair.

**Table 6.** Parameters obtained for oleic acid and hydrochlorothiazide across the three isoforms in the fluorescence displacement assays. Values represent the mean of three independent replicates ± standard deviation. ^a,b^ Significant differences according to Tukey’s test (α = 0.05).

| COMPOUND | $K_{app}$ (μM) ( ± SD) | | |
| --- | --- | --- | --- |
|  | EgFABP1 | EmFABP1 | EmFABP3 |
| Oleic acid | 1.15 (0.43) | 1.71 (0.29) | 0.12 (0.06) |
| Hydrochlorothiazide | 2.19<sup>a</sup> (0.51) | 9.90<sup>a,b</sup> (4.33) | 2.75<sup>b</sup> (1.64) |

Montelukast, fenticonazole, and naratriptan were also evaluated using the ANS displacement assay; however, due to their specific spectroscopic properties, it was not possible to determine their respective K_d,app_ . Further details are provided in the Supplementary Information (**Figures S13-15**).

## 3 Conclusion

In this study, a parallel ligand-based and structure-based virtual screening approach was implemented using EgFABP1 from *E. granulosus* as the molecular target.

Regarding the ligand-based VS, we initially followed the strategy proposed by Bélgamo et al. [18]. In that study, the authors compiled compounds with reported experimental activity against AFABP under a target repositioning paradigm [19], generating a dataset of 258 compounds. In the present work, this dataset was expanded to 288 compounds, incorporating both experimentally validated ACTIVE and INACTIVE molecules recently reported in the literature. Furthermore, the decoy sets were enlarged from 1,500 to 7,613 and 4,965 compounds using LUDe and DUDE-Z, respectively.

A total of 4,000 linear classifiers were constructed and validated, and the best-performing models were subsequently combined into the MIN32 ensemble. The performance of MIN32 was comparable between the RL1 and RL2 libraries in terms of Sp, Se, and PPV, with a slight trend favoring RL2. This difference has been previously observed in other databases [20] and may be attributed to differences in decoy generation methods (LUDe vs. DUDE-Z), considering that the sets of true ACTIVE and INACTIVE compounds were identical in both simulated libraries.

The MIN32 ensemble was applied to screen eight chemical databases comprising approximately 435,000 compounds. From this pool, only 56 compounds were classified as hits, underscoring the high specificity of the model. Among these, hydrochlorothiazide, fenticonazole, and naratriptan were selected for experimental validation. Compounds with repurposing potential were prioritized, as they are more readily accessible and offer a streamlined path through preclinical and clinical development, given that their pharmacokinetic, pharmacodynamic, and safety profiles are already established [21,22].

In parallel, a structure-based VS campaign was conducted using the EgFABP1 crystal structure (PDB: 1O8V). Several docking programs were evaluated, with AD Bias (-D-) demonstrating superior screening performance based on retrospective validation. This finding supports the hypothesis that specific water molecules within the binding site stabilize the triad Arg107, Arg127, and Tyr129, thereby favoring the binding of certain ligand classes to cestode FABPs [17]. The theoretical PPV values associated with the selected cutoff indicate high specificity of the docking model, consistent with the relatively low number of predicted hits (98 compounds out of over 400,000). In terms of PPV, the docking protocol performed better for dRL2 than for dRL1, with more pronounced differences than those observed for the ligand-based models. From this structure-based screening, montelukast was selected for *in vitro* evaluation.

Recombinant EgFABP1, EmFABP1, and EmFABP3 were purified as previously described [10,18]. In the case of hydrochlorothiazide, the in silico predictions were experimentally confirmed, with estimated K_d,app_ values of 2.19, 9.90, and 2.75 µM for EgFABP1, EmFABP1, and EmFABP3, respectively. The observed differences in K_d,app_ among isoforms may reflect structural variations and, to some extent, differences in residual hydrophobic ligand content carried over during purification.

The remaining selected compounds exhibited interference in the fluorescence assays. Experimentally assessing FABP inhibition is inherently challenging, as lipid-binding activity cannot be quantified directly in the same manner as enzymatic reactions, where product formation can be readily measured. Due to these methodological constraints, K_d,app_ values could not be determined for three of the four in silico hits.

Hydrochlorothiazide is a thiazide diuretic with reported antiseizure and anticancer activities [23,24]. These characteristics make it a compelling example of a drug repurposing candidate—that is, a compound originally approved for one indication that may be repositioned for new therapeutic applications [21]. The experimentally confirmed affinity of hydrochlorothiazide for cestode FABPs also provides a starting point for *de novo* anthelmintic development through a hit-to-lead optimization program.

In conclusion, this work highlights the value of integrated in silico screening strategies for the discovery of new therapeutic candidates for neglected tropical diseases (NTDs). Computational approaches enable the identification of promising compounds in significantly shorter timeframes and at substantially lower cost compared to traditional experimental screening methodologies. Drug repurposing strategies, in particular, may help mitigate the limited private-sector investment in NTDs, which disproportionately affect low-resource populations and regions.

## 4 Experimental section

### 4.1 Materials

Naratriptan HCl and fenticonazole nitrate were acquired from Cayman Chemical (Ann Arbor, MI). Oleic acid was purchased from Sigma-Aldrich (St. Louis, MO), and 1-anilinonaphthalene-8-sulfonic acid (ANS) was purchased from Molecular Probes (Eugene, Oregon, USA). Hydrochlorothiazide and montelukast were kindly donated by *Laboratorios Bagó* (Buenos Aires, Argentina).

### 4.2 Recombinant Protein Expression

Selected hits were experimentally validated through fluorescence displacement assays against recombinant FABPs from *E. granulosus* (EgFABP1) and *E. multilocularis* (EmFABP1 and EmFABP3). Expression clones for all three proteins were obtained as previously described [10,18].

EmFABP1 and EmFABP3 were expressed in *Escherichia coli* BL21(DE3) cells using pET28a(+) vectors (Novagen, 69864), whereas EgFABP1 was expressed from a pET11b vector (Novagen, 69437). Protein expression was induced with isopropyl β-D-1-thiogalactopyranoside (IPTG; GenBioTech, I2481C5). Cells were lysed by sonication and centrifuged at 8,000 × g for 40 min at 4 °C.

EmFABP1 and EmFABP3 were initially purified by nickel-affinity chromatography, followed by size-exclusion chromatography using a Superdex 75 column (GE Healthcare, 17-5174-01). EgFABP1 was purified by Sephadex G-50 size-exclusion chromatography (Pharmacia Biotech Inc., 17-0042-02) [10]. All proteins underwent a delipidation step using Lipidex-100 resin (Sigma, H6383). Purity was assessed by 15% SDS-PAGE, and protein concentration was determined spectrophotometrically at 280 nm using the corresponding extinction coefficients: 9,530 M^⁻1^cm^⁻1^ (EgFABP1), 10,033 M^⁻1^ cm^⁻1^ (EmFABP1), and 14,105 M^⁻1^ cm^⁻1^ (EmFABP3) [18,25].

### 4.3 Competition Experiments

Fluorescence displacement assays were performed to evaluate the binding of selected hits to cestode FABPs, following the protocol described by Bélgamo et al. [18]. ANS was used as the fluorescent probe, as it binds to the FABP ligand-binding site and exhibits distinct emission spectra in its bound and unbound states. Ligand binding was inferred from changes in ANS fluorescence upon increasing concentrations of the test compounds.

Apparent dissociation constants (K_d,app_) for each protein–ligand pair were estimated by calculating the area under the fluorescence emission curve (420–600 nm) after Raman peak correction, and the resulting data were fitted according to Equation 2.

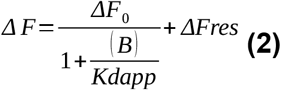

where ΔF represents the observed fluorescence at a given ligand concentration (B), ΔF_0_ corresponds to the amplitude of the fluorescence change, and ΔFres denotes the residual fluorescence at saturating ligand concentrations. Fluorescence data were normalized by dividing each value by ΔFmax (ΔF_0_ + ΔFres).

Assays were performed using 1 µM protein and 20 µM ANS in PBS (pH 7.4) at 25 °C, in a final volume of 200 µL. Ligands were added incrementally until no further changes in fluorescence were detected. Excitation was set at 350 nm, and emission spectra were recorded between 380 and 600 nm. Oleic acid was used as a positive control [18], and a 2-min incubation period was allowed between successive additions.

## 5 Computational methods

### 5.1 Dataset Compilation and Partition

Using a target repositioning approach, a previously reported dataset from Bélgamo et al.[18] was expanded by searching the scientific literature for compounds with reported activity against human adipocyte FABP (AFABP, also known as FABP4) [26–33]. Compounds were classified as ACTIVE or INACTIVE based on their half-maximal inhibitory concentration (IC_50_) or inhibition constant (K?). Compounds with IC_50_ or K_i_≤ 10 μM were considered ACTIVE, whereas those with values > 20 μM were classified as INACTIVE. Compounds with values between 10–20 μM were excluded to avoid ambiguity near the activity threshold. The final dataset comprised 288 compounds, including 187 ACTIVE and 101 INACTIVE compounds. A heatmap was generated to visualize the chemical diversity of the compiled dataset. Both the heatmap and the dataset are available in the Supplementary Information (**Figure S16** and **Table S3**, respectively).

The 101 INACTIVE compounds were included in the training set, while a representative subset of 101 ACTIVE compounds was selected using the iRaPCA subspace clustering method [34] to ensure class balance and chemical diversity within the training set (see **Supplementary Section E** for clustering details). The ACTIVE compounds included in the training set were sampled from each cluster proportionally to cluster size. The remaining 86 ACTIVE compounds were used to generate validation sets for retrospective screening experiments. These sets were enriched with synthetic decoys using the LUDe [20] and DUDE-Z [35] tools. As a result, two retrospective libraries (RL1 and RL2) were generated, comprising 7,699 and 5,051 compounds, respectively. **Table 7** summarizes the final composition of these libraries.

**Table 7.** Dataset composition.

|  | Training set | Validation sets |  |
| --- | --- | --- | --- |
|  |  | RL1 | RL2 |
| ACTIVE | 101 | 86 | 86 |
| INACTIVE | 101 | - | - |
| DECOYS | - | 7,613 | 4,965 |
| TOTAL | 202 | 7,699 | 5,051 |

### 5.2 Molecular Descriptor Calculation and Modeling Procedure

ACTIVE and INACTIVE compounds were retrieved using the ChEMBL web interface [36], and their SMILES representations were obtained and standardized using the MolVS library [37]. Decoy compounds were obtained in SMILES format from LUDe and DUDE-Z [20,35] and standardized using the same protocol. After standardization, 1,613 conformation-independent molecular descriptors were calculated using Mordred [14]. Descriptors with low variability (standard deviation < 0.05) were excluded. A dummy variable was assigned to label compounds as follows: ACTIVE = 1, whereas INACTIVE and decoys = 0.

A forward stepwise feature selection combined with a semi-correlation approach was applied to generate 4,000 linear classifiers, each built from random subsets of 200 descriptors. The number of descriptors per model was limited to a maximum of five to reduce the risk of overfitting. Models were validated using Leave-Group-Out (LGO) cross-validation [38] and Y-randomization [39], each performed over 1,000 iterations. In each LGO iteration, 10 ACTIVE and 10 INACTIVE compounds were excluded from the training set.

### 5.3 Ensemble Learning

To construct model ensembles, individual classifiers were first ranked according to the area under the Receiver Operating Characteristic curve (AUC-ROC) [40] obtained on the RL1 retrospective dataset. To avoid redundancy, only non-correlated models were retained; when two models shared more than one molecular descriptor, the model with the higher AUC-ROC was retained, and the other was discarded. Five ensemble combination strategies were evaluated to compute a consensus score: (i) MIN, the minimum score among ensemble members; (ii) PROD, the product of individual scores; (iii) AVE, the average of the individual scores; (iv) VOT, average voting as defined in [41]; and (v) RANK, the average of the rankings assigned by each individual model in the ensemble.

The RL1 dataset was split into two subsets of similar size: RL1-A (43 ACTIVE compounds and 3,807 decoys) and RL1-B (43 ACTIVE compounds and 3,806 decoys). RL1-A was used to optimize the number of models included in the ensemble and to select the most appropriate combination scheme. RL1-B and RL2 were subsequently used for independent validation of the selected ensemble. Early enrichment performance was quantified using the Boltzmann-enhanced discrimination of receiver operating characteristic (BEDROC) metric with α = 100 [42].

### 5.4 Score Threshold Selection and Prospective Ligand-Based VS

A critical aspect of virtual screening is estimating the probability that an *in silico* hit will confirm the predicted activity upon experimental validation. For this purpose, Positive Predictive Value (PPV) surfaces were constructed using the RL1-B and RL2 datasets. PPV values were calculated according to **Equation 3:**

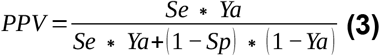

where Ya represents the proportion of ACTIVE compounds in the chemical library subjected to screening, whereas Se and Sp denote sensitivity and specificity, respectively. Ya cannot be known beforehand in a prospective VS experiment, as the true number of ACTIVE compounds is a priori unknown in real-world virtual screening applications. However, PPV surfaces can be estimated across the full range of possible score thresholds by varying Ya between 0.001 and 0.01, a range consistent with the proportion of ACTIVE compounds typically observed in high-throughput screening (HTS) campaigns [43,44].

After selecting an appropriate score cutoff, eight virtual chemical libraries were screened against cestode FABPs: DrugBank 5.1.6 [45], Drug Repurposing Hub (March 2020 release) [46], SWEETLEAD (November 2013 release) [47], HybridMolDB (September 2019 release) [48], NuBBEDB (November 2023 release) [49], NaturAr (November 2023 release) [50], COCONUT (January 2022 release) [51], and an in-house library comprising approximately 150 compounds. In total, approximately 435,000 molecular structures were evaluated. The applicability domain of the model was assessed for each predicted ACTIVE compound using the leverage approach [52].

### 5.5 Molecular Dynamics and WATCLUST Algorithm

A 100-ns of MD simulation was performed using the apo form of EgFABP1. The simulation was conducted with GROMACS 2020.4 employing the OPLS-AA force field [53,54]. The protein was protonated at pH 7.4, solvated in a dodecahedral box (205.13 nm^3^) containing 6,108 OPC water molecules [55], and neutralized with counterions.

Energy minimization was carried out using the steepest descent algorithm (step size = 0.01 nm; convergence criterion: maximum force < 1,000 kJ·mol^⁻1^·nm^⁻1^). System equilibration consisted of 100 ps in the NVT ensemble with positional restraints at 300 K using the modified Berendsen thermostat [56] (τ_t_ = 0.1 ps, time step = 4 fs), followed by 100 ps in the NPT ensemble under the same restraints using the Parrinello–Rahman barostat [57] (τ_p_ = 2 ps, compressibility γ = 4.5 × 10^⁻5^ bar^⁻1^, dt = 4 fs). A 100-ns production MD simulation was subsequently performed under NPT conditions without positional restraints.

Water sites (WS) within the EgFABP1 binding pocket were characterized using the WATCLUST clustering algorithm [58], which identifies regions exhibiting water occupancy higher than that of bulk solvent based on MD trajectories. WATCLUST, available as a VMD plugin [59], was employed to define WS that were subsequently incorporated as biasing potentials in the docking protocol using AutoDock-GPU. This strategy enables prioritization of interactions with specific water-mediated sites, thereby enhancing virtual screening performance [60].

### 5.6 Molecular Docking, Score Threshold Selection and Prospective VS Campaign

Four docking protocols were implemented: AutoDock-GPU (AD-GPU), AutoDock Bias (AD-Bias) [60,61], AutoDock Vina (Vina) [62], and Vinardo [63]. Docking parameters specific to each protocol are detailed in Supplementary **Table S4**. AD-Bias enables the incorporation of biasing potentials at predefined regions within the docking grid. Accordingly, the previously identified WS positions were used to define bias regions designed to favor hydrogen-bonding interactions mediated by water molecules.

The screening performance of each docking method was evaluated using the area under the receiver operating characteristic curve (AUC-ROC) [40]. For this purpose, the ACTIVE and INACTIVE compounds from the ligand-based model training set were merged with the RL1 and RL2 validation libraries, generating two new compound libraries comprising 7,901 and 5,253 molecules, respectively. These libraries are referred as dRL1 and dRL2, respectively.

The docking protocol demonstrating the best performance in both libraries was selected for the structure-based VS campaign. To determine an appropriate score threshold for structure-based VS, PPV surfaces were constructed for both dRL1 and dRL2, following the same procedure applied in the ligand-based retrospective experiments. Finally, the same chemical libraries previously screened in the ligand-based VS were subjected to structure-based VS.

### 5.7 Ligand and Protein Preparation for Docking

EgFABP1 and all ligands were protonated at pH 7.4. Ligands were energy-minimized using the steepest descent algorithm followed by the conjugate gradient algorithm, employing the default parameters implemented in OpenBabel 2.3 [64]. The minimized ligands and EgFABP1 were then converted to .pdbqt format using the prepare_ligand4.py and prepare_receptor4.py scripts, respectively [65].

A cubic grid box (22.5 Å per side) centered on the reported FABP binding site [17], with a grid spacing of 0.375 Å, was defined for both retrospective and prospective structure-based experiments. In the AD-Bias experiments, combinations of hydrogen-bond donor and acceptor biases were introduced using the prepare_bias.py script [65]. In all cases, a bias potential of ™2 kcal/mol and a decay ratio of 0.9 Å were applied. For all docking procedures, 20 independent runs per ligand were performed, and the top-scoring pose was used for performance evaluation.

## Author contributions

The study was conceived and designed by all the authors. SR, GFR, and AT performed material preparation, data collection, and analysis. SR, GFR, and AT drafted the initial version of the manuscript, and all authors contributed to its revision. All authors reviewed and approved the final version of the manuscript.

## Acknowledgements

All authors thank the Argentine National Council of Scientific and Technical Research (CONICET). This work was supported by CONICET, the National Agency for Scientific and Technological Promotion (ANPCYT) and the National University of La Plata (UNLP).

## Graphical abstract

*In silico* drug repurposing identified candidate compounds targeting cestode fatty acid binding proteins (FABPs). Hydrochlorothiazide bound multiple parasite FABPs *in vitro*, validating computational predictions and highlighting a translational approach to discover new antiparasitic leads against echinococcosis, a neglected disease with limited treatment options.

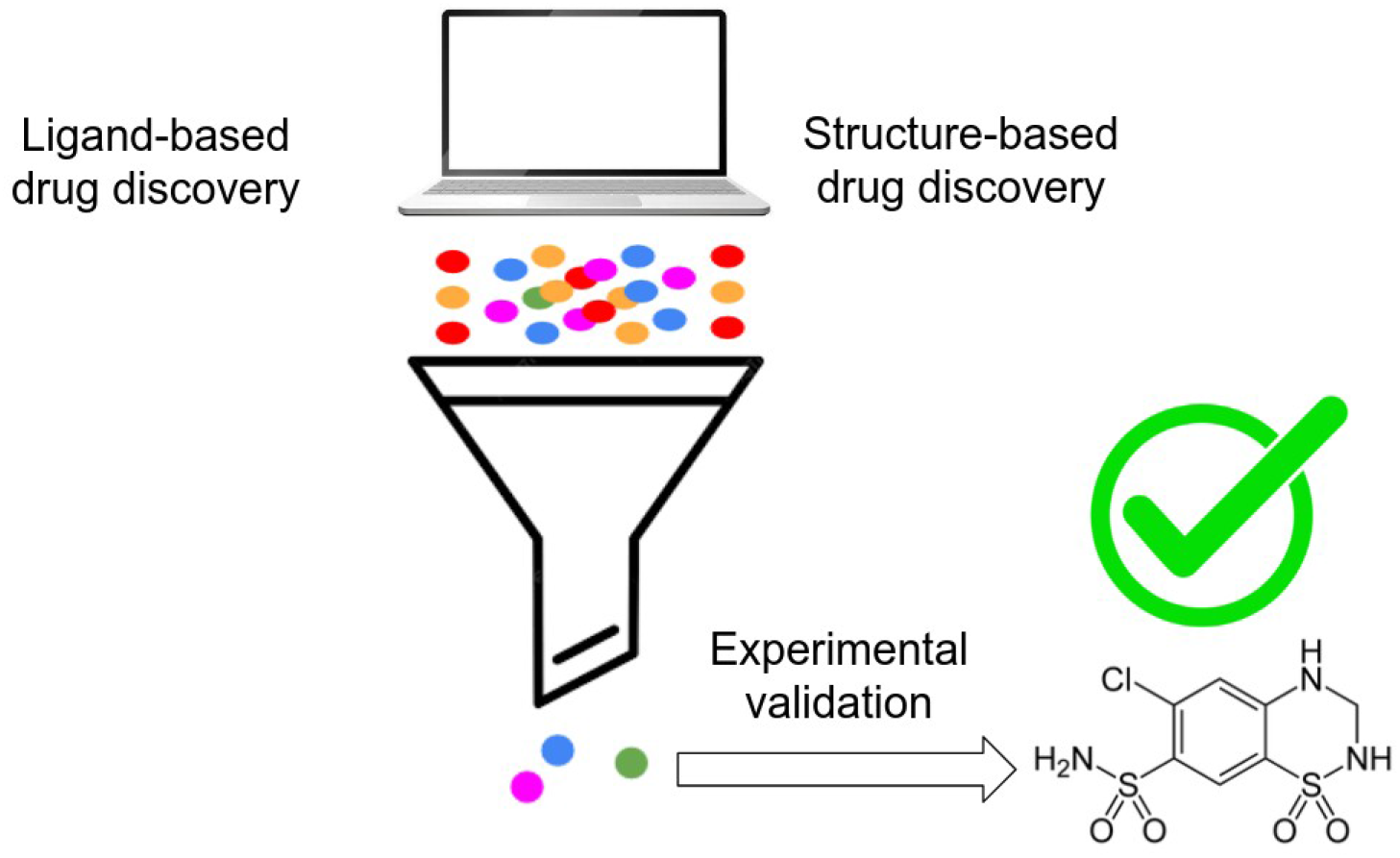

