## Supplementary material for "*In silico* drug repurposing and in vitro validation of cestode fatty acid binding proteins": Supp_material_rodriguez_et_al_2026.docx

**Supplementary section A: Ligand-Based VS results**

**List S1.** Mathematical equations for each of the models involved in the MIN32 ensemble.

**MODEL 3,963**

$$Y=0.3832*nAcid-0.0399*ATSC8are-0.0211*AATS6i$$

$-0.2626*AATSC8i+0.0014*ATSC3dv+3.5257$

**MODEL 1,940**

$$Y=0.3867*nAcid+0.2627*AATSC3dv-0.0835*AATSC8s$$

$$+0.1366*n5aHRing-0.0002*ATSC2m+0.2068$$

**MODEL 2,836**

$$Y=0.2017*AATS4d+0.4267*nAcid+0.1158*ATSC3pe$$

$$-0.0023*ATSC8s+0.2902*GATS8v-0.7271$$

**MODEL 3,392**

$$Y=0.5263*nAcid+0.1224*ATSC3are+0.2328*AATS2d$$

$$-0.4795*MATS8are-0.3280*GATS8dv-0.4254$$

**MODEL 3,555**

$$Y=0.5342*nAcid+0.0106*ATSC3s-0.3731*MATS8c$$

$$-0.0001*VR1_{A}-0.0084*AATS8i+1.5273$$

**MODEL 498**

$$Y=0.4899*nAcid-0.5219*ATSC8c+0.1612*ATSC3se$$

$$+0.2591*AATS1d-0.8769$$

**MODEL 3,520**

$$Y=0.4087*nAcid-0.3794*MATS8se-0.0313*SssO$$

$$+0.0196*nBondsA+0.2184*nG12FAHRing-0.0660$$

**MODEL 2,525**

$$Y=0.4302*nAcid+0.2725*AATS6d-0.0355*AATS8s$$

$$-0.0023*ATSC7s+0.5559*ATSC7c-0.4543$$

**MODEL 3,732**

$Y=0.4277*nAcid-0.0598*ATSC8se+0.4692*MWC02$

$$-0.2142*NssO+0.3363*GATS8p-2.3491$$

**MODEL 3,668**

$$Y=0.4084*nAcid-0.3812*MATS8s+0.1335*naHRing$$

$$+0.2821*IC5-0.0115*{PEOE}_{VSA9}-1.1934$$

**MODEL 2,808**

$$Y=0.3664*nAcid-0.7298*MATS8pe+0.4163*NaaS$$

$$-0.4189*GATS4s-0.2157*GATS8are+0.7976$$

**MODEL 2,892**

$$Y=0.6062*AATS4d+0.0384*C2SP3-0.2143*NssO$$

$$-0.2031*SdssC-0.1481*nFaRing-1.4184$$

**MODEL 527**

$$Y=0.2582*nAcid+0.0056*MPC4-0.0118*AATSC8m$$

$$+0.1545*ATSC1p-0.0247*{PEOE}_{VSA2}-0.0161$$

**MODEL 1,992**

$$Y=0.2523*nAcid+0.0058*AATS6v+0.2713*nG12FRing$$

$$-0.0288*{SlogP}_{VSA11}-0.1774*GATS5s-0.5156$$

**MODEL 2,346**

$$Y=0.1555*piPC9-0.2763*n9FHRing-0.0441*SssO$$

$$-0.3028*SdssC+0.0064*ATSC3s-0.6410$$

**MODEL 3,292**

$$Y=0.1756*piPC8-0.3382*GATS6are+0.2205*ATSC3se$$

$$-0.2331*SdssC-0.0009*{SpAD}_{Dzp}-0.1587$$

**MODEL 1,412**

$$Y=0.2095*nAcid+0.0004*TMPC10-0.0001*ATSC8m$$

$$-0.0106*{SlogP}_{VSA2}+0.2426$$

**MODEL 3,674**

$$Y=-0.1337*GATS6s-0.4315*MATS8se+0.0231*ATSC1i$$

$$+0.1451*ATSC3pe-0.1897*SdssC+0.6155$$

**MODEL 2,689**

$$Y=-0.0280*AATS6i-0.0498*AATSC8Z-0.2416*GATS5s$$

$$+0.1171*ATSC3are-0.1611*SdssC+5.1128$$

**MODEL 3,558**

$$Y=0.2178*nAcid-0.0682*AATSC8Z+0.0782*C3SP2$$

$$+0.2745*nG12FARing+0.2319*NaaNH+0.0608$$

**MODEL 3,896**

$$Y=-0.4483*GATS6are+0.2837*AATSC3dv-0.0377*AATS4i$$

$$-0.2303*NssO-0.2625*n9FRing+7.1551$$

**MODEL 2,933**

$$Y=0.0016*MPC10-0.0138*AATSC8v-0.0106*{EState}_{VSA2}$$

$$-0.3016*SdssC-0.5428*ATSC0c+0.6600$$

**MODEL 3,744**

$$Y=0.4281*AATS5d+0.4334*nG12FHRing-0.1883*SdssC$$

$$-0.2444*GATS5are-0.0159*{SlogP}_{VSA3}-0.5487$$

**MODEL 3,390**

$$Y=0.4446*GATS8m+0.0504*SaasC+0.3003*AATSC3dv$$

$$-0.3193*SdssC+0.1052*SaaNH-0.3248$$

**MODEL 769**

$$Y=0.0014*MPC8-0.0195*ATSC8p-0.0269*{PEOE}_{VSA2}$$

$$+0.0181*ATSC1i+0.2083*AATSC3dv+0.4909$$

**MODEL 2,424**

$$Y=-0.3069*GATS6s+0.3151*AATSC3dv+0.1220*AATSC1v$$

$$-0.0013*ATSC7dv-0.0869*SsNH2+0.6509$$

**MODEL 327**

$$Y=-0.0238*AATS6i-0.3807*GATS5se+0.2217*AATSC3dv$$

$$+0.0017*ATSC1v-0.0097*AATSC8m+4.5510$$

**MODEL 2,269**

$$Y=0.0743*C3SP2-0.0005*ATSC2v+0.7346*AATS3d$$

$$+0.6063*CIC5-0.0910*nHBAcc-1.9953$$

**MODEL 3,976**

$$Y=-0.6048*ATSC8c-0.0125*ATSC7d+0.0019*ATSC1v$$

$$-0.2355*GATS5s-0.0113*AATSC6v+0.5113$$

**MODEL 3,380**

$$Y=0.3658*AATS2d-0.5395*GATS4s-0.0320*{PEOE}_{VSA12}$$

$$-0.0752*ATSC8se-0.0106*AATSC8v-0.3636$$

**MODEL 1,358**

$$Y=-0.0001*ATSC8m-0.0264*AATS5i+0.3326*AATSC3dv$$

$$-0.0721*ATSC4are-0.3909*GATS4s+5.0729$$

**MODEL 1,178**

$$Y=0.5866*AATS4d-0.0104*{EState}_{VSA2}+0.0798*AATSC1v$$

$$-0.0314*{ETA}_{dBeta}-0.0072*ATSC8d-1.2462$$

**Table S1.** Internal validation results for the 32 models comprising the MIN32 ensemble. The accuracy score represents the mean for 1,000 rounds of randomization for each case.

| **MODEL NUMBER** | **TRAINING SET** | **Y-RANDOMIZATION** | | **LGO** | |
| --- | --- | --- | --- | --- | --- |
|  | **ACCURACY**  **(%)** | **ACCURACY**  **(%)** | **SD**  **(%)** | **ACCURACY**  **(%)** | **SD**  **(%)** |
| 3963 | 79.70 | 49.38 | 11.30 | 71.87 | 9.40 |
| 1940 | 80.20 | 50.00 | 12.05 | 73.02 | 9.24 |
| 2836 | 77.72 | 50.00 | 12.50 | 72.53 | 9.33 |
| 3392 | 80.20 | 50.00 | 13.15 | 73.88 | 9.42 |
| 3555 | 81.19 | 50.00 | 13.46 | 73.57 | 9.78 |
| 498 | 78.71 | 50.00 | 12.70 | 75.04 | 9.41 |
| 3520 | 80.69 | 50.00 | 13.33 | 74.53 | 8.92 |
| 2525 | 74.26 | 50.00 | 11.74 | 69.95 | 9.21 |
| 3732 | 76.73 | 50.00 | 11.90 | 72.30 | 9.42 |
| 3668 | 76.73 | 50.00 | 13.07 | 75.14 | 9.08 |
| 2808 | 77.23 | 50.00 | 12.45 | 71.03 | 9.10 |
| 2892 | 74.75 | 50.00 | 12.19 | 70.44 | 9.40 |
| 527 | 75.25 | 50.00 | 12.06 | 70.33 | 9.55 |
| 1992 | 78.22 | 50.00 | 13.83 | 75.40 | 9.22 |
| 2346 | 72.77 | 50.00 | 11.55 | 68.65 | 9.74 |
| 3292 | 75.74 | 49.00 | 14.39 | 71.09 | 9.25 |
| 1412 | 77.72 | 49.00 | 11.83 | 74.05 | 9.02 |
| 3674 | 74.75 | 50.00 | 12.08 | 70.80 | 10.20 |
| 2689 | 76.73 | 50.00 | 11.59 | 69.59 | 9.87 |
| 3558 | 77.23 | 49.00 | 13.05 | 74.11 | 8.93 |
| 3896 | 74.75 | 49.00 | 11.00 | 66.44 | 9.57 |
| 2933 | 73.27 | 49.00 | 11.73 | 67.10 | 9.85 |
| 3744 | 76.73 | 49.00 | 11.51 | 69.85 | 10.19 |
| 3390 | 73.76 | 50.00 | 11.86 | 69.81 | 9.56 |
| 769 | 74.26 | 50.00 | 11.71 | 70.00 | 9.27 |
| 2424 | 74.26 | 50.00 | 12.75 | 69.62 | 9.72 |
| 327 | 73.76 | 50.00 | 11.86 | 69.81 | 9.56 |
| 2269 | 76.73 | 50.00 | 12.31 | 70.94 | 9.64 |
| 3976 | 77.23 | 50.00 | 11.86 | 70.65 | 9.85 |
| 3380 | 76.24 | 49.00 | 11.99 | 71.47 | 9.70 |
| 1358 | 76.73 | 49.00 | 12.08 | 72.24 | 9.36 |
| 1178 | 72.77 | 50.00 | 14.06 | 70.61 | 9.73 |

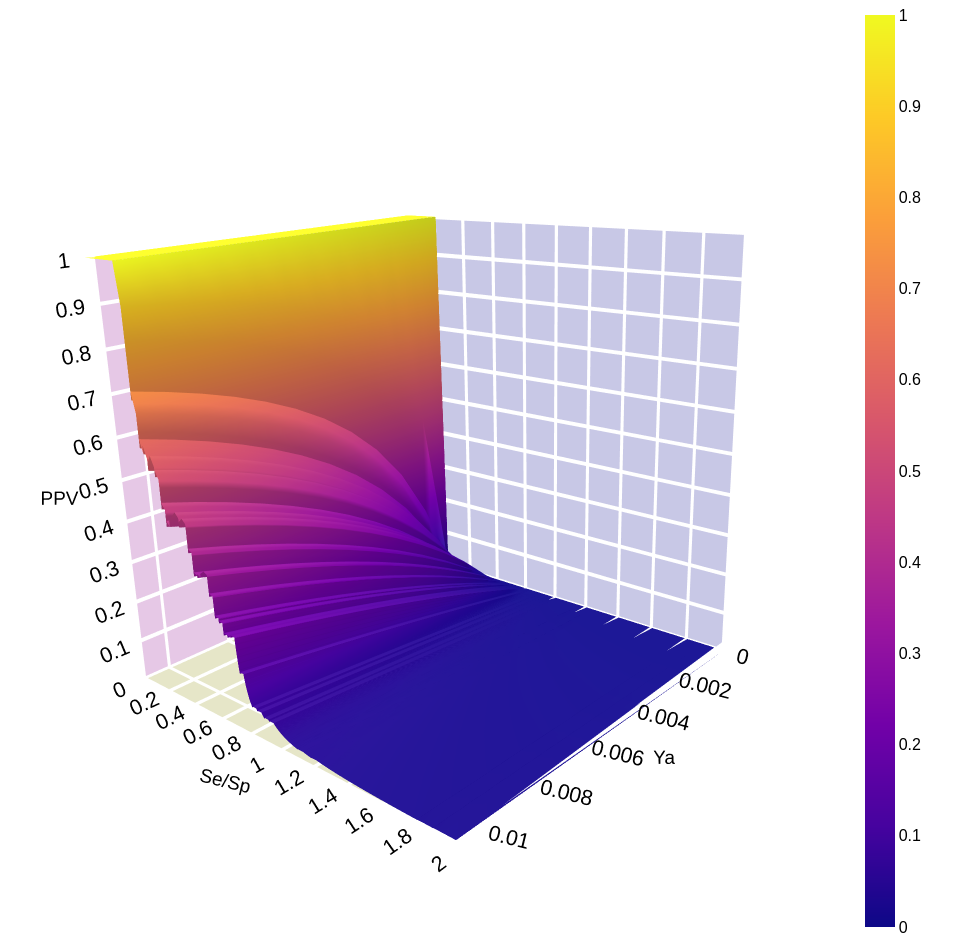
**Figure S1.** PPV surface for the MIN32 ensemble using the RL1-B library.

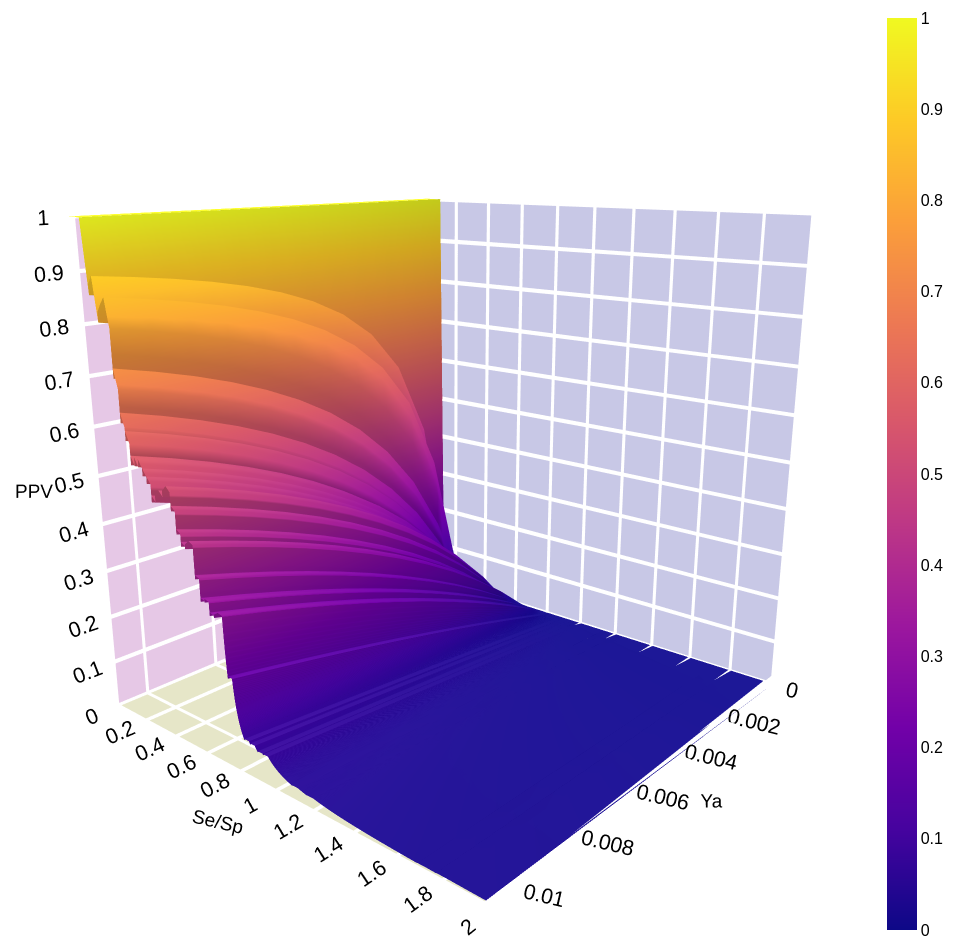
**Figure S2.** PPV surface for the MIN32 ensemble using the RL2 library.

**List S2.** SMILES code for the hit compounds found in prospective ligand-based VS.

CNS(=O)(=O)CCc1ccc2[nH]cc(C3CCN(C)CC3)c2c1

O=C1N(Cc2ccccc2)C(COc2ccccc2)C(O)C(O)C(COc2ccccc2)N1Cc1ccccc1

CC(C)COc1cccc(-c2nc3cc(C(=N)N)c(Cl)cc3[nH]2)c1O

Clc1ccc(CN(Cc2ccc(Cl)cc2)c2nn[nH]n2)cc1

O=C(O)c1cccnc1Nc1cccc(C(F)(F)F)c1

CC(C)(C)c1nc2c3cc[nH]c(=O)c3c3cc(F)ccc3c2[nH]1

CC1(C)Cc2c(-c3ccccc3)c(-c3ccc(Cl)cc3)c(CC(=O)O)n2C1

CCCCC(NC(=O)C(Cc1ccccc1)NC(=O)C(N)Cc1ccccc1)C(=O)NC(CCCNC(=N)N)C(=O)NCc1ccncc1

CC(NC(=O)C(=O)Nc1ccccc1C(C)(C)C)C(=O)NC(CC(=O)O)C(=O)COc1c(F)c(F)cc(F)c1F

COc1cc2c(cc1OC)C(Cc1cc(OC)c(OC)c(OC)c1)[N+](C)(CCCOC(=O)C(Cl)=CC(=O)OCCC[N+]1(C)CCc3cc(OC)c(OC)cc3C1c1cc(OC)c(OC)c(OC)c1)CC2

COc1ccc(-c2sc3cc(O)ccc3c2Oc2ccc(OCCN3CCCCC3)cc2)cc1

CC(=O)N1CCC(NC(=O)NC23CC4CC(CC(C4)C2)C3)CC1

Fc1ccc(-c2cncc(CNCC3CCc4ccccc4O3)c2)cc1

N=C(N)c1ccc(-c2ccc(OCC3CC(CC(=O)O)C(=O)N3)cc2)cc1

CCOC(Cc1ccc(OCCN2c3ccccc3Oc3ccccc32)cc1)C(=O)O

CCc1nc(N)nc(N)c1-c1ccc2c(c1)N(CCNC(C)=O)C(=O)C(C)(C)S2

NCc1ccc2c(c1)C1(CCN(C(=O)c3ccc(C#Cc4ccccc4)o3)CC1)CO2

NC(COc1cncc(C2=NCC3=NN=CC3=C2)c1)CC1=C2C=CCC=C2N=C1

NS(=O)(=O)c1ccc(NC(=O)c2ccccc2S)cc1

CC(C)C(CC(O)C(N)CC1CCCCC1)C(=O)O

COC(=O)CC(CCc1ccc(C(=N)N)cc1)c1cccc(C(=N)N)c1

COc1cc(N=Nc2ccccc2C(=O)O)cc(OC)c1O

CC(Cc1ccccc1)NN

CCC(=O)OCCNC(=O)C[n+]1ccccc1

CNC(=O)c1ccc(N2C(=S)N(c3cnc(C#N)c(C(F)(F)F)c3)C(=O)C23CCC3)cc1F

CCc1cnn2c(NCc3ccc[n+]([O-])c3)cc(N3CCCCC3CCO)nc12

CN1C(=O)N(c2ccc(C#N)c(C(F)(F)F)c2)C(=O)C1(CO)c1ccccc1

COc1ccc(C(=O)c2c(C)n(CCN3CCOCC3)c3cc(I)ccc23)cc1

CC1(C)CCC2=C(O1)c1ccccc1C(=O)C2=O

O=[N+]([O-])c1cnc(Sc2nnc(O)n2-c2ccc3c(c2)OCCO3)s1

CN(C)C(=O)COC(=O)Cc1ccc(OC(=O)c2ccc(NC(=N)N)cc2)cc1

CNCCCOc1cc(F)c(-c2c(Cl)nc3ncnn3c2NC(C)C(F)(F)F)c(F)c1

Nc1ccc(F)cc1NC(=O)c1ccc(CNC(=O)C=Cc2cccnc2)cc1

NS(=O)(=O)c1cc2c(cc1Cl)N=CNS2(=O)=O

Cc1ncsc1CCCl

c1ccc(-c2cnn3cc(-c4ccc(OCCN5CCOCC5)cc4)cnc23)cc1

CCOc1ccc(C=C2NCCc3cc(OCC)c(OCC)cc32)cc1OCC

Clc1ccc(C(Cn2ccnc2)OCc2ccc(Sc3ccccc3)cc2)c(Cl)c1

Oc1noc2c1CCNC2

O=C(NC1CC1)c1cc(-c2cccs2)on1

CCCCCOc1ccc(-c2cc(-c3ccc(C(=O)NC4CC(O)C(O)NC(=O)C5C(O)C(C)CN5C(=O)C(C(O)CC(N)=O)NC(=O)C(C(O)C(O)c5ccc(O)c(OS(=O)(=O)O)c5)NC(=O)C5CC(O)CN5C(=O)C(C(C)O)NC4=O)cc3)no2)cc1

CN1C(=O)NC(Cc2c[nH]c3c(Cl)cccc23)C1=O

COc1ccccc1S(=O)(=O)Nc1ccc2c(c1)CN(C)C(=O)N2

C=CC[N+]1(C2CC3C4CCC5CC(O)C(N6CCOCC6)CC5(C)C4CCC3(C)C2OC(C)=O)CCCC1

CN(C)c1cccc(Oc2cnc(Nc3cccc(O)c3)nc2)c1

Cc1[nH]c(C=C2C(=O)Nc3cc(-c4ccccc4)ccc32)c(C)c1CCC(=O)O

OCC(O)C(O)C(O)C(O)C(O)C(O)C(O)CO

CN(C)CCc1c[nH]c2cccc(OP(=O)(O)O)c12

CCCCNc1ccc(C(=O)OCCN(CC)CC)cc1

CC(Cc1ccc(O)cc1)NCC(O)c1cc(O)cc(O)c1

CC[N+]1(CC)CCC(=C(c2ccccc2)c2ccccc2)C1C

COc1cccc(N(C)C(=S)Oc2ccc3c(c2)CCCC3)n1

N#CC(=C1SCC(c2ccc(Cl)cc2Cl)S1)n1ccnc1

COC1=C(OC)C23CCc4ccc(O)c(OC)c4C2(CCN3C)CC1=O

NS(=O)(=O)c1cc2c(cc1Cl)NC(Cc1ccccc1)NS2(=O)=O

Cn1cc(NC(=O)c2cc(NC(=O)c3cc(C(=O)N4CC(CCl)c5c4cc(O)c4c5cnn4Cc4ccccc4)nn3C)cn2C)cc1C(=O)NCCC(=N)N

**Supplementary section B: Structured-Based VS results**

**Table S2.** AUCROC values for the dRL1 and dRL2 libraries for each program. ^*;^**^†^**Significant differences between AD Bias (DDD) and the different docking algorithms in the dRL1 and dRL2 libraries, respectively (Tukey's Test (α=0.05)).

| **ALGORITHM** | **AUCROC**  **dRL1**  **(+/- SD)** | **AUCROC**  **dRL2**  **(+/- SD)** |
| --- | --- | --- |
| **Vina** | 0.6624^*^  (+/- 0.0095) | 0.8016**^†^**  (+/- 0.0070) |
| **Vinardo** | 0.6923^*^  (+/- 0.0074) | 0.7736**^†^**  (+/- 0.0061) |
| **AD GPU** | 0.6759^*^  (+/- 0.0087) | 0.8265**^†^**  (+/- 0.0068) |
| **AD Bias (AAA)** | 0.7054^*^  (+/- 0.0096) | 0.8402**^†^**  (+/- 0.0068) |
| **AD Bias (DDD)** | 0.7670  (+/- 0.0080) | 0.9030  (+/- 0.0059) |
| **AD Bias (AAD)** | 0.7218^*^  (+/- 0.0085) | 0.8547**^†^**  (+/- 0.0063) |
| **AD Bias (ADA)** | 0.7497^*^  (+/- 0.0084) | 0.8804**^†^**  (+/- 0.0065) |
| **AD Bias (DAA)** | 0.7368^*^  (+/- 0.0093) | 0.8592**^†^**  (+/- 0.0066) |
| **AD Bias (ADD)** | 0.7513^*^  (+/- 0.0084) | 0.8838**^†^**  (+/- 0.0065) |
| **AD Bias (DAD)** | 0.7334^*^  (+/- 0.0089) | 0.8632**^†^**  (+/- 0.0064) |
| **AD Bias (DDA)** | 0.7676  (+/- 0.0083) | 0.8958**^†^**  (+/- 0.0062) |

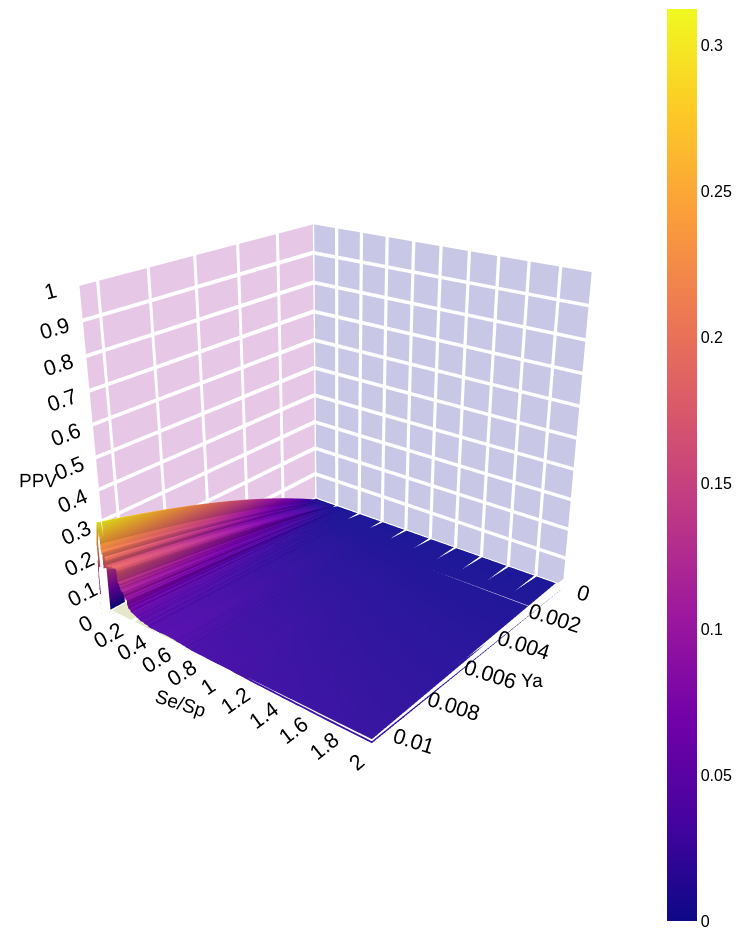

**Figure S3.** PPV surface for the -D- docking model using the dRL1 library.

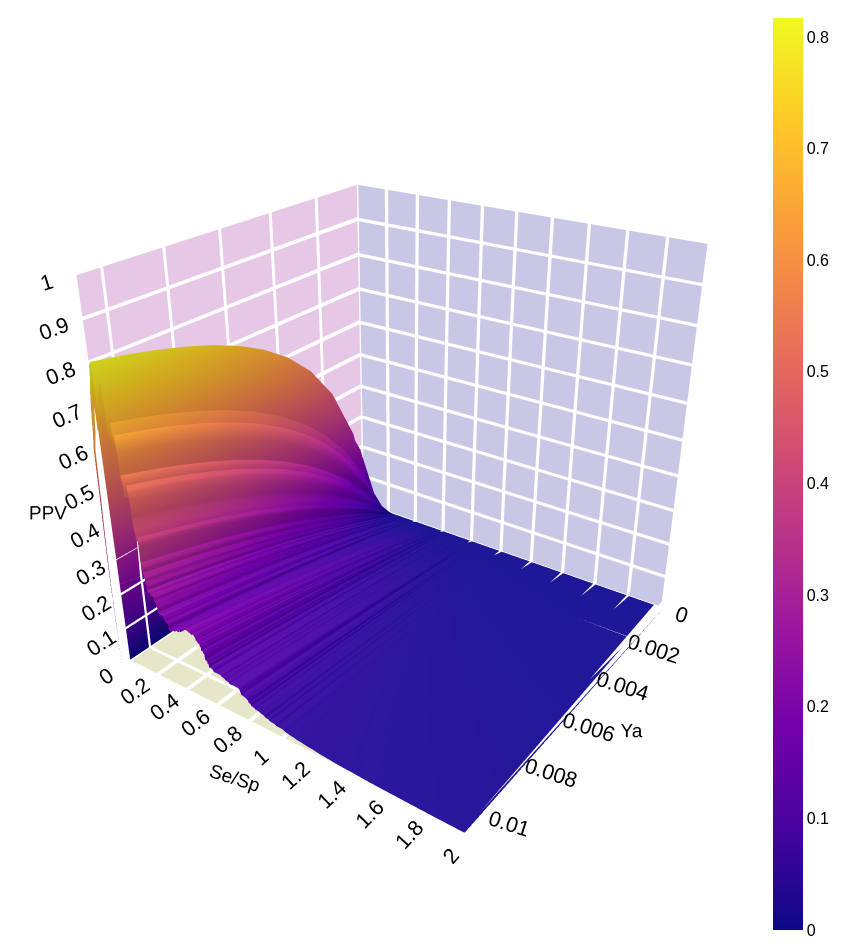

**Figure S4.** PPV surface for the -D- docking model using the dRL2 library.

**List S3.** SMILES code for the hit compounds found in structure-based VS.

O=C(O)CCC(=O)N1NC(c2c(-c3ccccc3)c3c4c(ccc3[nH]c2=O)C4)CC1c1ccc(Br)cc1

COC1=NC2(CO2)N=CC1c1ccc2c(NC3=CC4(C(=O)O)CC4(Oc4cc(F)cc(F)c4)C3)c(S(=O)(=O)NC3CC3)cnc2c1

CCOc1ccc(-c2ccc(Cn3c(CC(C)(C)C(=O)O)c([SH]456C7C48C5C786)c4cc(OCc5ccc(C)cn5)ccc43)cc2)cn1

CC(C)=CCC=C1CC1CCC(C)=CC=CC(C)CCCC=C1CC1CC=C(C)C=CC=C(C)CCC=C(C)C

CC(C)(C)Sc1c(CC(C)(C)C(=O)O)n(Cc2ccc(Cl)cc2)c2ccc(OCc3ccc4ccccc4n3)cc12

O=C(O)CCC1=C(OCc2ccc3c(O)noc3c2)C2C3C(O)(c4ccc(OC5CCCC5)cc4O)C23C1

O=C(O)CC1CCN(C2(O)C3OCCN(C(=O)Cc4ccc(NC(=O)Nc5cccc6c5C6)cc4)C32)CC1

CC1=C(C=CC2CC2CCC2CC2CCCC2CC2CC2CC2C=CC2=C(C)C(=O)C(O)CC2(C)C)C(C)(C)CC(O)C1=O

CC1=C2CC23C(=O)C(C2CCCC2)CC3C=C1OCc1cccc(-c2ccc(C(=O)O)cc2)c1

CN(C(=O)c1cc(OS(O)(O)c2ccc3ccccc3c2)ccc1NS(=O)(=O)c1ccc2ccccc2c1)C(CC(=O)O)C(=O)O

O=C(O)C(Cc1ccccc1)N(Cc1ccc(-c2cc3ccccc3o2)s1)C(=O)c1ccc(Cl)cc1Cl

CC(C)CCN(CC(O)C1Cc2ccc(cc2)OCCCC(=O)NC(CC(N)=O)C(=O)N1)S(=O)(=O)c1ccccc1

C=C1C2CCC3C(CCC4C(C(=O)O)(C(=O)O)CC(OC5OC(COO)C(OS(O)(O)O)C(OS(O)(O)O)C5OC(=O)CC(C)C)C5CC543)(C2)C1O

NC1=CC2C(=c3ccc(N)cc3=C(c3ccccc3)N2CCCCCCc2cnnn2CCNc2c3c(nc4ccccc24)CCCC3)C=C1

CN(C(=O)c1cc2ccccc2cc1C(=O)C(c1cccc2ccccc12)P(=O)(O)O)C1CCN(C(=O)c2ccc3ccccc3c2)CC1

N=C(N)c1cccc(CC(NS(=O)(=O)c2ccc3ccccc3c2)C(=O)N2CCCCC2C(=O)O)c1

CCC1=C(C)C(=O)NC1=CC1=NC(=Cc2[nH]c(C=C3NC(=O)C(C)=C3CC)c(CCC(=O)O)c2C)C(C)=C1CCC(=O)O

Cc1cc(-c2c3c(c(Br)c4ccccc24)SC(C)C3C)cc(C)c1OC(Cc1ccccc1)C(=O)O

CC1=CCC(C(=O)N(C2CCC(O)(COC3CCOC3)CC2)C2CC(CC34CC3(C)C4)SC2C(=O)O)CC1

CNC(=O)c1cc(Oc2ccc(NC(=O)NC3CC(C(C)(C)C)NN3c3ccc4ncccc4c3)c(F)c2)ccn1

COc1ccc(NS(=O)(=O)c2ccc(NC(C3C(=O)N(c4ccccc4)N(C)C3C)S(O)(O)O)cc2)nn1

CC1CC2C3CCC(O)(C(=O)COC(=O)CCC(=O)O)C3(C)CC(O)C23CC32CCC(=O)CC12

O=C(O)CN1C(=O)C(NC(=O)C2(CC(CCc3cccc4ccccc34)C(=O)O)CCCC2)CCc2ccccc21

CN(Cc1ccccc1)Cc1ccc(CC2Cc3ccc(OCCCCCN4CC5C6C4C56n4c(=O)[nH]c5ccccc54)cc3C2=O)cc1

C=CN(CC)c1ccc2cc(C(=O)c3ccc(N4CCN(C(=O)COC56CC5C(Cl)C=C6Cl)CC4)cc3)oc2c1

CC1CCC2C(C)C(CCS(=O)(=O)CCCS(=O)(=O)CCC3OC4OC5(C)CCC6C(C)CCC(C3C)C46OO5)OC3OC4(C)CCC1C32OO4

COP(=O)(OCCC1OC2OC3(C)CCC4C(C)CCC(C1C)C24OO3)OCCC1OC2OC3(C)CCC4C(C)CCC(C1C)C24OO3

Cc1ccc(Nc2c3ccccc3nc3ccccc23)cc1NC(=O)Oc1ccc(N(CCCl)CCCl)cc1

Cc1ccc(NC2=C3C=CC=C4CC43Nc3ccccc32)cc1NC(=O)Oc1ccc(N(CCCl)CCCl)cc1

CC1CCC2C(C)C(OCc3cc(O)cc(OC4OC5OC6(C)CCC7C(C)CCC(C4C)C57OO6)c3)OC3OC4(C)CCC1C32OO4

CC1CCC2C(C)C(OCC(COC3OC4OC5(C)CCC67CC6CCC(C3C)C47OO5)OS(C)(O)O)OC3OC4(C)CCC1C32OO4

Cc1nc2ccccc2c2ccc(NC(=O)NCCNC(=O)Nc3ccc4c(c3)=C3CN3C3C=CC=CC=43)cc12

O=C(NCCCCCCNC(=O)Nc1ccc2c(c1)=C1CN1C1C=CC=CC=21)Nc1ccc2c(c1)=C1CN1C1=CC=CCC=21

COc1cc2c(cc1OCC1CN(CCCCCOc3ccc4c(c3)CCC3C45CCC4(C)C(O)CCC345)NN1)N=CC1CCCN1C2=O

COc1cc2c(cc1OCc1cn(CCCCCOc3ccc4c(c3)CCC3C45CCC4(C)C(O)CCC345)nn1)N=CC1CC(F)(F)CN1C2=O

O=C(CCC(=O)Nc1ccc2c(c1)C13CCCCC1C(C2)N(CC1CCC1)CC3)Nc1ccc2c(c1)C13CCCCC1C(C2)N(CC1CCC1)CC3

COc1ccccc1-c1[nH]c2ccccc2c1C#CC1(O)CCC2C3CCc4cc(O)ccc4C3CCC21C

O=C(C1CC1)N(CCCCNC1=C2C=CC=CC2N=C2CCCCCCN21)CCCNC1=C2C=CC=CC2N=C2CCCCCCN21

Nc1nc2ccccc2c2c1NC(CCCCCCc1nc3c(N)nc4ccccc4c3n1Cc1ccccc1)N2Cc1ccccc1

CCC1(OC(=O)C(Cc2ccccc2)NC(=O)C2CC(C)(C)N(O)C2(C)C)C(=O)OCC2=C1Cc1c3nc4ccccc4cc3cn1C2=O

CCC1(OC(=O)C(CCSC)NC(=O)C2CC(C)(C)N(O)C2(C)C)C(=O)OCC2C(=O)n3cc4cc5ccccc5nc4c3C=C21

C#CCON=C(CC1OC2OC3(C)CCC4C(C)CCC(C1C)C24OO3)C1OC2OC3(C)CCC4C(C)CCC(C1C)C24OO3

C=CCON=C(CC1OC2OC3(C)CCC4C(C)CCC(C1C)C24OO3)C1OC2OC3(C)CCC4C(C)CCC(C1C)C24OO3

O=C1Nc2ccccc2C1=Cc1cn(CCCCOc2cc(O)c3c(=O)cc(-c4ccccc4)oc3c2)c2ccccc12

O=S(=O)(Cc1nnc(CS(=O)(=O)C2CNNC2S(=O)(=O)c2ccc(Cl)cc2)s1)Nc1ccc(Cl)cc1

COC1=C(OC)C2(C=C(C3c4cc5c(cc4C(NC(=O)CCCCOc4ccc6nc(C)n(C7C=CC8(C=C7)CO8)c(=O)c6c4)C4COC(=O)C34)OCO5)C1)CO2

O=C1c2ccccc2-c2c1c1ccccc1c(=O)n2CCCNCCn1c2c(c3ccccc3c1=O)C(=O)c1ccccc1-2

Cc1ccc(-n2nnnc2SCCCCCCCOc2ccc3c(c2)CCC2C34CCC3(C)C(=O)CCC234)cc1

CC12CCC3c4ccc(OCCCCCCCSc5nnnn5-c5ccccc5)cc4CCC3C1CCC2=O

C1=C(c2ccccc2)C23CC2CCC3=NC1c1ccc(Oc2c(-c3cc4c5c(c3)C45)nc3ccccc3c2-c2ccccc2)cc1

Brc1ccc(-c2nc3ccccc3c(-c3ccccc3)c2Oc2ccc(C3=CC(c4ccccc4)C45CC4C=CC5=N3)cc2)cc1

COc1ccc(-c2nc3ccccc3c(-c3ccccc3)c2Oc2ccc(C3C=C(c4ccccc4)C45CC4C=CC5=N3)cc2)cc1

C1=CC2(c3c(Oc4ccc(-c5cc(-c6ccccc6)c6ccccc6n5)cc4)c(-c4ccc5ccccc5c4)nc4ccccc34)CC2C1

Clc1cccc(-c2nc3ccccc3c(-c3ccccc3)c2Oc2ccc(C3C=C(c4ccccc4)C45CC4C=CC5=N3)cc2)c1

Brc1cccc(-c2nc3ccccc3c(-c3ccccc3)c2Oc2ccc(C3C=C(c4ccccc4)C45CC4C=CC5=N3)cc2)c1

COc1cccc(-c2nc3ccccc3c(-c3ccccc3)c2Oc2ccc(C3C=C(c4ccccc4)C45CC4C=CC5=N3)cc2)c1

Clc1ccc2nc(-c3ccc(Oc4c(-c5ccccc5)nc5ccc(Cl)cc5c4C45C=CCC4C5)cc3)cc(-c3ccccc3)c2c1

Clc1ccc(-c2nc3ccc(Cl)cc3c(-c3ccccc3)c2Oc2ccc(C3C=C(c4ccccc4)C45CC4(Cl)C=CC5=N3)cc2)cc1

O=C(CN1CCCCC1)Nc1cc(COC(=O)CCC2=CC(=O)c3ccccc3C2=O)cc(Nc2ccnc3ccc(Cl)cc23)c1

CC1CCC2C(C)C(CC(OC(=O)Nc3cccc(F)c3)C3OC4OC5(C)CCC6C(C)CCC(C3C)C46OO5)OC3OC4(C)CCC1C32OO4

CC1CCC2C(C)C(CC(OC(=O)Nc3ccc(F)c(F)c3)C3OC4OC5(C)CCC6C(C)CCC(C3C)C46OO5)OC3OC4(C)CCC1C32OO4

CC1CCC2C(C)C(CC(OC(=O)Nc3ccc(F)c(Cl)c3)C3OC4OC5(C)CCC6C(C)CCC(C3C)C46OO5)OC3OC4(C)CCC1C32OO4

CC1CCC2C(C)C(CC(OC(=O)Nc3cccc(Cl)c3)C3OC4OC5(C)CCC6C(C)CCC(C3C)C46OO5)OC3OC4(C)CCC1C32OO4

CC1CCC2C(C)C(CC(OC(=O)Nc3ccc(Br)cc3)C3OC4OC5(C)CCC6C(C)CCC(C3C)C46OO5)OC3OC4(C)CCC1C32OO4

CC1CCC2C(C)C(CC(OC(=O)Nc3ccc([N+](=O)[O-])cc3)C3OC4OC5(C)CCC6C(C)CCC(C3C)C46OO5)OC3OC4(C)CCC1C32OO4

CC1CCC2C(C)C(C3C(O)C3CC3OC4OC5(C)CCC67C8C(CC(C3C)C46OO5)C87)OC3OC4(C)CCC1C32OO4

COc1cc(CCC(=O)C=Cc2cccc3ccccc23)ccc1OCc1cn(CCN2C(=O)C(=O)c3cc(Br)ccc32)nn1

COc1cc(CCC(=O)C=Cc2cccc(Br)c2)ccc1OCc1cn(CCN2C(=O)C(=O)c3cc(Br)ccc32)nn1

COc1cc(OC)c2c(=O)cc(-c3ccc(OCc4cn(-c5ccccc5Cl)nn4)c(OCc4cn(-c5ccccc5Cl)nn4)c3)oc2c1

COC1=CC2(C=C3OC(c4ccc(OCc5cn(-c6ccc(Cl)cc6)nn5)c(OCc5cn(-c6ccc(Cl)cc6)nn5)c4)=CC(=O)C13)CO2

O=[N+]([O-])c1ccc(Cl)c(-c2nccc3c4ccccc4n(CCCCCCn4c5ccccc5c5ccnc(-c6cc([N+](=O)[O-])ccc6Cl)c54)c23)c1

O=[N+]([O-])c1ccc(Cl)c(-c2nccc3c4ccccc4n(CCCCCCCCCn4c5ccccc5c5ccnc(-c6cc([N+](=O)[O-])ccc6Cl)c54)c23)c1

CC1=C(CN2CCN(c3ccc(N4CC(Cn5cc(-c6ccccn6)nn5)OC4=O)cc3F)CC2)CC(c2ccc3c(c2)C3)N1c1ccc(Cl)c(Cl)c1

N#Cc1c(C23CC2C2(Cl)CC23)nc(SCC(=O)Nn2c(COc3ccc(Cl)cc3Cl)nc3ccccc3c2=O)n(Cc2ccccc2)c1=O

O=C(CCC1C=C2CC2(O)C(CNc2ccc(NC34CC3Nc3cc(Cl)ccc34)cc2)C1)C1=CCC2(C=C1)CO2

O=C(C(=CC12CCC3C1C32)Oc1ccc(CNNC23CC2Nc2cc(Cl)ccc23)cc1)c1ccc(Cl)cc1

O=C(c1ccc(Cl)cc1)C(CC12C(Cl)=CC3(Cl)C1C32)OC1C=CC(C=NNC23CC2Nc2cc(Cl)ccc23)=CC1

O=C(C(=Cc1cccc([N+](=O)[O-])c1)Oc1ccc(CNNC23c4ccc(Cl)cc4N4C2C43)cc1)C12C=CC(Cl)C1C2

O=C(NCCCCCNc1c2c(nc3ccccc13)CCCC2)c1cc2c3c4c(cc2oc1=O)C43

CC(C)(O)c1ccccc1CCC(SCC1(CC(=O)O)CC1)c1cccc(C=Cc2ccc3ccc(Cl)cc3n2)c1

C=C1OC(OC=C2OC(Oc3c(C4=CCC(=O)C=C4)oc4c(c3=O)C(=O)CC(=O)C4)C(=O)C(=O)C2=O)C(=O)C(OC2OC3COOC3C(=O)C2=O)C1=O

CCCC(=O)c1c(O)c(CC2C(=O)C(C(=O)C3CC3)=C(O)C(C)(CC=C(C)CCC3CC3C)C2O)c2c(c1O)C1CC(C)(O)CCC1C(C)(C)O2

C=CC(C)=CCC1(C)C(C)CC(O)C23C(=CC(OC(=O)c4ccccc4)CC12)C(OC(C)=O)OC3OC(C)=O

CCON=C(C(=O)NC1C(=O)N2C1SCC(Sc1nc(C3=CCN(C)C=C3)cs1)C2C(=O)O)c1nsc(NP(=O)(O)O)n1

O=C1CC(c2ccc(-c3ccccc3C(=O)O)o2)c2cn(Cc3cccc(F)c3)c3cccc(c23)N1

Cc1c(CC(=O)NC(CSCc2ccccc2)C(=O)O)c(=O)oc2cc3occ(-c4ccc(F)cc4)c3cc12

COc1cc(C2CC(=O)Oc3ccc4c(c32)OC(=Cc2ccco2)C4=O)ccc1OCCc1ccc2c(c1)CCO2

Cc1c(OCC(=O)NC(CSCc2ccccc2)C(=O)O)ccc2c(-c3ccccc3)cc(=O)oc12

CCC(C=C1OC(=O)C2=CC3CCC12C1C2=C(CCC31)C(=CCCC1CCCC1)OC2=O)Cc1ccccc1

Cc1c(CCC(=O)N2CCC3(O)CCCCC3C2)c(=O)oc2cc(OCc3ccc(C(C)(C)C)cc3)ccc12

Cc1oc2cc3oc(=O)c(CC(=O)N4CCC(C(=O)O)(c5ccccc5)CC4)c(C)c3cc2c1-c1ccccc1

Cc1c(CC(=O)NC(CSCc2ccccc2)C(=O)O)c(=O)oc2c(C)c3occ(-c4ccc(F)cc4)c3cc12

CC(C)CCC(C)NC1CCC(c2cc(O)cc(CC3CCCC(C4(C)C=CCC(C)C4)C3)c2)c2ccccc21

O=C(C=Cc1ccccc1)OC1C=CC(=O)OC1C(OC(=O)C=Cc1ccccc1)C(OC(=O)C=Cc1ccccc1)c1ccccc1

COc1ccc(C2CC(=O)Oc3ccc4c(=O)c(-c5ccc(O)cc5)coc4c32)cc1OCCc1ccc2c(c1)CCO2

COn1c2ccccc2c2cc(COCc3ccc4c(c3)c3cc(Cc5ccc6c(c5)c5ccccc5n6OC)ccc3n4OC)ccc21

O=C1CC(c2ccc(-c3ccccc3C(=O)O)o2)c2cn(Cc3ccccc3)c3cccc(c23)N1

Cc1ccc(S(=O)(=O)NC(C)C(=O)N2CCC(C(=O)NC(Cc3c[nH]c4ccccc34)C(=O)O)CC2)cc1

**Supplementary section C: Fluoresense assays results**

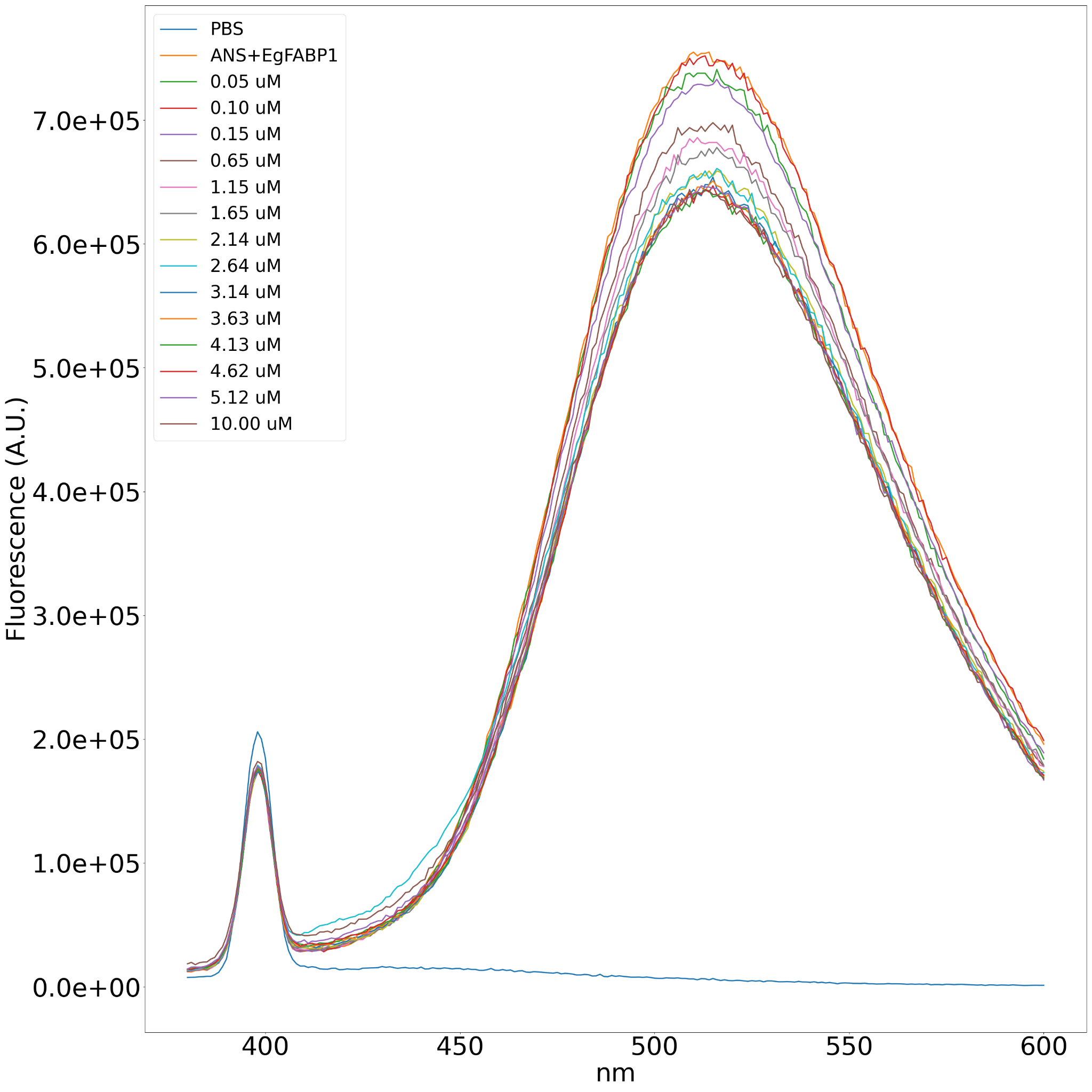

**Figure S5.** Fluorescence displacement spectrum for EgFABP1 with increasing concentrations of oleic acid. The concentration of ANS is 20 µM, while the protein concentration is 1 µM. This spectrum is representative of three independent replicas.

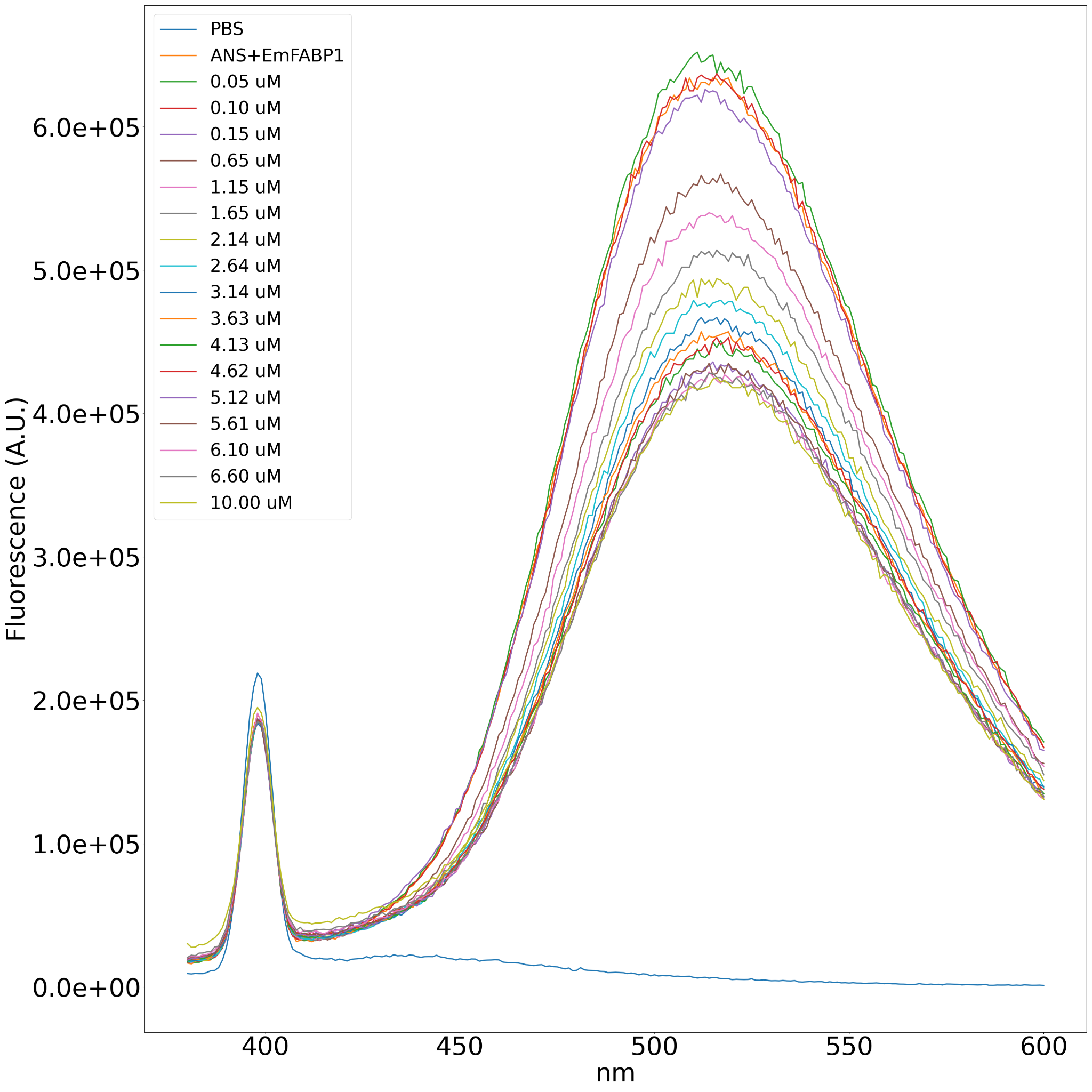
**Figure S6.** Fluorescence displacement spectrum for EmFABP1 with increasing concentrations of oleic acid. The concentration of ANS is 20 µM, while the protein concentration is 1 µM. This spectrum is representative of three independent replicas.

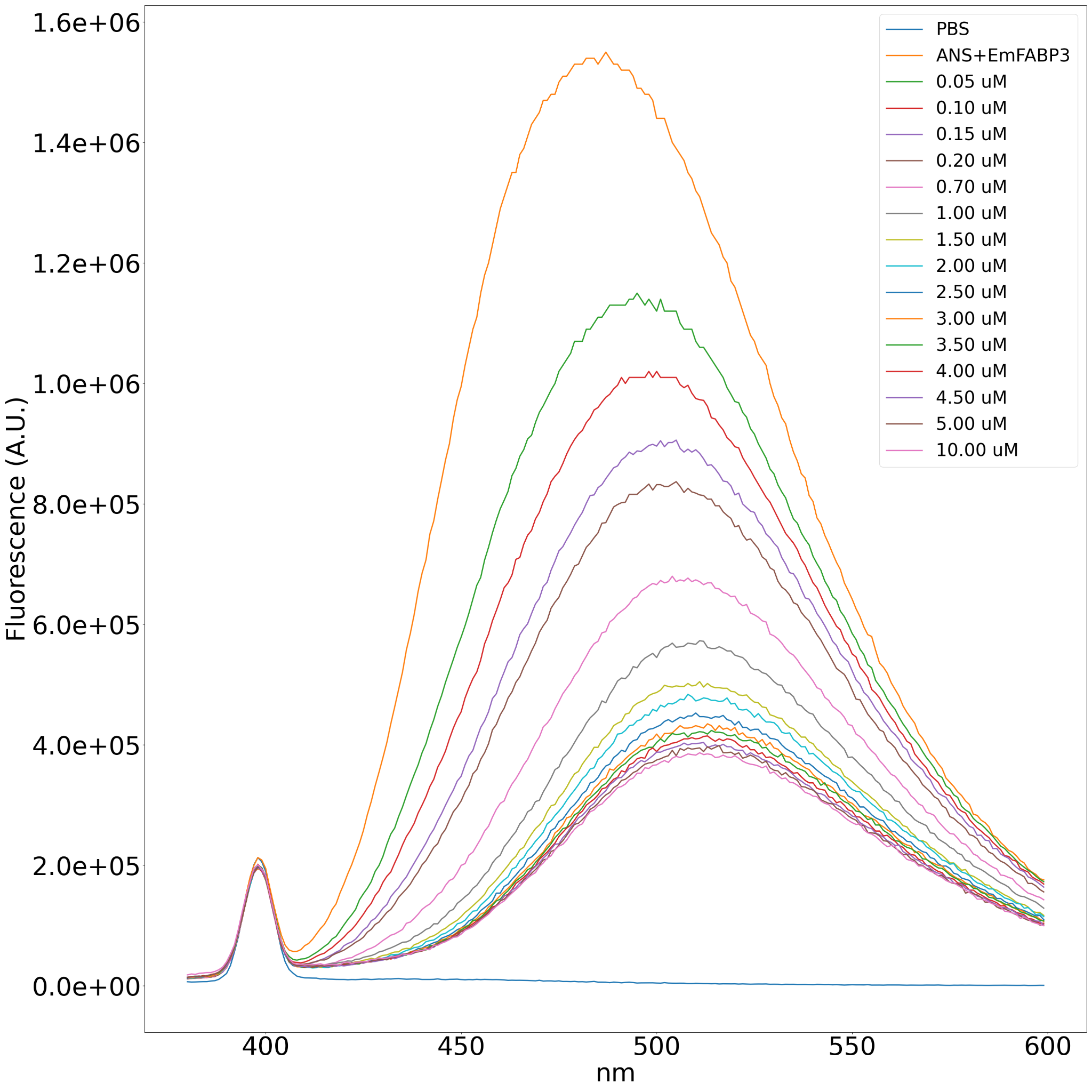
**Figure S7**. Fluorescence displacement spectrum for EmFABP3 with increasing concentrations of oleic acid. The concentration of ANS is 20 µM, while the protein concentration is 1 µM. This spectrum is representative of three independent replicas.

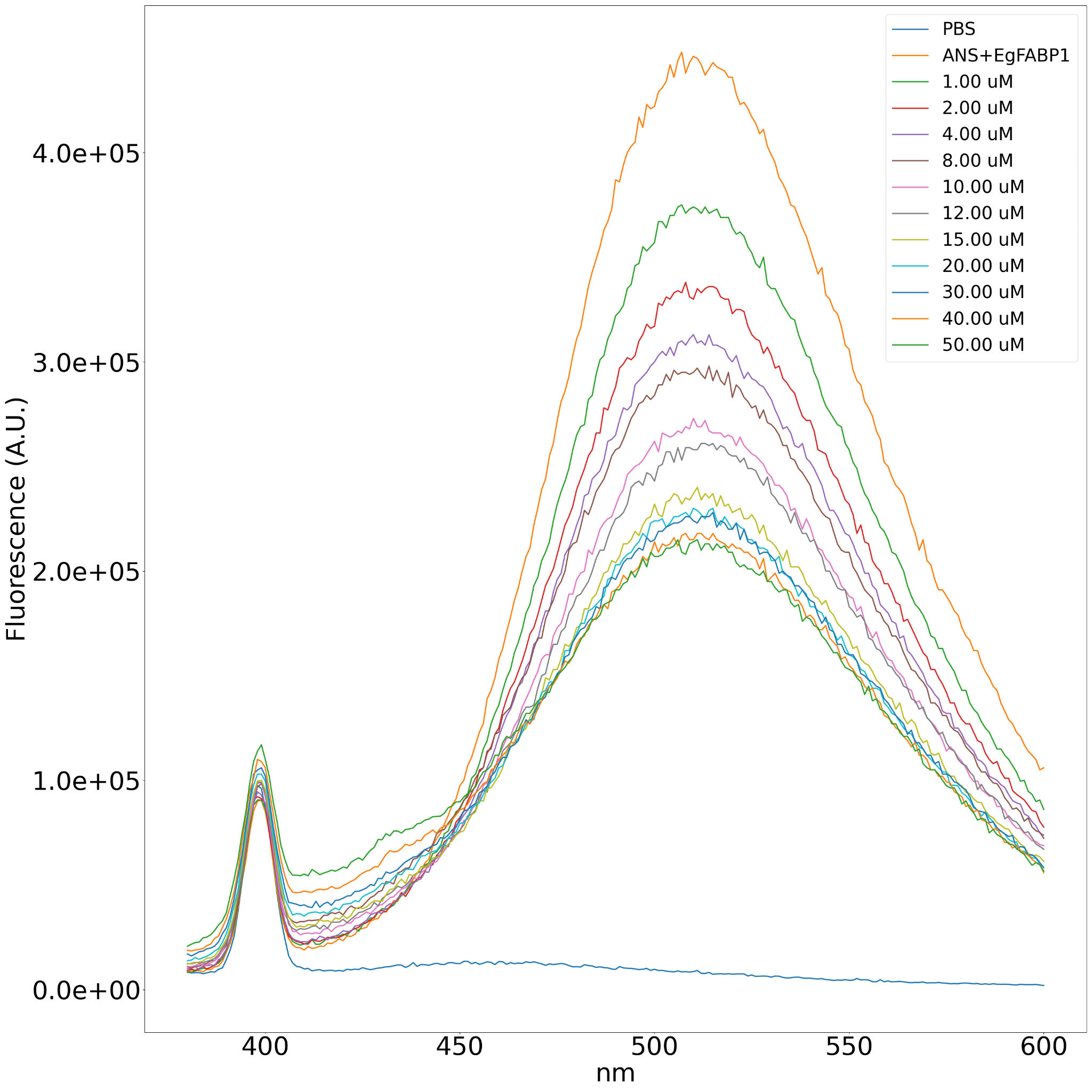
**Figure S8.** Fluorescence displacement spectrum for EgFABP1 with increasing concentrations of hydrochlorothiazide. The concentration of ANS is 20 µM, while the protein concentration is 1 µM. This spectrum is representative of three independent replicas.

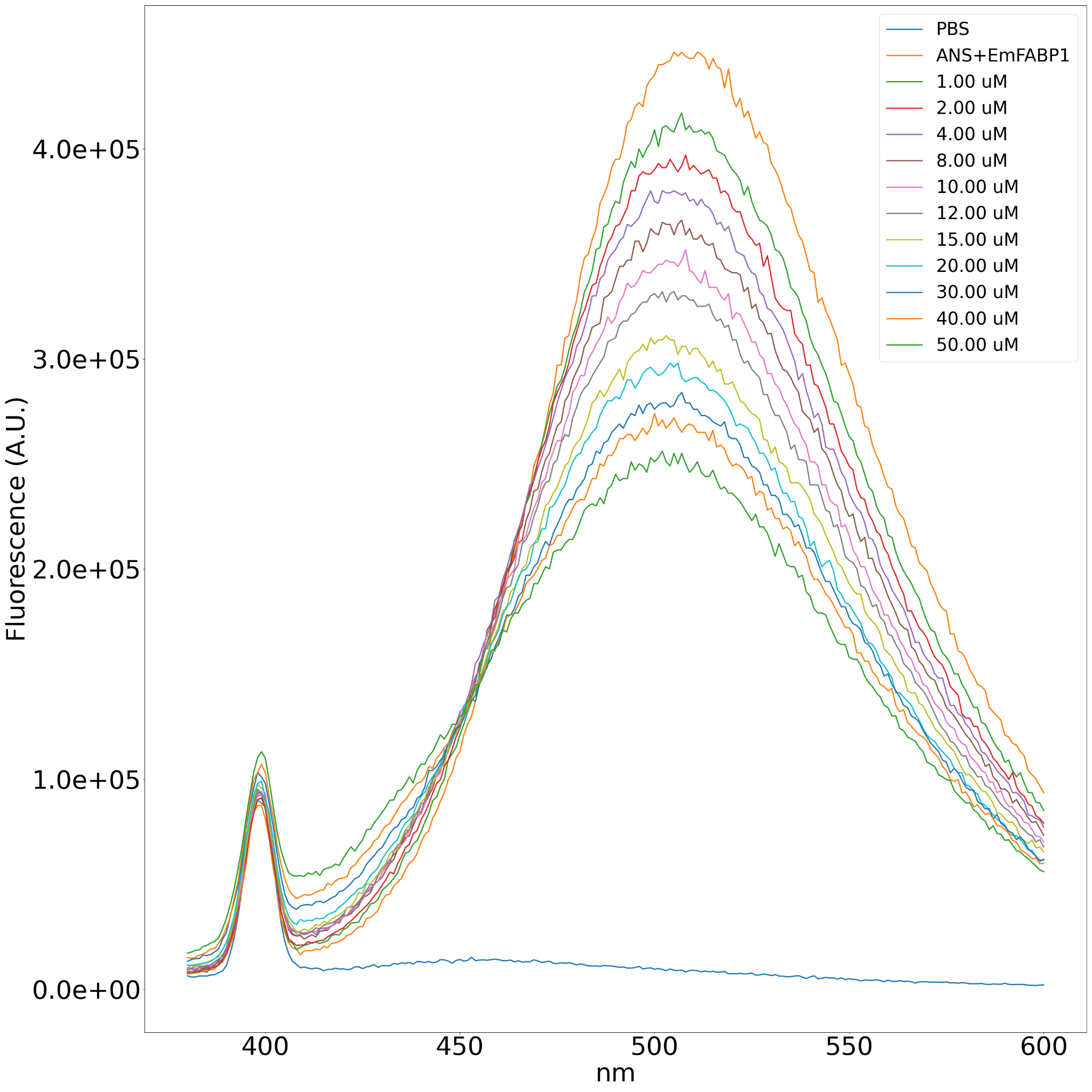
**Figure S9.** Fluorescence displacement spectrum for EmFABP1 with increasing concentrations of hydrochlorothiazide. The concentration of ANS is 20 µM, while the protein concentration is 1 µM. This spectrum is representative of three independent replicas.

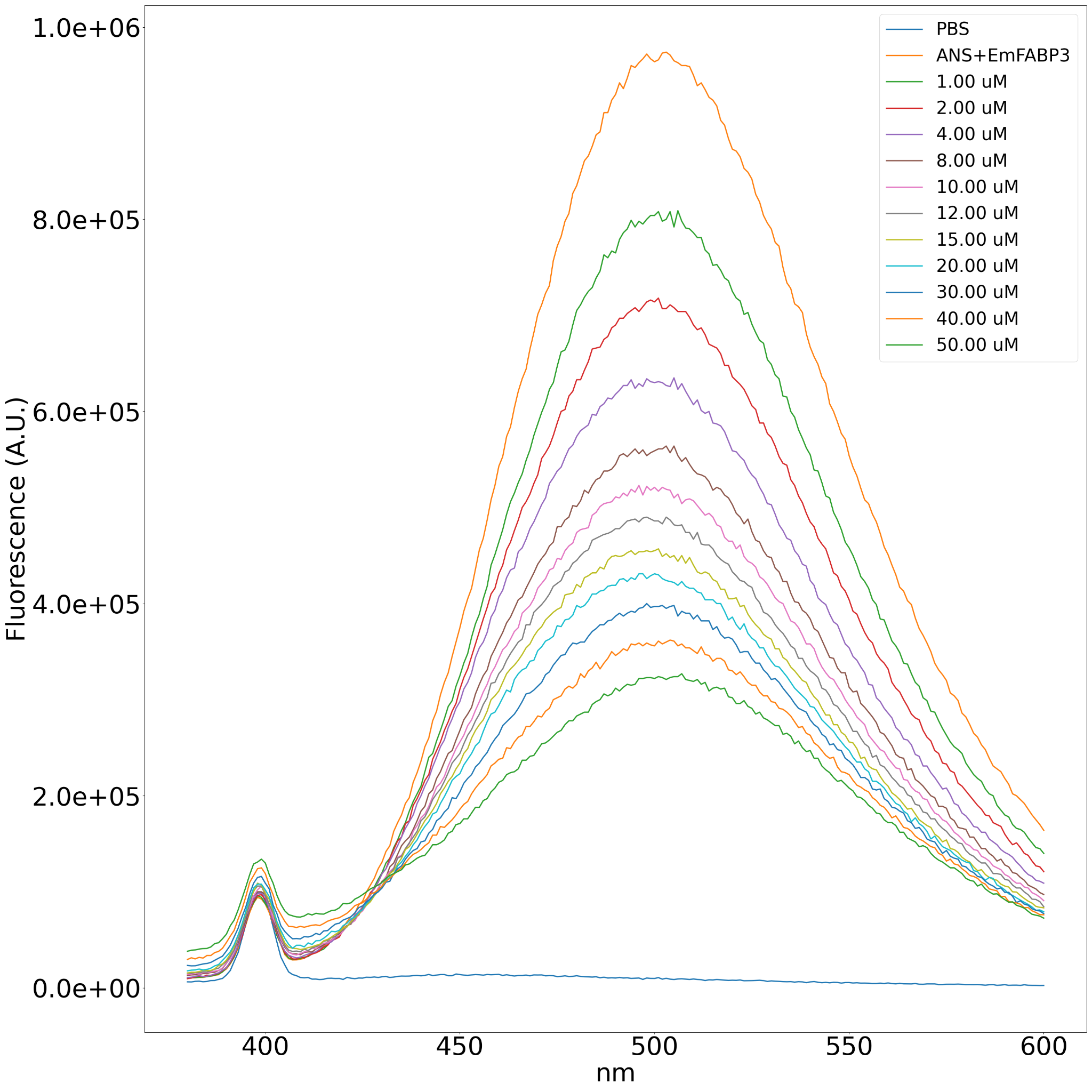
**Figure S10.** Fluorescence displacement spectrum for EmFABP3 with increasing concentrations of hydrochlorothiazide. The concentration of ANS is 20 µM, while the protein concentration is 1 µM. This spectrum is representative of three independent replicas.

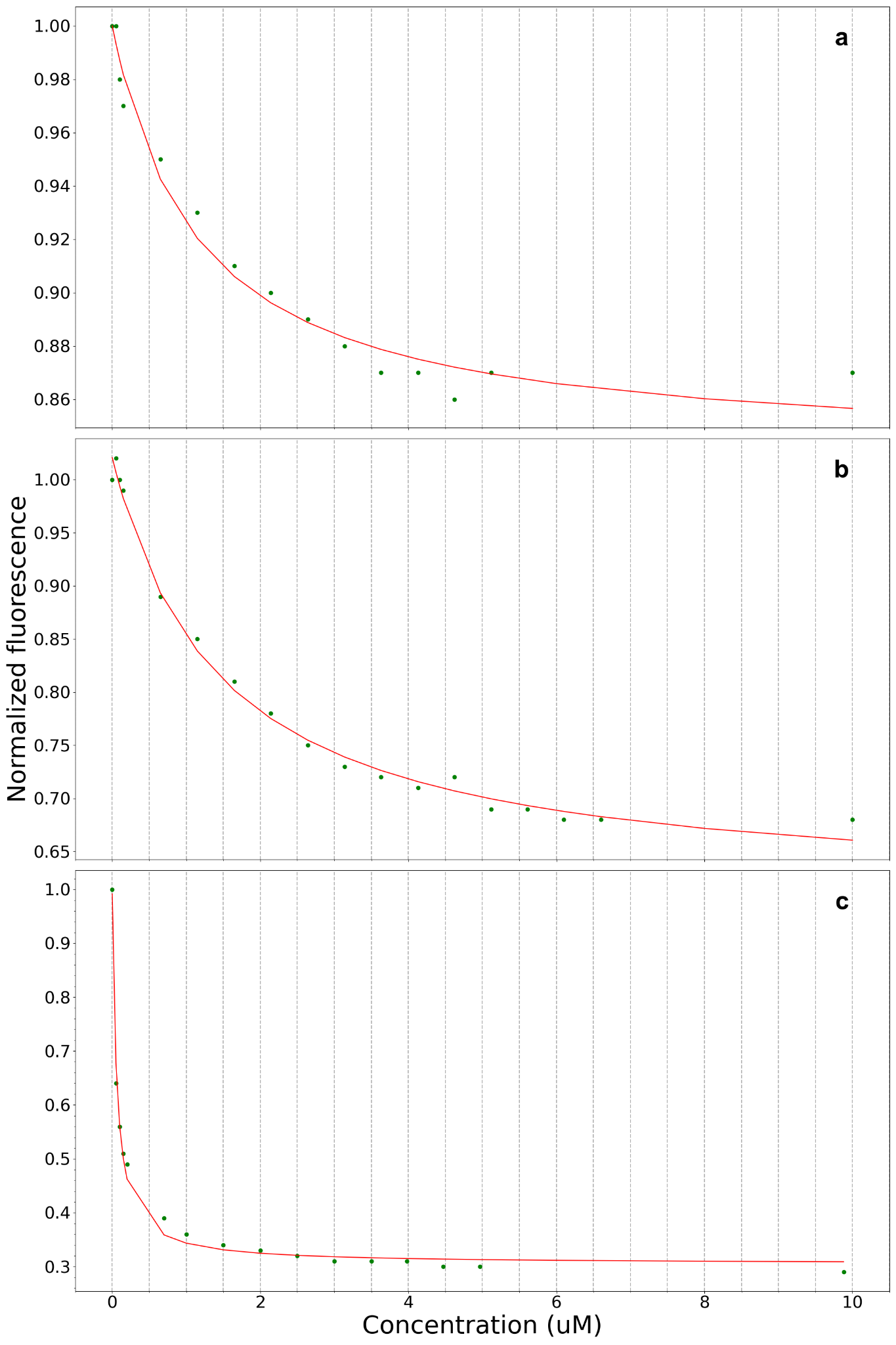

**Figure S11**. Fitting of integrated fluorescence data for oleic acid assayed in presence of (a) EgFABP1, (b) EmFABP1, (c) EmFABP3. Each plot is representative of three replicates in each case.

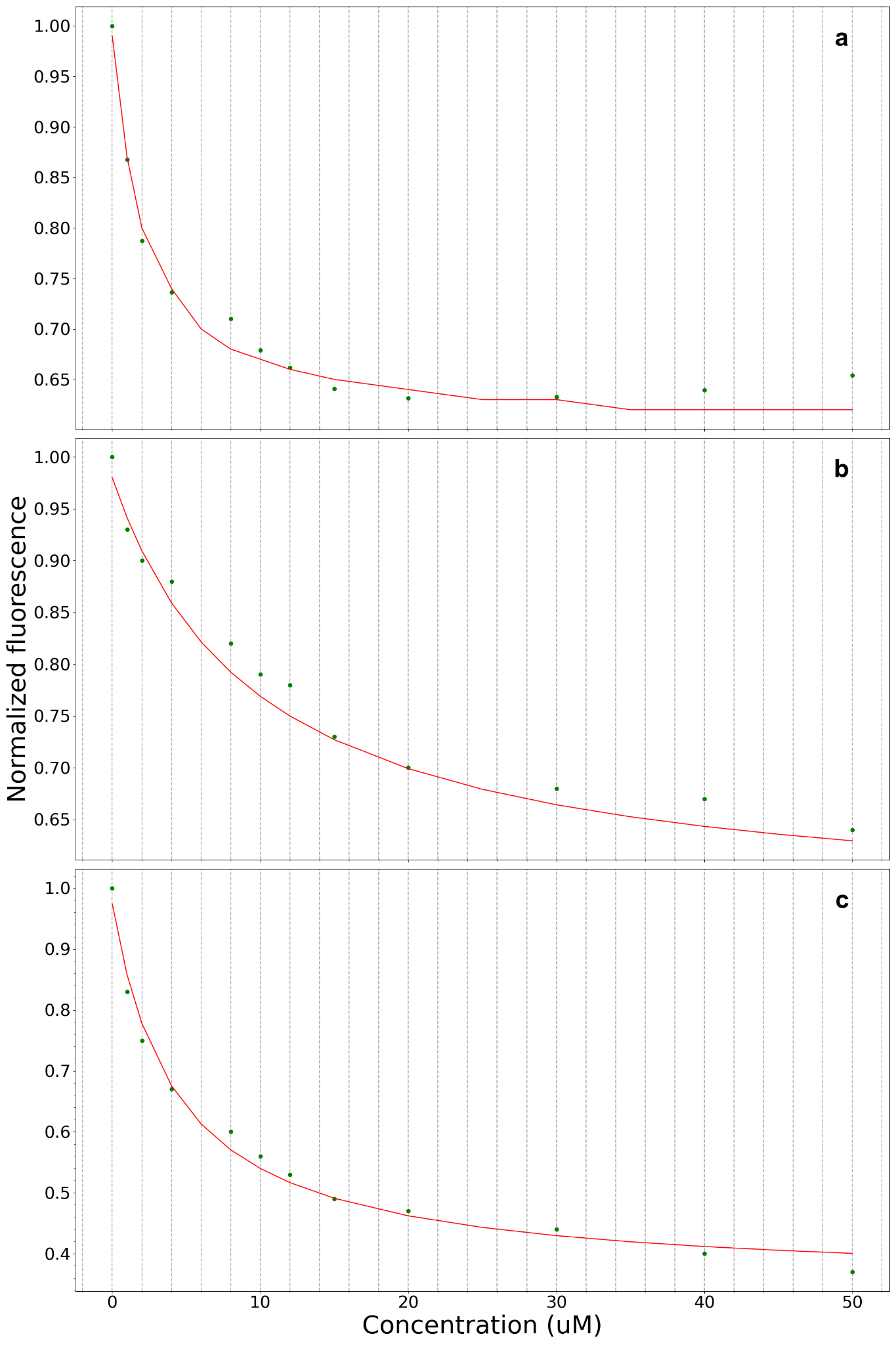

**Figure S12**. Fitting of integrated fluorescence data for hydrochlorothiazide assayed in presence of (a) EgFABP1, (b) EmFABP1, (c) EmFABP3. Each plot is representative of three replicates in each case.

We tested montelukast, fenticonazole, and naratriptan in the ANS displacement assay, but for different reasons it was not possible to reliably determine their 𝐾𝑑𝑎𝑝𝑝 values. As discussed in the main text, these cases illustrate the limitations of the assay and the need for complementary strategies to confirm ligand binding.

In the case of montelukast with EmFABP1, increased emission near the Raman peak without changes in the ANS emission maximum compromised fluorescence integration (**Figure S14**). For fenticonazole, fluorescence increased with its concentration both in the presence and absence of EmFABP1, preventing reliable assessment of ANS displacement (**Figure S15**). Similarly, naratriptan caused a general increase in fluorescence across the emission spectrum, making the assay unsuitable for 𝐾𝑑𝑎𝑝𝑝 estimation (**Figure S16**). However, a rightward shift in the ANS emission peak was observed in the presence of EmFABP3, suggesting a possible interaction, though not quantifiable with this method (**Figure S16**).

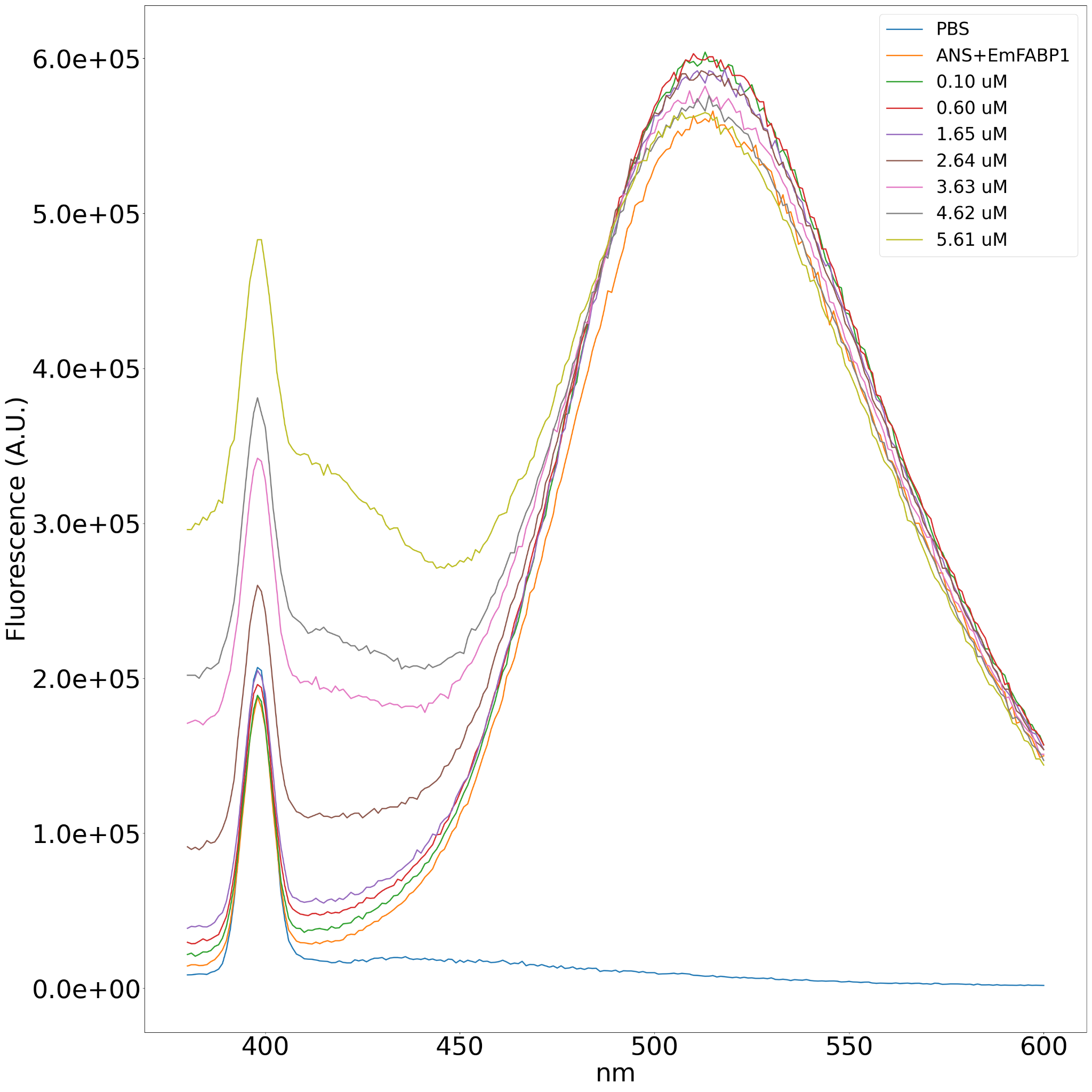
**Figure S13.** Fluorescence displacement spectrum for EmFABP1 with increasing concentrations of montelukast. The concentration of ANS is 20 µM, while the protein concentration is 1 µM.

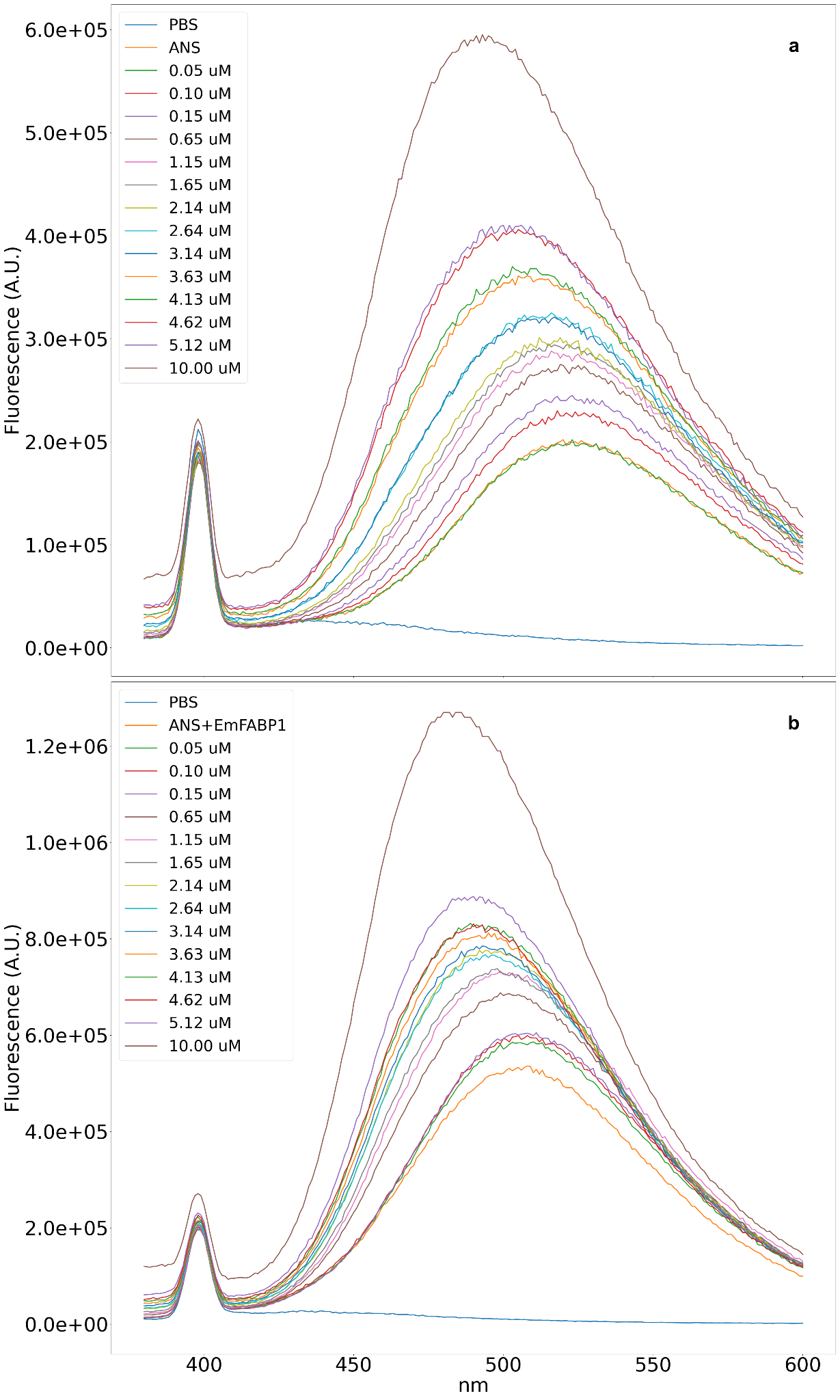

**Figure S14.** Fluorescence emission spectra for increasing concentrations of fenticonazole without (a) and with (b) EmFABP1. The concentration of ANS is 20 µM, while the protein concentration is 1 µM.

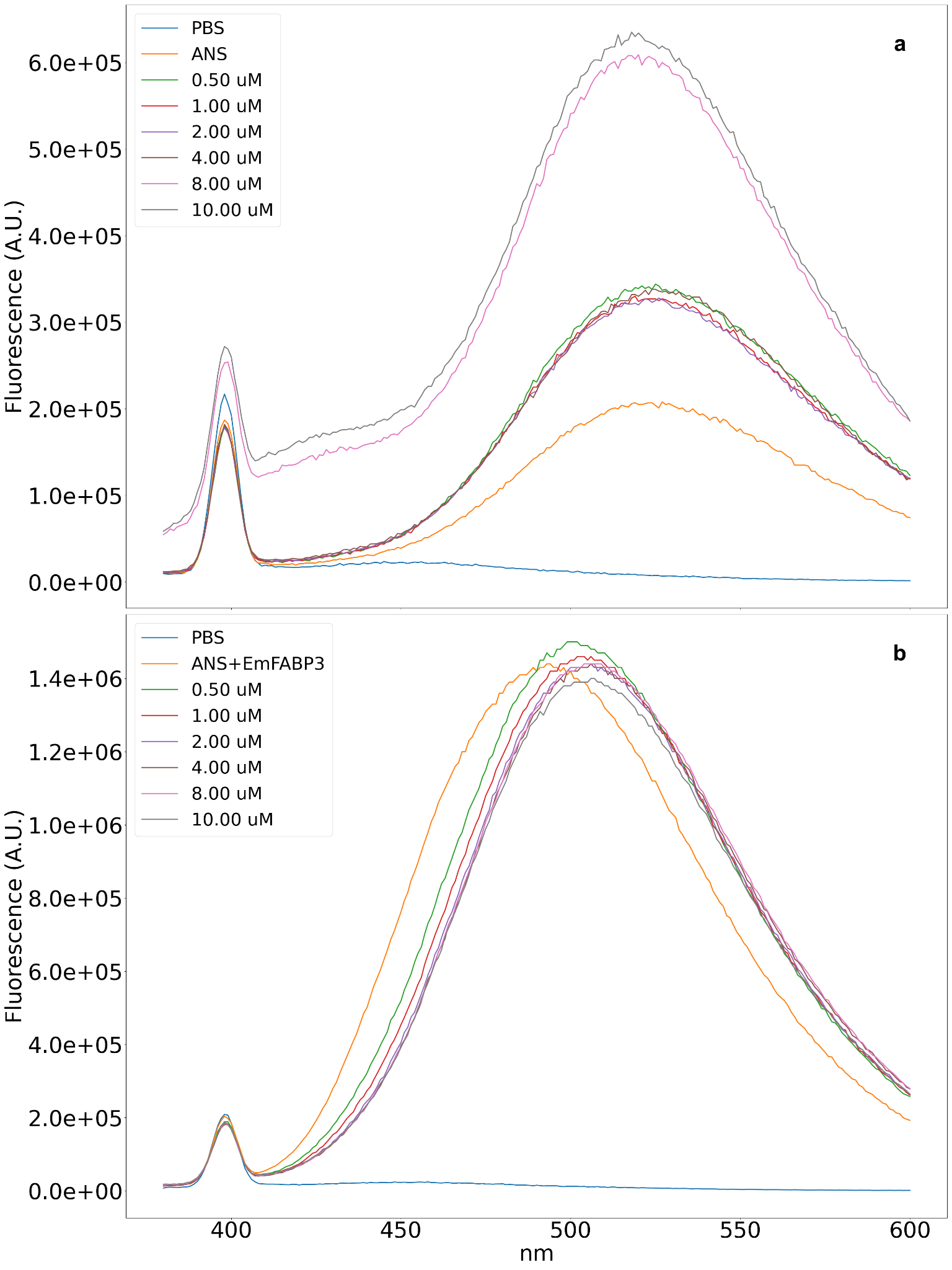

**Figure S15.** Fluorescence emission spectra for increasing concentrations of naratriptan without (a) and with (b) EmFABP3. The concentration of ANS is 20 µM, while the protein concentration is 1 µM.

**Supplementary section D: Dataset composition**

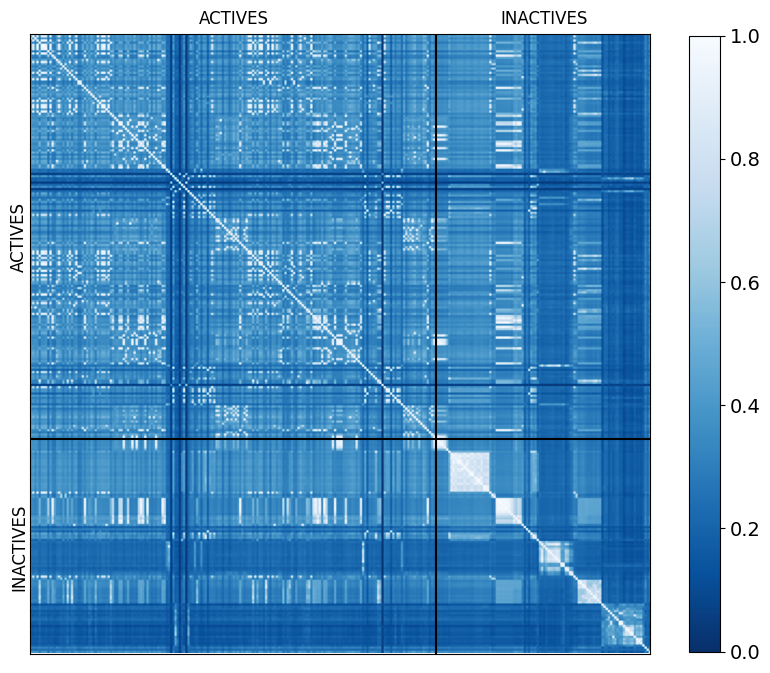
**Figure S16.** Similarity heatmap of ACTIVE and INACTIVE compounds from the compiled dataset.

**Table S3.** Compiled dataset used in this work.

| **MOLECULE** | **CATEGORY** | **SMILES** | **SET** |
| --- | --- | --- | --- |
| A1 | ACTIVE | O=C(O)c1cc(NS(=O)(=O)c2ccc(OCc3ccccc3)c3ccccc23)ccc1F | Training |
| A2 | ACTIVE | CCC(CC)c1nc2cc(F)c(Cl)cc2c(-c2ccccc2)c1-c1nnn[nH]1 | Training |
| A3 | ACTIVE | O=C(O)c1cc(NS(=O)(=O)c2ccc(OCCCC3CCCCC3)c3ccccc23)ccc1F | Training |
| A4 | ACTIVE | COc1ccc2ccn(S(=O)(=O)c3ccccc3C(=O)O)c2c1 | Training |
| A5 | ACTIVE | COc1c(-c2ccc(O)cc2)cc(S(=O)(=O)Nc2ccc(F)c(C(=O)O)c2)c2ccccc12 | Training |
| A6 | ACTIVE | Cc1cn(S(=O)(=O)c2ccsc2C(=O)O)c2ccccc12 | Training |
| A7 | ACTIVE | O=C(c1nc2cccc(Cl)c2nc1O)N1CCCC1Cc1cc(Cl)cc(Cl)c1 | Training |
| A8 | ACTIVE | COc1cccc2ccn(S(=O)(=O)c3ccsc3C(=O)O)c12 | Training |
| A9 | ACTIVE | O=C(O)c1cccc(NS(=O)(=O)c2ccc(Br)c3ccccc23)c1 | Training |
| A10 | ACTIVE | Cc1ccc2ccn(S(=O)(=O)c3ccsc3C(=O)O)c2c1 | Training |
| A11 | ACTIVE | O=C(O)c1cc(NS(=O)(=O)c2ccc(OCC3CCCCC3)c3ccccc23)ccc1F | Training |
| A12 | ACTIVE | CC(C)c1nc2ccc(Cl)cc2c(-c2ccccc2)c1C(=O)O | Training |
| A13 | ACTIVE | CCOC(=O)C(C)C(=O)CCc1cccc(-c2c(C(=O)O)c(C(C)C)nc3ccc(Cl)cc23)c1 | Training |
| A14 | ACTIVE | O=C(O)c1cccc2c3c(n(Cc4ccccc4)c12)CCCCC3 | Training |
| A15 | ACTIVE | O=C(Nc1cc(C(F)(F)F)ccc1-c1ccccc1)NC1(C(=O)O)CCCC1 | Training |
| A16 | ACTIVE | COc1c(-c2ccncc2)cc(S(=O)(=O)Nc2ccc(F)c(C(=O)O)c2)c2ccccc12 | Training |
| A17 | ACTIVE | Fc1cc2nc(N3CCCCC3)c(-c3nnn[nH]3)c(-c3ccccc3)c2cc1Cl | Training |
| A18 | ACTIVE | COc1c(-c2cccc(O)c2)cc(S(=O)(=O)Nc2ccc(F)c(C(=O)O)c2)c2ccccc12 | Training |
| A19 | ACTIVE | CCN(CC)c1nc2ccc(C(F)(F)F)cc2c(-c2ccccc2)c1-c1nnn[nH]1 | Training |
| A20 | ACTIVE | COc1ccc(CSc2nc(O)cc(C(F)(F)F)n2)cc1 | Training |
| A21 | ACTIVE | Cc1nc2ccc(Cl)cc2c(-c2ccccc2)c1C(=O)O | Training |
| A22 | ACTIVE | O=C(O)c1sccc1S(=O)(=O)n1ccc2cccc(F)c21 | Training |
| A23 | ACTIVE | CCCCCCCCOc1ccc(S(=O)(=O)Nc2ccc(F)c(C(=O)O)c2)c2ccccc12 | Training |
| A24 | ACTIVE | O=C(O)c1c(N2CCCCC2)nc2c(Cl)cc(Cl)cc2c1-c1ccccc1 | Training |
| A25 | ACTIVE | Cc1cc(Cl)cc2c(-c3ccccc3)c(C(=O)O)c(N3CCCCC3)nc12 | Training |
| A26 | ACTIVE | O=C(O)c1cc(NS(=O)(=O)c2ccc(OCCC3CCCCC3)c3ccccc23)ccc1F | Training |
| A27 | ACTIVE | CCOc1ccc(S(=O)(=O)Nc2ccc(F)c(C(=O)O)c2)c2ccccc12 | Training |
| A28 | ACTIVE | COc1c(-c2ccc(F)cc2)cc(S(=O)(=O)Nc2ccc(F)c(C(=O)O)c2)c2ccccc12 | Training |
| A29 | ACTIVE | O=C(O)c1c(N2CCCCC2)nc2ccc(Cl)cc2c1-c1ccccc1 | Training |
| A30 | ACTIVE | O=C(O)C=Cc1cn(Cc2ccccc2F)c2ccccc12 | Training |
| A31 | ACTIVE | Cc1cc(C)c(C)c(S(=O)(=O)Nc2ccc(O)c(Sc3nc[nH]n3)c2)c1C | Training |
| A32 | ACTIVE | CCN(C)c1nc2ccc(Cl)cc2c(-c2ccccc2)c1-c1nnn[nH]1 | Training |
| A33 | ACTIVE | O=C(O)Cc1cn(-c2ccccc2-c2nc(-c3ccc4ccccc4c3)c(-c3ccccc3)s2)nn1 | Training |
| A34 | ACTIVE | FC(F)(F)c1ccc2nc(N3CCCCC3)c(-c3nnn[nH]3)c(-c3ccccc3)c2c1 | Training |
| A35 | ACTIVE | CCN(C)c1nc2c(Cl)cc(Cl)cc2c(-c2ccccc2)c1C(=O)O | Training |
| A36 | ACTIVE | O=C(O)c1cc(S(=O)(=O)n2ccc3ccc(Br)cc32)cs1 | Training |
| A37 | ACTIVE | O=C(O)c1ccc2c3c(n(Cc4ccccc4)c2c1)CCCC3 | Training |
| A38 | ACTIVE | COc1ccc(Cl)cc1NCc1cc(=O)n2nc(-c3ccccc3)nc2[nH]1 | Training |
| A39 | ACTIVE | CN(Cc1ccc(Cl)cc1)c1nc(O)cc(C(F)(F)F)n1 | Training |
| A40 | ACTIVE | COc1ccc(S(=O)(=O)Nc2ccccc2C(=O)O)cc1C(C)(C)C | Training |
| A41 | ACTIVE | O=C(O)CCCOc1ccccc1-c1cc(-c2ccccc2)n(-c2ccc(Br)cc2)n1 | Training |
| A42 | ACTIVE | O=C(O)COc1cccc(-c2ccccc2-c2cc(-c3ccccc3)c(-c3ccccc3)o2)c1 | Training |
| A43 | ACTIVE | CCc1c(-c2ccccc2)c(-c2ccccc2)nn1-c1ccccc1-c1cccc(OCC(=O)O)c1 | Training |
| A44 | ACTIVE | CCc1c(-c2ccccc2)c(-c2ccccc2)nn1-c1ccccc1-c1ccc(OCC(=O)O)cc1 | Training |
| A45 | ACTIVE | O=C(Nc1cc(Cl)c(F)cc1-c1ccccc1)NC1(C(=O)O)CCCC1 | Training |
| A46 | ACTIVE | O=C(O)CCCOc1ccccc1-c1cc(-c2ccc(Cl)cc2)n(-c2ccccc2)n1 | Training |
| A47 | ACTIVE | O=C(O)CCc1ccc(O)c(Cc2ccccc2)c1 | Training |
| A48 | ACTIVE | O=C(O)Cc1cn(-c2ccccc2-c2nc(-c3ccc(Cl)cc3)c(-c3ccccc3)s2)nn1 | Training |
| A49 | ACTIVE | O=C(Nc1cc(Cl)ccc1-c1ccccc1)NC1(C(=O)O)CCCC1 | Training |
| A50 | ACTIVE | Cn1c(-c2ccccc2-c2cccc(NCC(=O)O)c2)nc(-c2ccccc2)c1-c1ccccc1 | Training |
| A51 | ACTIVE | O=C(O)C1C2CC=CC2c2cc(Cl)cc3c2N1CC1CC=CC31 | Training |
| A52 | ACTIVE | COc1ccc(C(C)C)cc1S(=O)(=O)Nc1cccc(C(=O)O)c1 | Training |
| A53 | ACTIVE | CCc1c(-c2ccccc2-c2cccc(OCC(=O)O)c2)oc(-c2ccccc2)c1-c1ccccc1 | Training |
| A54 | ACTIVE | O=C(O)CCCOc1ccccc1-c1cc(-c2ccccc2)n(-c2ccccc2Cl)n1 | Training |
| A55 | ACTIVE | CCc1c(-c2ccccc2)c(-c2ccccc2)nn1-c1ccccc1-c1cccc(OC(CC)C(=O)O)c1 | Training |
| A56 | ACTIVE | O=S(=O)(Nc1ccc(O)c(Sc2nc[nH]n2)c1)c1cccc2ccccc12 | Training |
| A57 | ACTIVE | CCc1c(-c2ccccc2)c(-c2ccccc2)nn1-c1ccccc1-c1cccc(OC(C)C(=O)O)c1 | Training |
| A58 | ACTIVE | COc1ccc(C(C)(C)C)cc1S(=O)(=O)Nc1cccc(C(=O)O)c1 | Training |
| A59 | ACTIVE | Oc1cc(C(F)(F)F)nc(NCc2ccc(Cl)cc2)n1 | Training |
| A60 | ACTIVE | COc1ccc(C(C)(C)C)cc1S(=O)(=O)Nc1ccccc1C(=O)O | Training |
| A61 | ACTIVE | O=C(O)COc1cccc(-c2ccccc2-c2cc(-c3ccccc3)c(-c3ccccc3)[nH]2)c1 | Training |
| A62 | ACTIVE | CCCCCCCCC=CCCCCCCCC(=O)O | Training |
| A63 | ACTIVE | O=C(O)CCOc1ccccc1-c1cc(-c2ccccc2)n(-c2ccccc2)n1 | Training |
| A64 | ACTIVE | Cc1cc(-c2c3c(nc(C4CCCC4)c2-c2nnn[nH]2)CCC(C)C3)ccn1 | Training |
| A65 | ACTIVE | O=C(O)CNc1cccc(-c2ccccc2-c2nc(-c3ccccc3)c(-c3ccccc3)n2CCF)c1 | Training |
| A66 | ACTIVE | O=C(O)C1=CC=CC2C1NC(c1ccccc1[N+](=O)[O-])C1CC(Sc3ccccc3[N+](=O)[O-])C(Cl)C21 | Training |
| A67 | ACTIVE | Cc1cn(S(=O)(=O)c2csc(C(=O)O)c2)c2ccccc12 | Training |
| A68 | ACTIVE | O=C(O)COc1cccc(-c2ccccc2-c2nc(-c3ccccc3)c(-c3ccccc3)[nH]2)c1 | Training |
| A69 | ACTIVE | O=C(Nc1cc(Cl)ccc1-c1ccc(F)cc1)NC1(C(=O)O)CCCC1 | Training |
| A70 | ACTIVE | CCN(CC)c1nc2cc(F)c(Cl)cc2c(-c2ccccc2)c1-c1nnn[nH]1 | Training |
| A71 | ACTIVE | CC(Oc1cccc(-c2ccccc2-c2cc(-c3cccs3)n(-c3ccccc3)n2)c1)C(=O)O | Training |
| A72 | ACTIVE | CCOc1ccc(C(C)C)cc1S(=O)(=O)Nc1cccc(C(=O)O)c1 | Training |
| A73 | ACTIVE | O=C(Sc1nc(O)cc(C(F)(F)F)n1)N1CCCCC1 | Training |
| A74 | ACTIVE | CC(C)c1cc(C(C)C)c(C(=O)O)c(C(C)C)c1 | Training |
| A75 | ACTIVE | COc1ccc(Nc2nc(O)cc(C(F)(F)F)n2)cc1 | Training |
| A76 | ACTIVE | CCC(C)n1c(-c2ccccc2-c2cccc(NCC(=O)O)c2)nc(-c2ccccc2)c1-c1ccccc1 | Training |
| A77 | ACTIVE | CCn1c(-c2ccccc2-c2cccc(NCC(=O)O)c2)nc(-c2ccccc2)c1-c1ccccc1 | Training |
| A78 | ACTIVE | CC(C)c1cc(C(C)C)c(S(=O)(=O)O)c(C(C)C)c1 | Training |
| A79 | ACTIVE | Cc1ccc(C=c2[nH]c(=O)c(=Cc3cccn3-c3cccc(C(=O)O)c3)s2)cc1C | Training |
| A80 | ACTIVE | Cn1c(-c2ccccc2-c2cccc(OCC(=O)O)c2)nc(-c2ccccc2)c1-c1ccccc1 | Training |
| A81 | ACTIVE | O=C(O)CCCOc1ccccc1-c1cc(-c2ccccc2)n(-c2ccc(Cl)cc2)n1 | Training |
| A82 | ACTIVE | CCOc1ccc(S(=O)(=O)Nc2cccc(C(=O)O)c2)c(C)c1C | Training |
| A83 | ACTIVE | Cc1ccc(Cn2c3c(c4cccc(C(=O)O)c42)CCCC3)cc1 | Training |
| A84 | ACTIVE | CCCc1cn(Cc2ccccc2)c2c(C(=O)O)cccc12 | Training |
| A85 | ACTIVE | Cc1c(C)n(Cc2ccccc2)c2c(C(=O)O)cccc12 | Training |
| A86 | ACTIVE | O=C(O)c1cccc2c3c(n(Cc4cccc(F)c4)c12)CCCCC3 | Training |
| A87 | ACTIVE | CCCn1c2c(c3cccc(C(=O)O)c31)CCCC2 | Training |
| A88 | ACTIVE | O=C(O)c1cccc2c3c(n(Cc4ccc(F)cc4)c12)CCCCC3 | Training |
| A89 | ACTIVE | CCCn1c2c(c3cccc(C(=O)O)c31)CCCCC2 | Training |
| A90 | ACTIVE | O=C(O)c1cccc2c3c(n(Cc4ccc(C(F)(F)F)cc4)c12)CCCCC3 | Training |
| A91 | ACTIVE | COc1cccc(Cn2c3c(c4cccc(C(=O)O)c42)CCCC3)c1 | Training |
| A92 | ACTIVE | O=C(O)c1cccc2c3c(n(Cc4cccc(F)c4)c12)CCCC3 | Training |
| A93 | ACTIVE | O=C(O)c1cccc2c3c(n(Cc4ccc(C(F)(F)F)cc4)c12)CCCC3 | Training |
| A94 | ACTIVE | O=C(O)c1cccc2c3c(n(Cc4ccccc4F)c12)CCCCC3 | Training |
| A95 | ACTIVE | COc1ccccc1Cn1c2c(c3cccc(C(=O)O)c31)CCCC2 | Training |
| A96 | ACTIVE | O=C(O)CCCN1C2C=CC=CC2C2C=CC=CC21 | Training |
| A97 | ACTIVE | CCCOc1cc(S(=O)(=O)Nc2ccc(C(=O)O)cc2)ccc1Cl | Training |
| A98 | ACTIVE | O=C(O)c1ccccc1S(=O)(=O)N1C2C=CC=CC2C2C=CC=CC21 | Training |
| A99 | ACTIVE | Cc1cc(C)c2c3c1C1C=CCC1CN3C(C(=O)O)C1CC=CC21 | Training |
| A100 | ACTIVE | O=C(O)c1sccc1S(=O)(=O)N1C2C=CC=CC2C2C=CC=CC21 | Training |
| A101 | ACTIVE | O=C(O)C1C2CC(Sc3ccccc3[N+](=O)[O-])C(Cl)C2c2cccc3c2N1CC1CC=CC31 | Training |
| I1 | INACTIVE | O=C(O)COc1ccccc1-c1cc(-c2ccc(Cl)cc2)n(-c2ccccc2)n1 | Training |
| I2 | INACTIVE | O=C(O)CCCCOc1ccccc1-c1cc(-c2ccc(Cl)cc2)n(-c2ccccc2)n1 | Training |
| I3 | INACTIVE | O=C(O)CCCCCCOc1ccccc1-c1cc(-c2ccc(Cl)cc2)n(-c2ccccc2)n1 | Training |
| I4 | INACTIVE | Cc1ccc(-c2cc(-c3ccccc3OCCCCC(=O)O)nn2-c2ccccc2)cc1 | Training |
| I5 | INACTIVE | O=C(O)CCCCOc1ccccc1-c1cc(-c2ccc(Br)cc2)n(-c2ccccc2)n1 | Training |
| I6 | INACTIVE | COc1ccccc1-n1nc(-c2ccccc2OCCCC(=O)O)cc1-c1ccccc1 | Training |
| I7 | INACTIVE | O=C(O)CCCOc1ccccc1-c1cc(-c2ccccc2)n(C2CCCCC2)n1 | Training |
| I8 | INACTIVE | O=C(O)CN1C2C=CC=CC2C2C=CC=CC21 | Training |
| I9 | INACTIVE | NC(=O)Cc1cn(S(=O)(=O)c2ccsc2C(=O)O)c2ccccc12 | Training |
| I10 | INACTIVE | Cc1ccc2c(ccn2S(=O)(=O)c2ccccc2C(=O)O)c1 | Training |
| I11 | INACTIVE | COc1cccc2c1ccn2S(=O)(=O)c1ccccc1C(=O)O | Training |
| I12 | INACTIVE | NC(=O)Cc1cn(S(=O)(=O)c2ccccc2C(=O)O)c2ccccc12 | Training |
| I13 | INACTIVE | CC(C)c1cc(C(C)C)c(S(=O)(=O)n2cnc3ccccc32)c(C(C)C)c1 | Training |
| I14 | INACTIVE | Cc1nc2ccccc2n1S(=O)(=O)c1c(C(C)C)cc(C(C)C)cc1C(C)C | Training |
| I15 | INACTIVE | Cc1ccc2c(c1)ncn2S(=O)(=O)c1c(C(C)C)cc(C(C)C)cc1C(C)C | Training |
| I16 | INACTIVE | Cc1ccc2nc(C)n(S(=O)(=O)c3c(C(C)C)cc(C(C)C)cc3C(C)C)c2c1 | Training |
| I17 | INACTIVE | Cc1ccc2c(c1)nc(C)n2S(=O)(=O)c1c(C(C)C)cc(C(C)C)cc1C(C)C | Training |
| I18 | INACTIVE | Cc1nc2ccc(Br)cc2n1S(=O)(=O)c1c(C(C)C)cc(C(C)C)cc1C(C)C | Training |
| I19 | INACTIVE | CC(C)c1cc(C(C)C)c(S(=O)(=O)n2cnc3cc(Br)ccc32)c(C(C)C)c1 | Training |
| I20 | INACTIVE | Cc1nc2cc(Br)ccc2n1S(=O)(=O)c1c(C(C)C)cc(C(C)C)cc1C(C)C | Training |
| I21 | INACTIVE | Cc1nc2ccc(Cl)cc2n1S(=O)(=O)c1c(C(C)C)cc(C(C)C)cc1C(C)C | Training |
| I22 | INACTIVE | CC(C)c1cc(C(C)C)c(S(=O)(=O)n2cnc3ccc(Cl)cc32)c(C(C)C)c1 | Training |
| I23 | INACTIVE | CC(C)c1cc(C(C)C)c(S(=O)(=O)n2cnc3cc(Cl)ccc32)c(C(C)C)c1 | Training |
| I24 | INACTIVE | COc1ccc2nc(C)n(S(=O)(=O)c3c(C(C)C)cc(C(C)C)cc3C(C)C)c2c1 | Training |
| I25 | INACTIVE | COc1ccc2ncn(S(=O)(=O)c3c(C(C)C)cc(C(C)C)cc3C(C)C)c2c1 | Training |
| I26 | INACTIVE | COc1ccc2c(c1)ncn2S(=O)(=O)c1c(C(C)C)cc(C(C)C)cc1C(C)C | Training |
| I27 | INACTIVE | COc1ccc2c(c1)nc(C)n2S(=O)(=O)c1c(C(C)C)cc(C(C)C)cc1C(C)C | Training |
| I28 | INACTIVE | CC(C)c1cc(C(C)C)c(S(=O)(=O)n2cnc3ccc([N+](=O)[O-])cc32)c(C(C)C)c1 | Training |
| I29 | INACTIVE | CC(C)c1cc(C(C)C)c(S(=O)(=O)n2cnc3cc([N+](=O)[O-])ccc32)c(C(C)C)c1 | Training |
| I30 | INACTIVE | CC(C)c1cc(C(C)C)c(S(=O)(=O)n2cnc3ccc(N)cc32)c(C(C)C)c1 | Training |
| I31 | INACTIVE | CC(C)c1cc(C(C)C)c(S(N)(=O)=O)c(C(C)C)c1 | Training |
| I32 | INACTIVE | O=C(CNc1nc(O)cc(C(F)(F)F)n1)N1CCCCC1 | Training |
| I33 | INACTIVE | Oc1cc(C(F)(F)F)nc(NCc2ccccc2)n1 | Training |
| I34 | INACTIVE | Oc1cc(C(F)(F)F)nc(Nc2ccccc2)n1 | Training |
| I35 | INACTIVE | Oc1cc(C(F)(F)F)nc(NCCc2ccccc2)n1 | Training |
| I36 | INACTIVE | Oc1cc(C(F)(F)F)nc(NCc2cccc(Cl)c2)n1 | Training |
| I37 | INACTIVE | Oc1cc(C(F)(F)F)nc(NCc2ccccc2Cl)n1 | Training |
| I38 | INACTIVE | Oc1cc(C(F)(F)F)nc(NCc2ccc(-c3ccccc3)cc2)n1 | Training |
| I39 | INACTIVE | Oc1cc(C(F)(F)F)nc(NCc2ccncc2)n1 | Training |
| I40 | INACTIVE | Cc1ccc(CNc2nc(O)cc(C(F)(F)F)n2)cc1 | Training |
| I41 | INACTIVE | CN(Cc1ccccc1)c1nc(O)cc(C(F)(F)F)n1 | Training |
| I42 | INACTIVE | Oc1cc(C(F)(F)F)nc(CCc2ccccc2)n1 | Training |
| I43 | INACTIVE | COc1ccc(CCc2nc(O)cc(C(F)(F)F)n2)cc1 | Training |
| I44 | INACTIVE | Cc1cc(O)nc(NCc2ccc(Cl)cc2)n1 | Training |
| I45 | INACTIVE | CCc1cc(O)nc(NCc2ccc(Cl)cc2)n1 | Training |
| I46 | INACTIVE | Oc1cc(-c2ccccc2)nc(NCc2ccc(Cl)cc2)n1 | Training |
| I47 | INACTIVE | Oc1cc(-c2ccccc2F)nc(NCc2ccc(Cl)cc2)n1 | Training |
| I48 | INACTIVE | O=C(O)c1ccc2c(c1)c1c(n2Cc2ccccc2)CCCC1 | Training |
| I49 | INACTIVE | O=C1CCCc2c1c1cccc(C(=O)O)c1n2Cc1ccccc1 | Training |
| I50 | INACTIVE | O=C(c1nc2ccccc2nc1O)N1CCCC1CN1CCCCC1 | Training |
| I51 | INACTIVE | O=C(c1nc2ccccc2nc1O)N1CCCC1Cn1cccn1 | Training |
| I52 | INACTIVE | O=C(OCc1ccccc1)C1CCCN1C(=O)c1nc2ccccc2nc1O | Training |
| I53 | INACTIVE | CC(C)Oc1ccc(S(=O)(=O)Nc2ccc(F)c(C(=O)O)c2)c2ccccc12 | Training |
| I54 | INACTIVE | CCC(C)Oc1ccc(S(=O)(=O)Nc2ccc(F)c(C(=O)O)c2)c2ccccc12 | Training |
| I55 | INACTIVE | CCCC(C)Oc1ccc(S(=O)(=O)Nc2ccc(F)c(C(=O)O)c2)c2ccccc12 | Training |
| I56 | INACTIVE | CCCCCCCCCCCCOc1ccc(S(=O)(=O)Nc2ccc(F)c(C(=O)O)c2)c2ccccc12 | Training |
| I57 | INACTIVE | O=C(O)c1cc(NS(=O)(=O)c2ccc(OC3CCCC3)c3ccccc23)ccc1F | Training |
| I58 | INACTIVE | O=C(O)c1cc(NS(=O)(=O)c2ccc(OC3CCCCCCC3)c3ccccc23)ccc1F | Training |
| I59 | INACTIVE | O=C(O)c1cc(NS(=O)(=O)c2ccc(OCC3CCOCC3)c3ccccc23)ccc1F | Training |
| I60 | INACTIVE | O=C(O)c1cc(NS(=O)(=O)c2ccc(Oc3ccccc3)c3ccccc23)ccc1F | Training |
| I61 | INACTIVE | COc1c(-c2cncnc2)cc(S(=O)(=O)Nc2ccc(F)c(C(=O)O)c2)c2ccccc12 | Training |
| I62 | INACTIVE | COc1c(-c2ccnn2C)cc(S(=O)(=O)Nc2ccc(F)c(C(=O)O)c2)c2ccccc12 | Training |
| I63 | INACTIVE | COc1c(-c2ccc(CO)cc2)cc(S(=O)(=O)Nc2ccc(F)c(C(=O)O)c2)c2ccccc12 | Training |
| I64 | INACTIVE | COc1ccc(-c2cc(S(=O)(=O)Nc3ccc(F)c(C(=O)O)c3)c3ccccc3c2OC)cc1OC | Training |
| I65 | INACTIVE | COc1ccc(OC)c(S(=O)(=O)Nc2ccccc2C(=O)O)c1 | Training |
| I66 | INACTIVE | COc1cc(S(=O)(=O)Nc2ccccc2C(=O)O)c(OC)cc1C | Training |
| I67 | INACTIVE | COc1cc(C)c(C)cc1S(=O)(=O)Nc1ccccc1C(=O)O | Training |
| I68 | INACTIVE | COc1ccc(C)cc1S(=O)(=O)Nc1ccccc1C(=O)O | Training |
| I69 | INACTIVE | COc1ccc(F)cc1S(=O)(=O)Nc1ccc(C(=O)O)cc1 | Training |
| I70 | INACTIVE | COc1ccc(Cl)cc1S(=O)(=O)Nc1ccc(C(=O)O)cc1 | Training |
| I71 | INACTIVE | COc1ccc(Br)cc1S(=O)(=O)Nc1ccc(C(=O)O)cc1 | Training |
| I72 | INACTIVE | COc1cc(C)c(C)cc1S(=O)(=O)Nc1ccc(C(=O)O)cc1 | Training |
| I73 | INACTIVE | COc1ccc(C)cc1S(=O)(=O)Nc1ccc(C(=O)O)cc1 | Training |
| I74 | INACTIVE | COc1cc(C)c(Cl)cc1S(=O)(=O)Nc1ccc(C(=O)O)cc1 | Training |
| I75 | INACTIVE | COc1ccc(OC)c(S(=O)(=O)Nc2ccc(C(=O)O)cc2)c1 | Training |
| I76 | INACTIVE | O=C(O)C1C2CC=CC2c2c3c(cc4ccccc24)C2C=CCC2CN31 | Training |
| I77 | INACTIVE | CC(CC(=O)O)c1ccc(N)cc1 | Training |
| I78 | INACTIVE | CC(CC(=O)O)c1ccc(O)c([N+](=O)[O-])c1 | Training |
| I79 | INACTIVE | C=CC(=O)Oc1cc(O)ccc1C(C)CC(=O)O | Training |
| I80 | INACTIVE | O=C(O)CC(c1ccccc1)c1ccc(O)cc1 | Training |
| I81 | INACTIVE | Cc1cccc(C(Cc2ccc(O)cc2)C(=O)O)c1 | Training |
| I82 | INACTIVE | CCOc1cc(CCC(=O)O)ccc1O | Training |
| I83 | INACTIVE | Cc1cc(O)ccc1C(CC(=O)O)c1ccccc1 | Training |
| I84 | INACTIVE | COc1cc(O)ccc1C(CC(=O)O)c1ccc(O)cc1 | Training |
| I85 | INACTIVE | O=C(O)C(Cc1ccc(O)cc1)c1ccc(O)cc1 | Training |
| I86 | INACTIVE | O=C(O)C(Cc1ccccc1)c1ccc(O)cc1 | Training |
| I87 | INACTIVE | O=C(O)CCc1ccc(O)cc1 | Training |
| I88 | INACTIVE | COc1ccc(CCC(=O)O)cc1C | Training |
| I89 | INACTIVE | CCc1cc(CCC(=O)O)ccc1OC | Training |
| I90 | INACTIVE | CCc1cc(CCC(=O)O)ccc1O | Training |
| I91 | INACTIVE | Cc1cc(CCC(=O)O)ccc1O | Training |
| I92 | INACTIVE | O=C(O)CCc1ccc2c(c1)OCCO2 | Training |
| I93 | INACTIVE | CCc1cc(O)ccc1CCC(=O)O | Training |
| I94 | INACTIVE | CC(C)c1cc(O)ccc1CCC(=O)O | Training |
| I95 | INACTIVE | Cc1cc(O)ccc1CCC(=O)O | Training |
| I96 | INACTIVE | O=C(O)CCc1ccc2c(c1)CC=N2 | Training |
| I97 | INACTIVE | c1cc(CSc2nc3ccccc3[nH]2)nc(CSc2nc3ccccc3[nH]2)c1 | Training |
| I98 | INACTIVE | COC(=O)c1cc(OC)c(OC)cc1NC(=O)CSCC(=O)O | Training |
| I99 | INACTIVE | Cc1ccc(SC(CC(=O)c2ccc(C)cc2)C(=O)O)cc1 | Training |
| I100 | INACTIVE | O=[N+]([O-])c1ccc(S(=O)(=O)c2ccccc2)cc1OS(=O)(=O)c1cc(Cl)cc(Cl)c1 | Training |
| I101 | INACTIVE | COc1ccc(-c2c(C(F)(F)F)nn(C)c2NC(=O)CSCC(=O)O)cc1 | Training |
| A102 | ACTIVE | S=c1[nH]nc(-c2c(N3CCCCC3)nc3ccc(Cl)cc3c2-c2ccccc2)o1 | Validation |
| A103 | ACTIVE | O=C(O)c1sccc1S(=O)(=O)n1ccc2cc(Br)ccc21 | Validation |
| A104 | ACTIVE | COc1c(-c2cccc(N)c2)cc(S(=O)(=O)Nc2ccc(F)c(C(=O)O)c2)c2ccccc12 | Validation |
| A105 | ACTIVE | Cc1cccc2c1ccn2S(=O)(=O)c1ccsc1C(=O)O | Validation |
| A106 | ACTIVE | Cc1c(C(C)(C)C)ccc(S(=O)(=O)Nc2cccc(C(=O)O)c2)c1C | Validation |
| A107 | ACTIVE | O=C(c1nc2ccccc2nc1O)N1CCCC1C(O)c1cccc(Cl)c1 | Validation |
| A108 | ACTIVE | O=C(O)c1sccc1S(=O)(=O)n1ccc2ccc(F)cc21 | Validation |
| A109 | ACTIVE | CCCCCCCCCCOc1ccc(S(=O)(=O)Nc2ccc(F)c(C(=O)O)c2)c2ccccc12 | Validation |
| A110 | ACTIVE | COc1c(-c2ccc(F)c(F)c2)cc(S(=O)(=O)Nc2ccc(F)c(C(=O)O)c2)c2ccccc12 | Validation |
| A111 | ACTIVE | Cc1cn(S(=O)(=O)c2ccccc2C(=O)O)c2ccccc12 | Validation |
| A112 | ACTIVE | CCCCCCOc1ccc(S(=O)(=O)Nc2ccc(F)c(C(=O)O)c2)c2ccccc12 | Validation |
| A113 | ACTIVE | O=C(O)c1sccc1S(=O)(=O)n1ccc2cc(F)ccc21 | Validation |
| A114 | ACTIVE | Cc1ccc2ccn(S(=O)(=O)c3csc(C(=O)O)c3)c2c1 | Validation |
| A115 | ACTIVE | Cc1ccc2c(ccn2S(=O)(=O)c2ccsc2C(=O)O)c1 | Validation |
| A116 | ACTIVE | Clc1ccc2nc(N3CCCCC3)c(-c3nnn[nH]3)c(-c3ccccc3)c2c1 | Validation |
| A117 | ACTIVE | CCCCCOc1ccc(S(=O)(=O)Nc2ccc(F)c(C(=O)O)c2)c2ccccc12 | Validation |
| A118 | ACTIVE | COc1cc(C=CC2SC(=S)N(CCC(=O)O)C2=O)ccc1O | Validation |
| A119 | ACTIVE | O=C(O)c1cc(NS(=O)(=O)c2ccc(OC3CCCCCC3)c3ccccc23)ccc1F | Validation |
| A120 | ACTIVE | CCOc1ccc(S(=O)(=O)Nc2ccc(C(=O)O)cc2)c2ccccc12 | Validation |
| A121 | ACTIVE | COc1c(-c2ccc(Cl)c(Cl)c2)cc(S(=O)(=O)Nc2ccc(F)c(C(=O)O)c2)c2ccccc12 | Validation |
| A122 | ACTIVE | NC(=O)c1ccccc1Cn1c2c(c3cccc(C(=O)O)c31)CCCCC2 | Validation |
| A123 | ACTIVE | O=C(O)c1cc(NS(=O)(=O)c2ccc(OC3CCCCC3)c3ccccc23)ccc1F | Validation |
| A124 | ACTIVE | COc1c(-c2ccoc2)cc(S(=O)(=O)Nc2ccc(F)c(C(=O)O)c2)c2ccccc12 | Validation |
| A125 | ACTIVE | COc1ccc(S(=O)(=O)Nc2ccc(F)c(C(=O)O)c2)c2ccccc12 | Validation |
| A126 | ACTIVE | COc1ccc2ccn(S(=O)(=O)c3ccsc3C(=O)O)c2c1 | Validation |
| A127 | ACTIVE | Cc1ccc(-c2cc(-c3ccccc3OCCCC(=O)O)nn2-c2ccccc2)cc1 | Validation |
| A128 | ACTIVE | COc1c(-c2ccccc2)cc(S(=O)(=O)Nc2ccc(F)c(C(=O)O)c2)c2ccccc12 | Validation |
| A129 | ACTIVE | Cc1c(Cl)cccc1OCc1cc(=O)n2nc(-c3ccccc3)nc2[nH]1 | Validation |
| A130 | ACTIVE | COc1c(-c2ccc(F)c(Cl)c2)cc(S(=O)(=O)Nc2ccc(F)c(C(=O)O)c2)c2ccccc12 | Validation |
| A131 | ACTIVE | O=C(O)c1c(C2CC2)nc2ccc(Cl)cc2c1-c1ccccc1 | Validation |
| A132 | ACTIVE | COc1ccc(S(=O)(=O)Nc2cccc(C(=O)O)c2)c2ccccc12 | Validation |
| A133 | ACTIVE | c1cc(-c2c3c(nc(C4CCCCC4)c2-c2nnn[nH]2)CCCCC3)ccn1 | Validation |
| A134 | ACTIVE | O=C(CSc1nc(O)cc(C(F)(F)F)n1)N1CCCCC1 | Validation |
| A135 | ACTIVE | COc1ccc(Sc2nc(O)cc(C(F)(F)F)n2)cc1 | Validation |
| A136 | ACTIVE | Cc1cc(-c2c3c(nc(C4CCCCC4)c2-c2nnn[nH]2)CCC(C(F)(F)F)C3)ccn1 | Validation |
| A137 | ACTIVE | O=C(O)COc1ccc(-c2ccccc2-n2cc(-c3ccccc3)c(-c3ccccc3)n2)cc1 | Validation |
| A138 | ACTIVE | CCc1c(-c2ccccc2)c(-c2ccc(F)cc2)nn1-c1ccccc1-c1cccc(OCC(=O)O)c1 | Validation |
| A139 | ACTIVE | O=C(O)CCOc1ccccc1-c1cc(-c2ccc(Br)cc2)n(-c2ccccc2)n1 | Validation |
| A140 | ACTIVE | O=C(O)CNc1cccc(-c2ccccc2-c2nc(-c3ccccc3)c(-c3ccccc3)[nH]2)c1 | Validation |
| A141 | ACTIVE | COC(=O)c1sc(-c2cccs2)cc1NC(=O)C=CC(=O)O | Validation |
| A142 | ACTIVE | O=C(O)CCCCOc1ccccc1-c1cc(-c2ccccc2)n(-c2ccccc2)n1 | Validation |
| A143 | ACTIVE | CCc1ccc(OC)c(S(=O)(=O)Nc2cccc(C(=O)O)c2)c1 | Validation |
| A144 | ACTIVE | Cc1cc(-c2c3c(nc(C4(C)CCCC4)c2-c2nnn[nH]2)CCCCC3)ccn1 | Validation |
| A145 | ACTIVE | O=C(O)CCCCOc1ccccc1-c1nc(-c2ccccc2)c(-c2ccccc2)o1 | Validation |
| A146 | ACTIVE | CCCCCC1CCc2nc(C3(COC)CCCC3)c(-c3nnn[nH]3)c(-c3ccccc3)c2C1 | Validation |
| A147 | ACTIVE | O=C(O)COc1cccc(-c2nc(-c3ccccc3)c(-c3ccccc3)o2)c1 | Validation |
| A148 | ACTIVE | Cc1cc(C(C)C)cc(S(=O)(=O)Nc2ccccc2C(=O)O)c1C | Validation |
| A149 | ACTIVE | CCn1c(-c2ccccc2-c2cccc(OCC(=O)O)c2)nc(-c2ccccc2)c1-c1ccccc1 | Validation |
| A150 | ACTIVE | CC1COc2c3c(cc(F)c2N2CCN(C)CC2)C(=O)C(C(=O)O)=CC31 | Validation |
| A151 | ACTIVE | COc1ccc(C(C)(C)C)cc1S(=O)(=O)Nc1ccc(C(=O)O)cc1 | Validation |
| A152 | ACTIVE | O=c1cc(COc2cccc(Cl)c2C2CC2)[nH]c2nc(-c3ccccc3)nn12 | Validation |
| A153 | ACTIVE | Cc1ccc(-c2cc(-c3ccccc3OCCC(=O)O)nn2-c2ccccc2)cc1 | Validation |
| A154 | ACTIVE | O=c1[nH]nc(-c2c(N3CCCCC3)nc3ccc(Cl)cc3c2-c2ccccc2)o1 | Validation |
| A155 | ACTIVE | O=C(O)Cc1cn(-c2ccccc2-c2nc(-c3ccccc3)c(-c3ccccc3)s2)nn1 | Validation |
| A156 | ACTIVE | COc1cc(C)c(C(C)C)cc1S(=O)(=O)Nc1cccc(C(=O)O)c1 | Validation |
| A157 | ACTIVE | Cc1cc(-c2c3c(nc(C4(CO)CCCC4)c2-c2nnn[nH]2)CCCCC3)ccn1 | Validation |
| A158 | ACTIVE | O=C(Nc1cc(Cl)cc(Cl)c1-c1ccccc1)NC1(C(=O)O)CCCC1 | Validation |
| A159 | ACTIVE | CC(C)OC(=O)C(C(=O)C(F)(F)F)c1cc(NS(=O)(=O)c2ccccc2)ccc1O | Validation |
| A160 | ACTIVE | COc1ccc(S(=O)(=O)Nc2ccc(C(=O)O)cc2)c2ccccc12 | Validation |
| A161 | ACTIVE | O=C(O)CCCOc1ccccc1-c1cc(-c2ccccc2)n(-c2cccc(Cl)c2)n1 | Validation |
| A162 | ACTIVE | O=C(Nc1cc(Cl)c(Cl)cc1-c1ccccc1)NC1(C(=O)O)CCCC1 | Validation |
| A163 | ACTIVE | CCCCCC=CCC=CCCCCCCCC(=O)O | Validation |
| A164 | ACTIVE | CCc1c(-c2ccccc2)c(-c2ccccc2)nn1-c1ccccc1-c1cccc(C(O)CC(=O)O)c1 | Validation |
| A165 | ACTIVE | CCCCCC=CCC=CCC=CCC=CCCCC(=O)O | Validation |
| A166 | ACTIVE | COc1ccc(S(=O)(=O)Nc2cccc(C(=O)O)c2)c(C)c1C | Validation |
| A167 | ACTIVE | CCOc1ccc(S(=O)(=O)Nc2cccc(C(=O)O)c2)cc1C(C)(C)C | Validation |
| A168 | ACTIVE | CCCCCCCCCCCCCCCC(=O)O | Validation |
| A169 | ACTIVE | O=C(O)CCc1ccc(O)c(C(=O)c2ccccc2)c1 | Validation |
| A170 | ACTIVE | COc1ccc(Cn2c3c(c4cccc(C(=O)O)c42)CCCC3)cc1 | Validation |
| A171 | ACTIVE | O=C(O)c1cccc2c3c(n(Cc4c(F)cccc4C(F)(F)F)c12)CCCC3 | Validation |
| A172 | ACTIVE | NC(=O)c1cccc(Cn2c3c(c4cccc(C(=O)O)c42)CCCCC3)c1 | Validation |
| A173 | ACTIVE | O=C(O)c1cccc2c3c(n(Cc4cccc(C(F)(F)F)c4)c12)CCCCC3 | Validation |
| A174 | ACTIVE | O=C(O)c1cccc2c3c(n(Cc4ccc(F)cc4)c12)CCCC3 | Validation |
| A175 | ACTIVE | O=C(O)c1cccc2c3c(n(Cc4ccccc4F)c12)CCCC3 | Validation |
| A176 | ACTIVE | O=C(O)c1cccc2c3c(n(Cc4cccc(C(F)(F)F)c4)c12)CCCC3 | Validation |
| A177 | ACTIVE | CCc1c(-c2ccc(F)cc2)c(-c2ccccc2)nn1-c1ccccc1-c1cccc(OCC(=O)O)c1 | Validation |
| A178 | ACTIVE | O=C(O)c1cccc2c3c(n(Cc4ccccc4C(F)(F)F)c12)CCCCC3 | Validation |
| A179 | ACTIVE | O=C(O)c1cccc2c3c(n(Cc4ccccc4)c12)CCCC3=NO | Validation |
| A180 | ACTIVE | O=C(O)c1cccc2c3c(n(Cc4ccccc4)c12)CCCC3 | Validation |
| A181 | ACTIVE | O=C(O)c1cccc2c3c(n(Cc4ccccc4C(F)(F)F)c12)CCCC3 | Validation |
| A182 | ACTIVE | O=C(O)CSc1cc(NS(=O)(=O)c2ccc(Cl)cc2)c2ccccc2c1O | Validation |
| A183 | ACTIVE | Cc1cc2c3c(c1)C1C=CCC1C(C(=O)O)N3CC1CC=CC21 | Validation |
| A184 | ACTIVE | O=C(O)CCN1C2C=CC=CC2C2C=CC=CC21 | Validation |
| A185 | ACTIVE | O=C(O)C1C2CC=CC2c2cc(Br)cc3c2N1CC1CC=CC31 | Validation |
| A186 | ACTIVE | O=C(O)COc1cccc(-c2ccccc2-n2cc(-c3ccccc3)c(-c3ccccc3)n2)c1 | Validation |
| A187 | ACTIVE | O=C(O)CCCCN1C2C=CC=CC2C2C=CC=CC21 | Validation |

**Supplementary section E: Dataset partition**

A balanced training set was representatively sampled from the dataset to avoid any bias towards the overrepresented category. The compiled dataset consisted of 187 compounds from the ACTIVE category and 101 compounds from the INACTIVE category. In order to have a balanced training set with the highest possible number of compounds in each category, all INACTIVE compounds were allocated to this set. To ensure a representative partition of the ACTIVE compounds into training and external validation sets, iRaPCA subspace clustering approach was used.

The following values were used for the tunable parameters of iRaPCA: variance cutoff for descriptors: 0.05; number of subspaces: 100; number of descriptors per subspace: 200; correlation coefficient cutoff between descriptor pairs: 0.4; minimum number of descriptors per subspace: 4; maximum number of descriptors per subspace: 25; minimum number of clusters per iteration: 2; maximum number of clusters per iteration: 25; maximum ratio of molecules per cluster/total molecules: 1; number of iterations: 1; number of principal components to use: 2.

The iRaPCA clustering on the 187 ACTIVE compounds resulted in four clusters with 69, 81, 25, and 12 compounds. Training instances were sampled representatively from each cluster proportionally to the cluster, until a 101-compound set of ACTIVE compounds was obtained. The remaining 86 ACTIVE compounds were used for the generation of decoys, as described in the main text. Final dataset composition is shown in **Table 1** in the main text.

**Supplementary section F: Docking parameters**

**Table S4.** Parameters implemented in the docking runs.

| **ALGORITHM** | **PARAMETERS** |
| --- | --- |
| **Vina/Vinardo** | Exhaustiveness=32 |
| **AD GPU/AD Bias** | No (max) of eval per run=2,500,000  No (max) of generations per run=42,000  Population size=150  Mutation rate=2%  Crossover rate=80%  Local-search method= ADADELTA  Local-search iterations (max)=300  Local search rate=100 %  Tournament (selection) rate= 60 % |
