## Supplementary figures and images for "*In silico* drug repurposing and in vitro validation of cestode fatty acid binding proteins"

### FigureS1.png

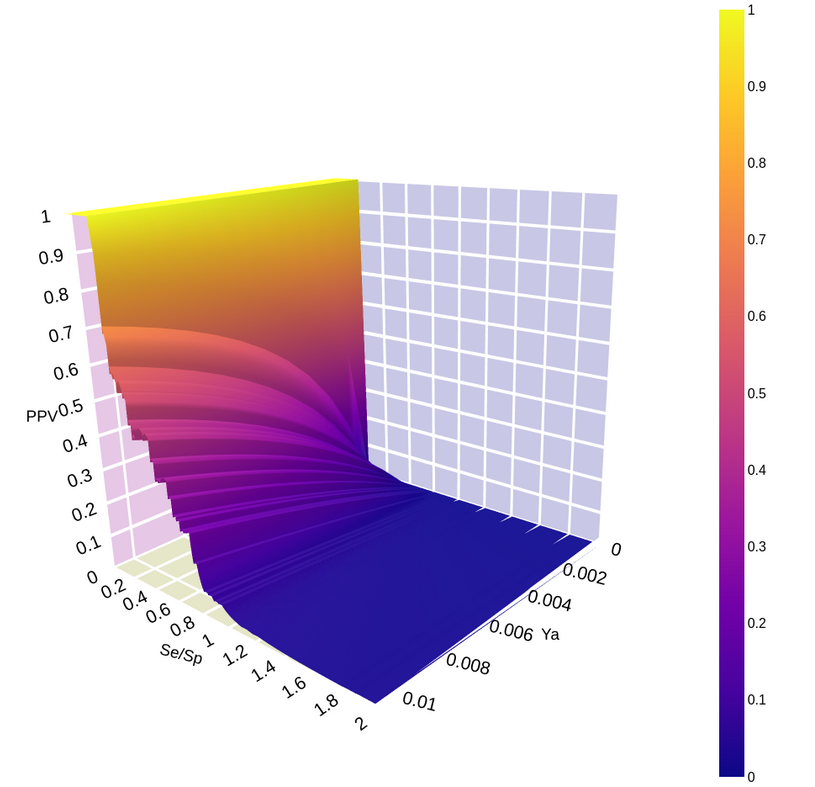

### FigureS2.png

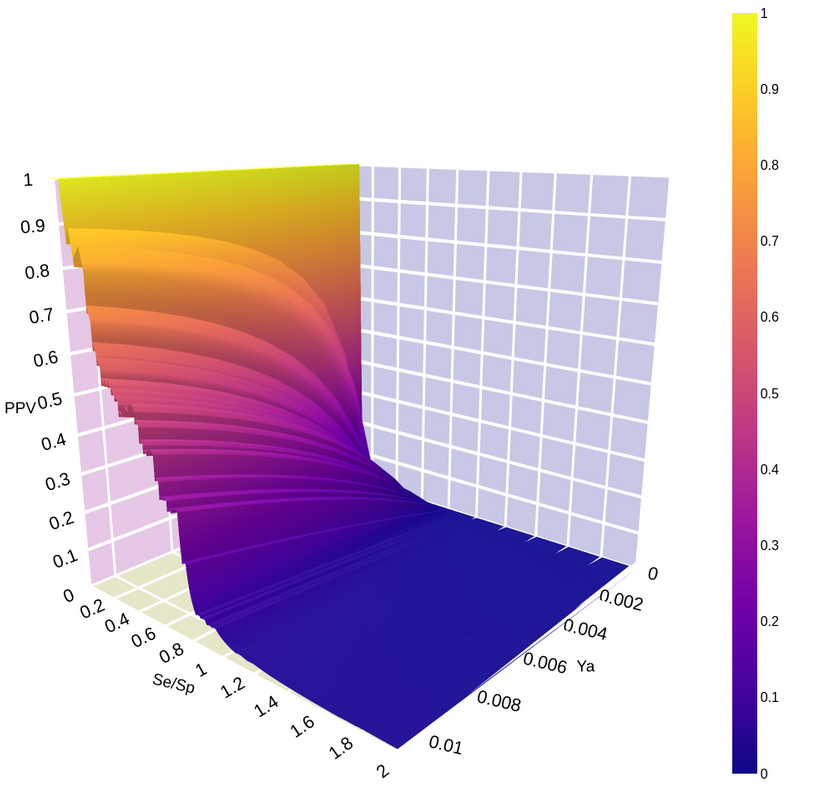

### FigureS3.png

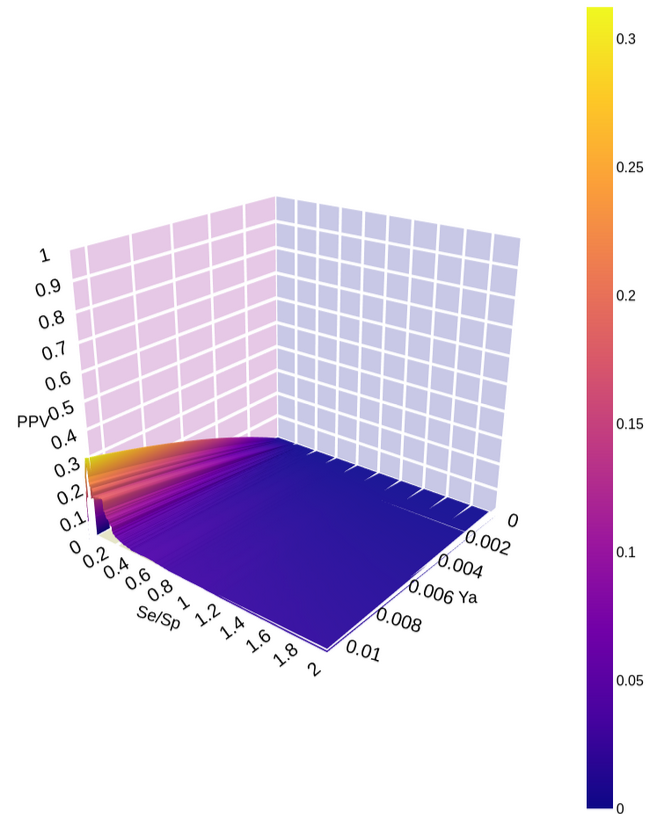

### FigureS4.png

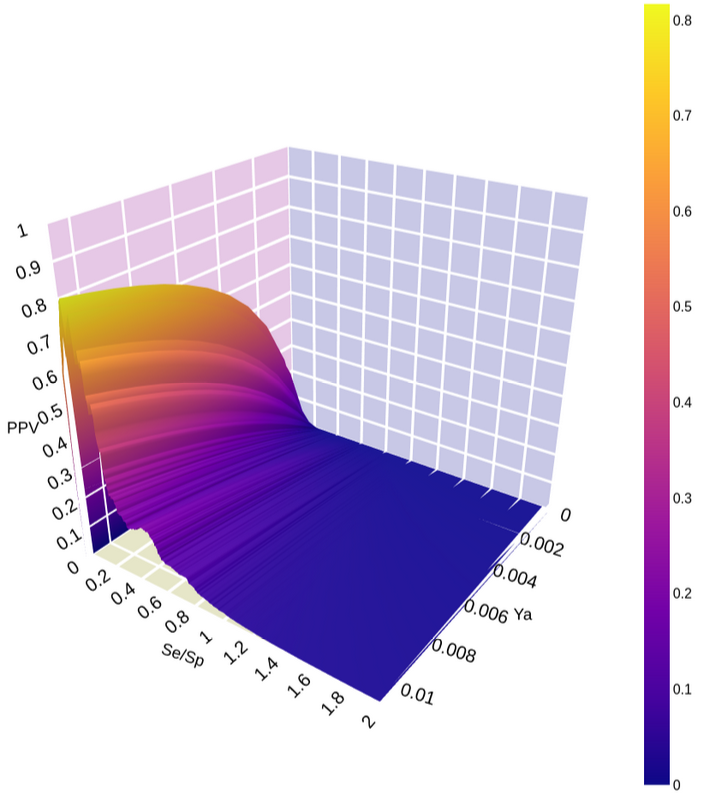

### FigureS5.png

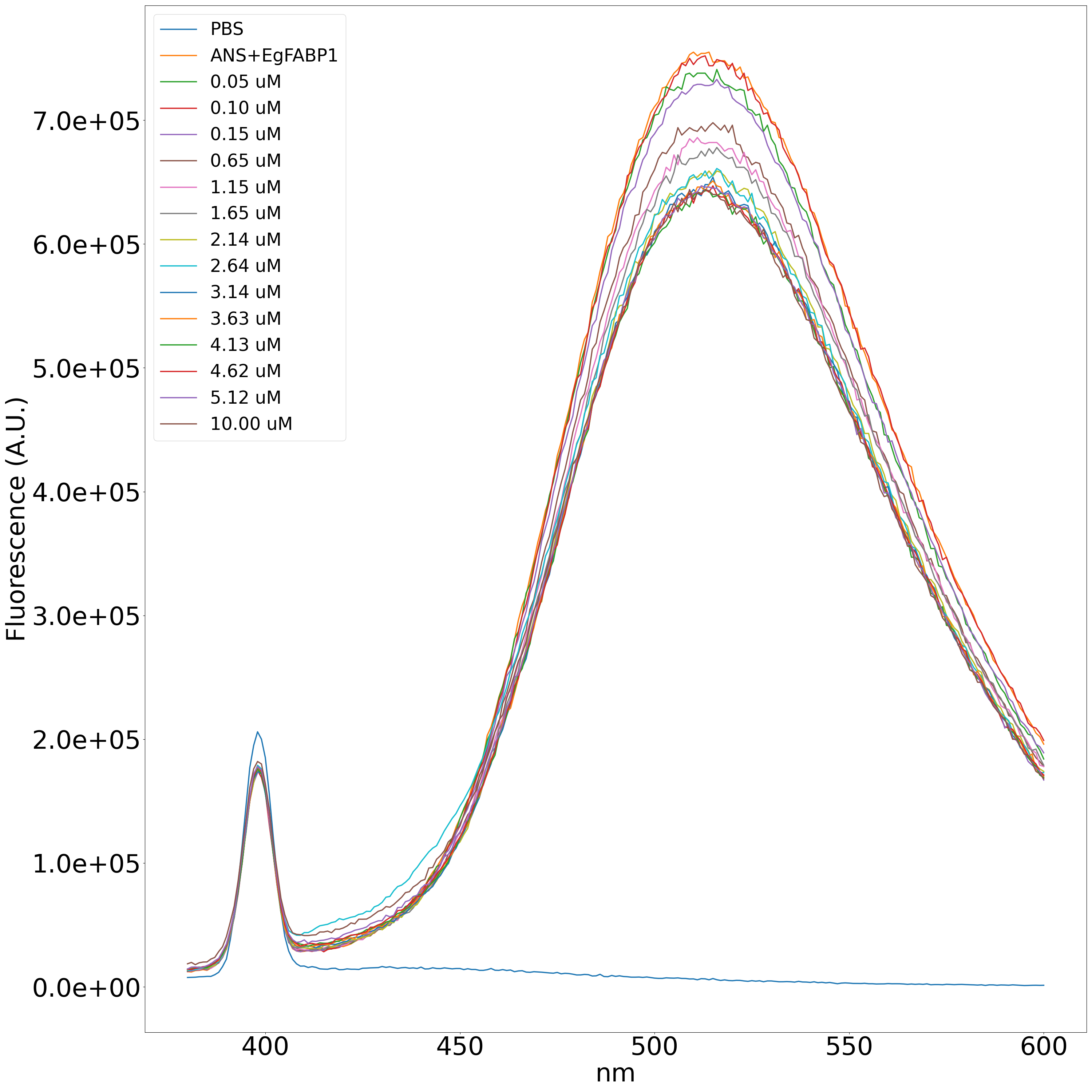

### FigureS6.png

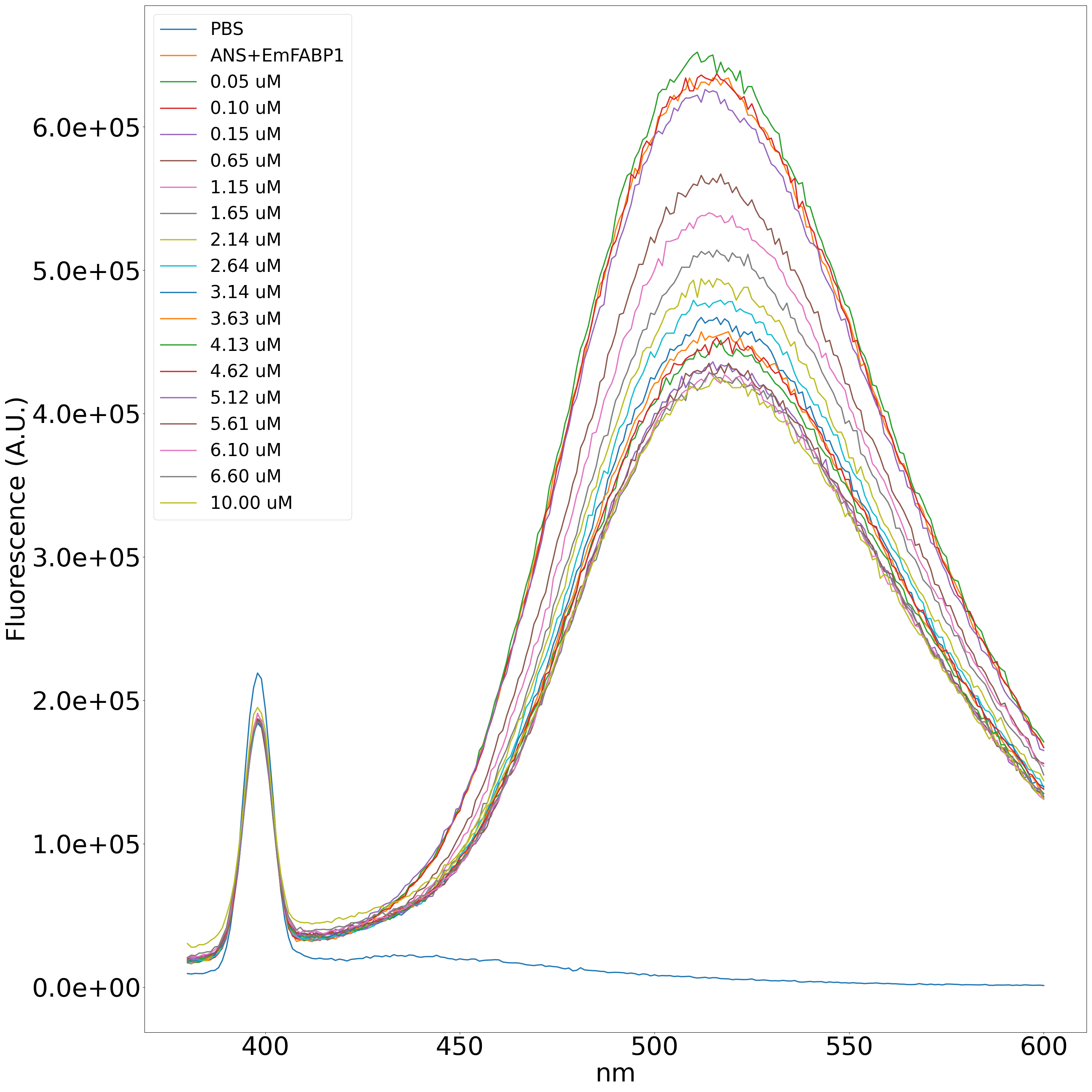

### FigureS7.png

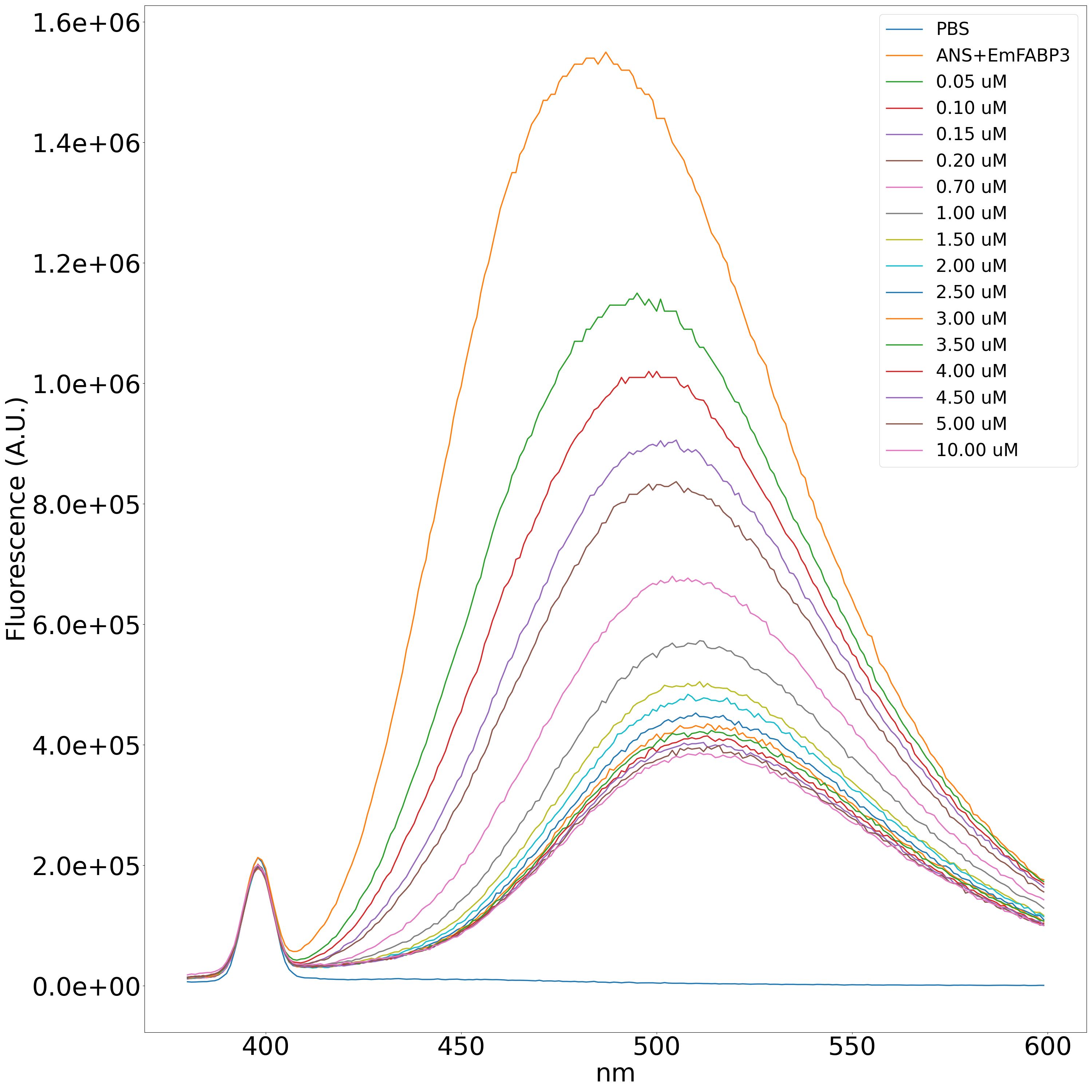

### FigureS8.png

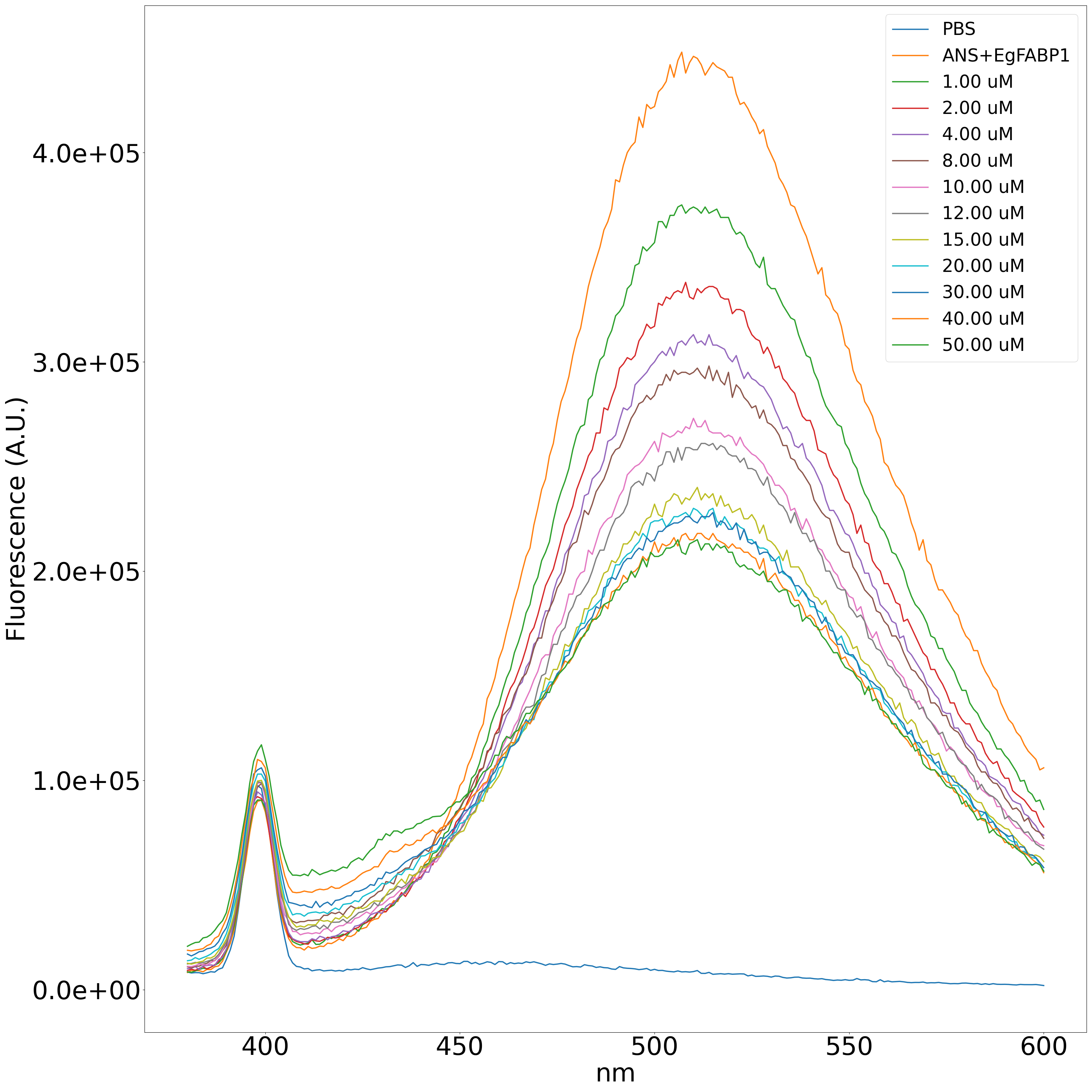

### FigureS9.png

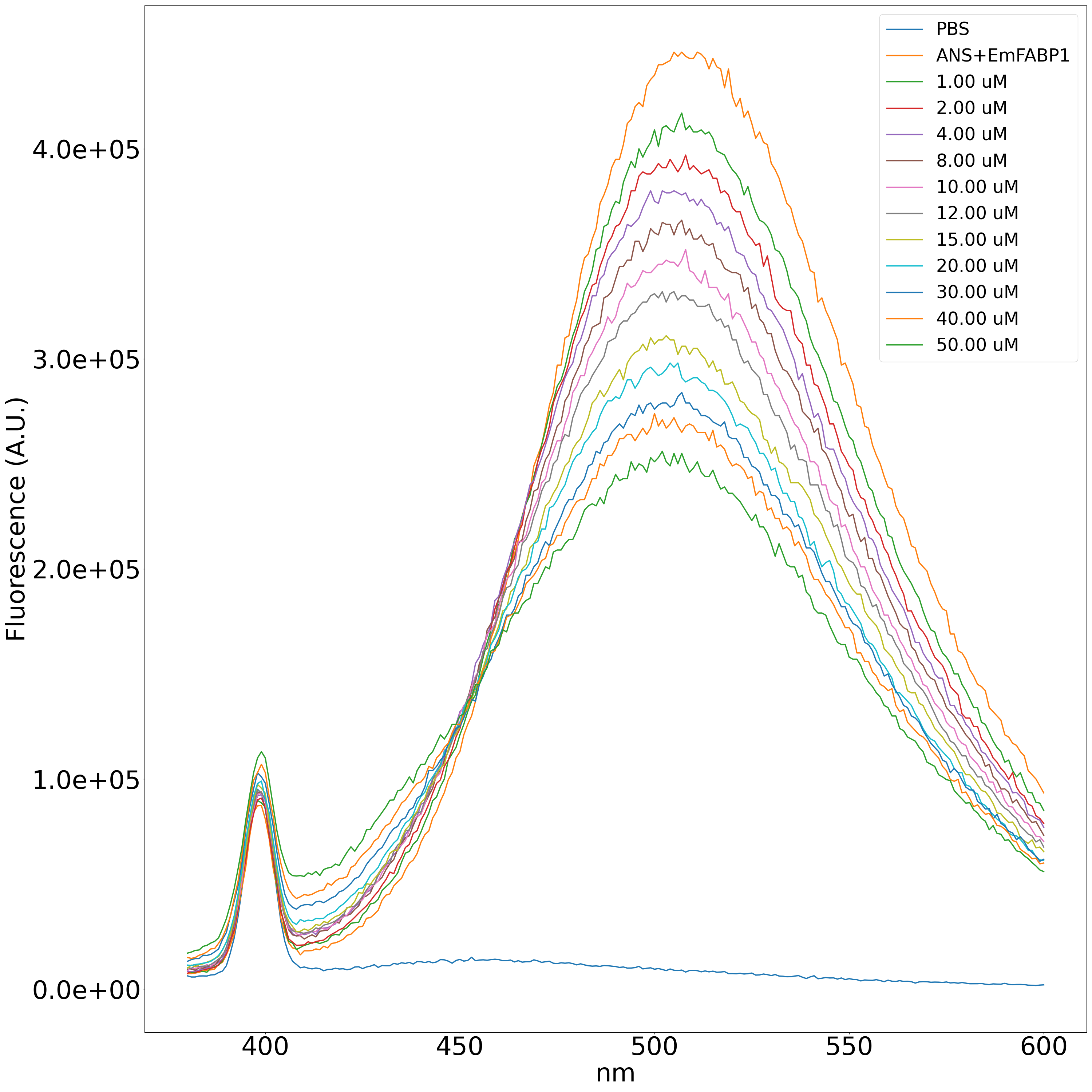

### FigureS10.png

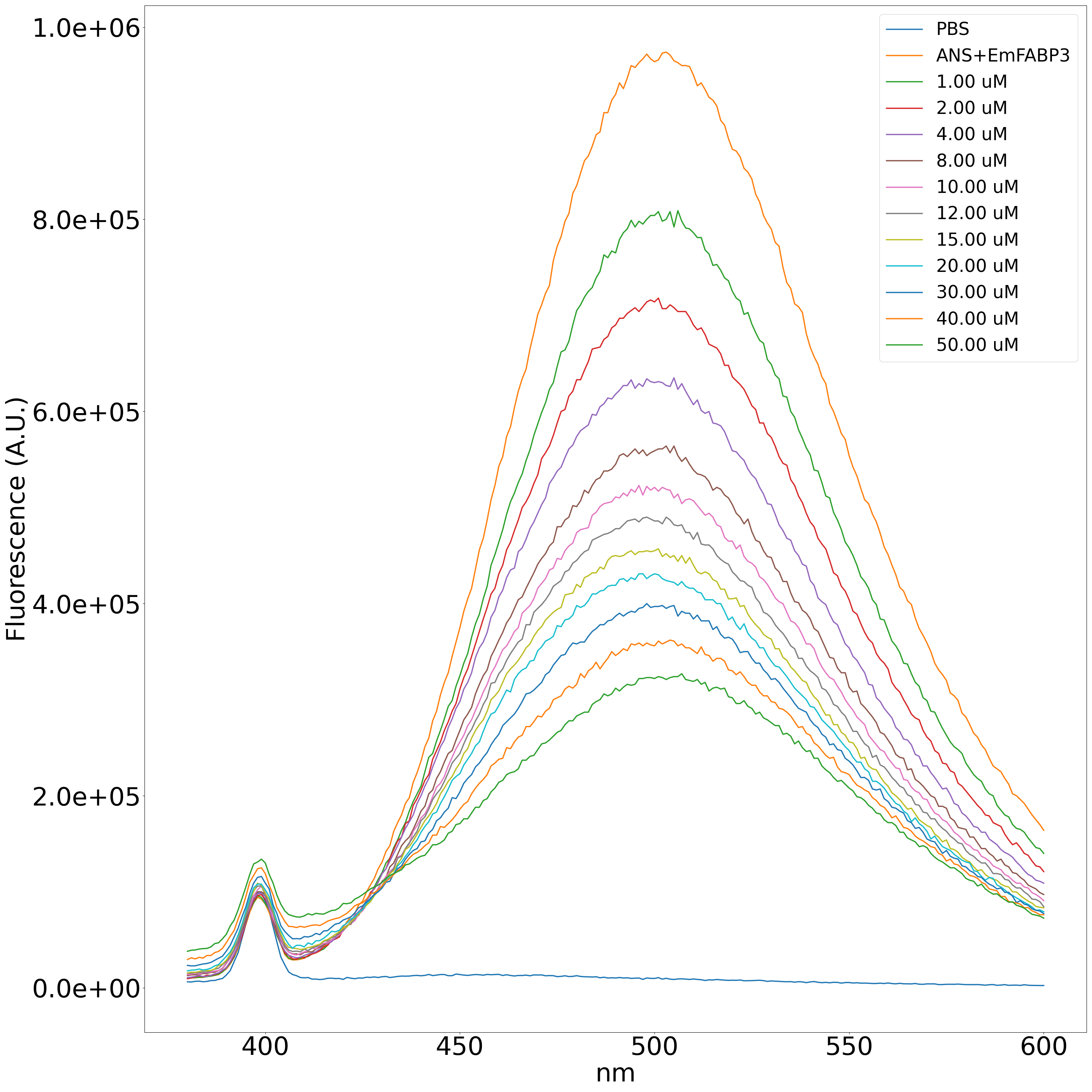

### FigureS11.png

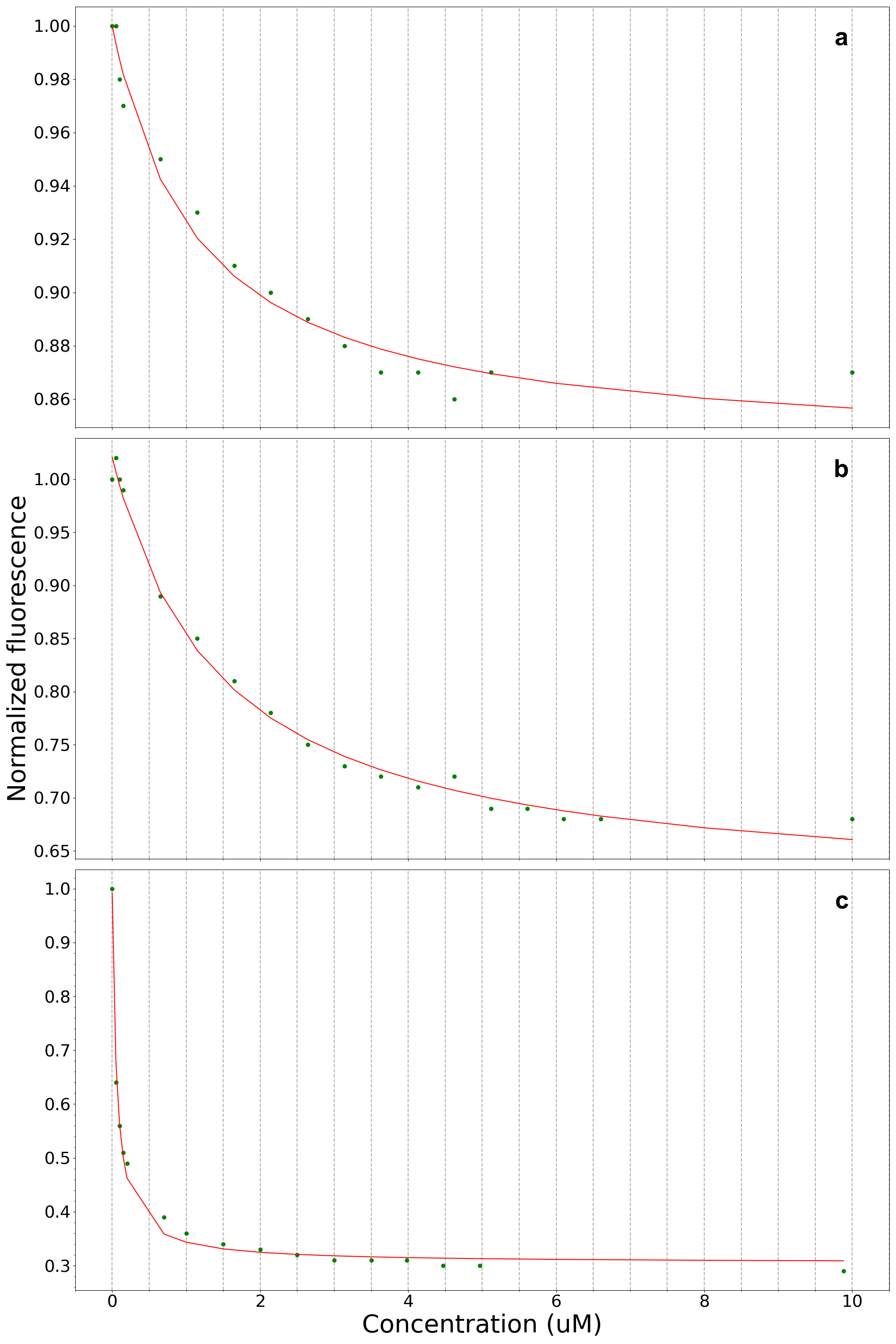

### FigureS12.png

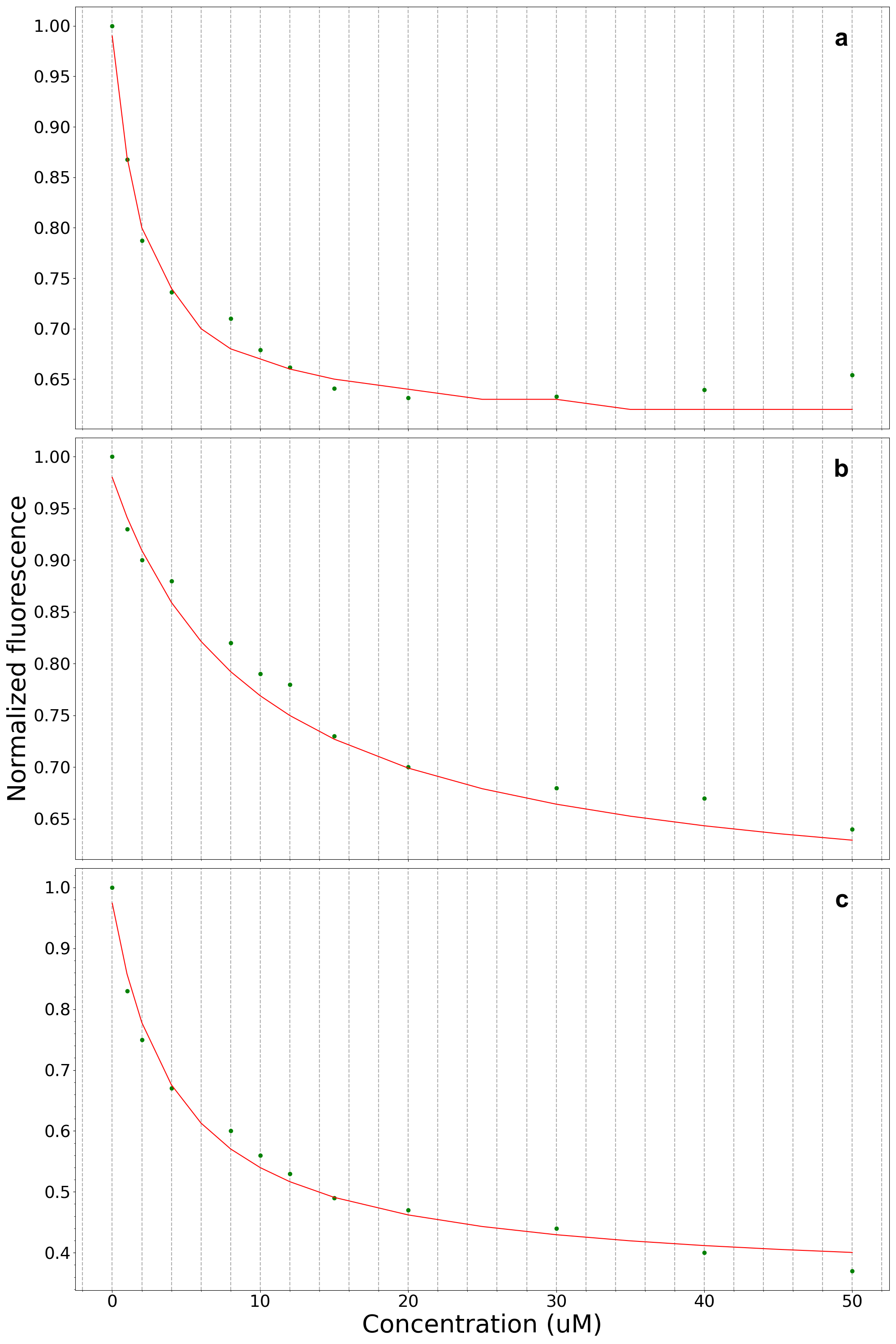

### FigureS13.png

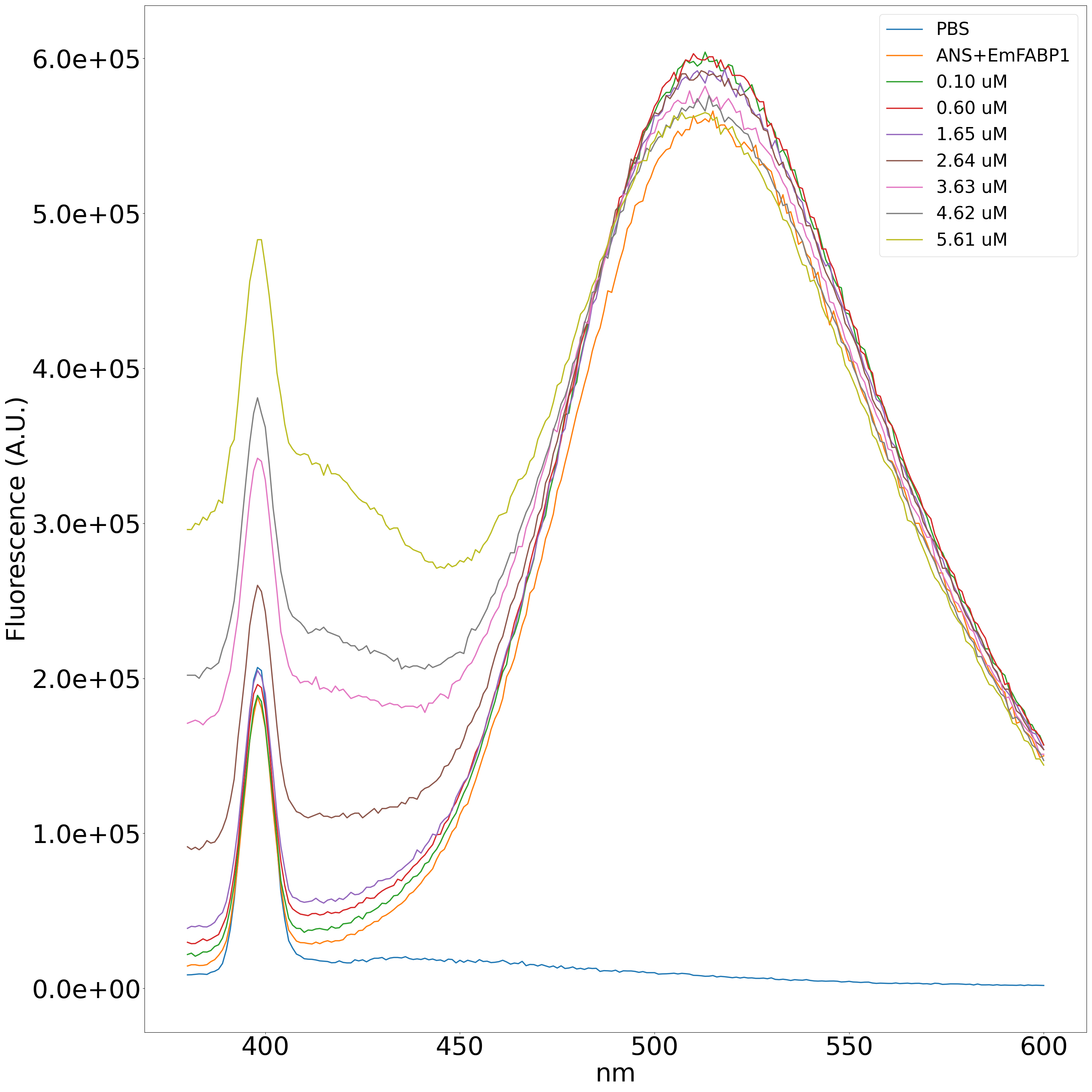

### FigureS14.png

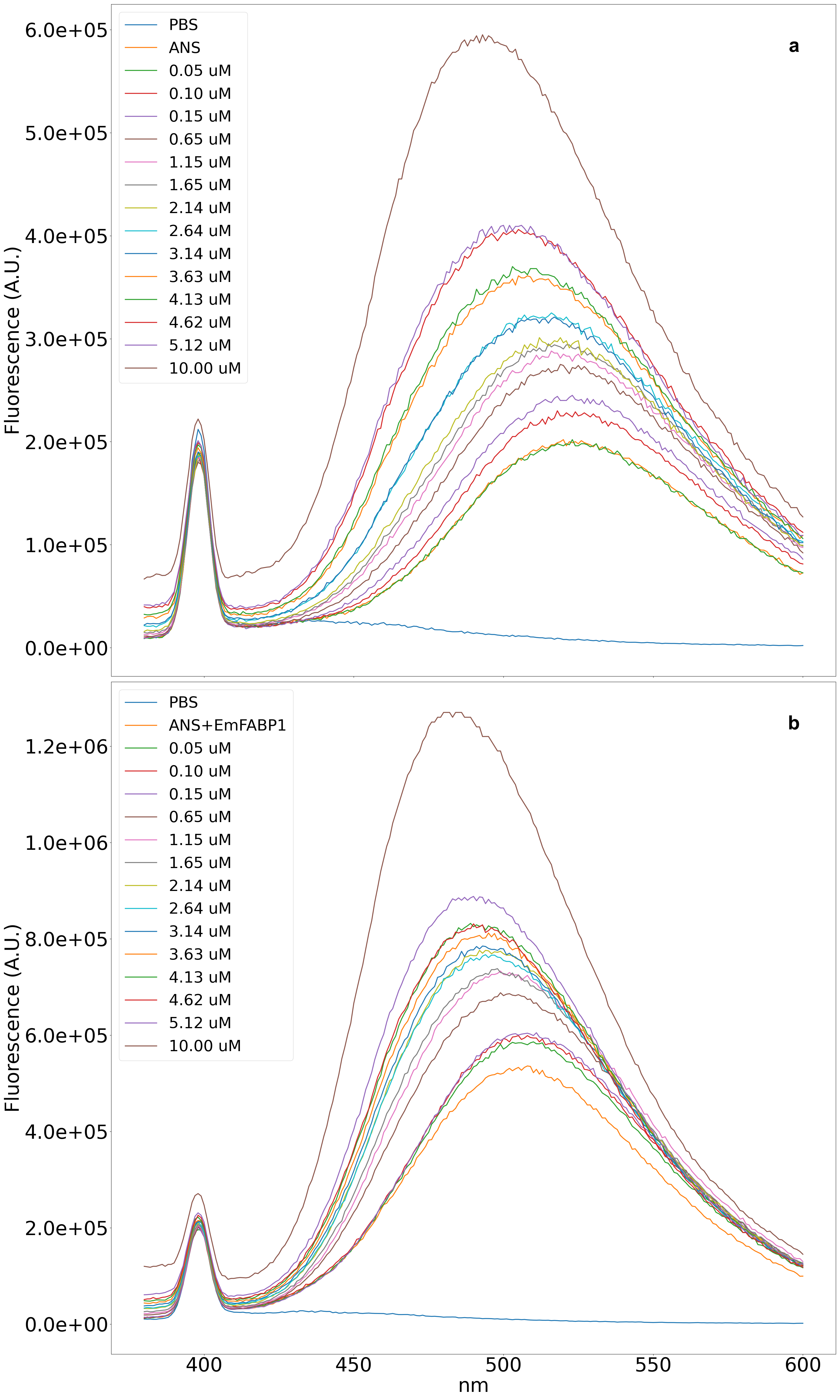

### FigureS15.png

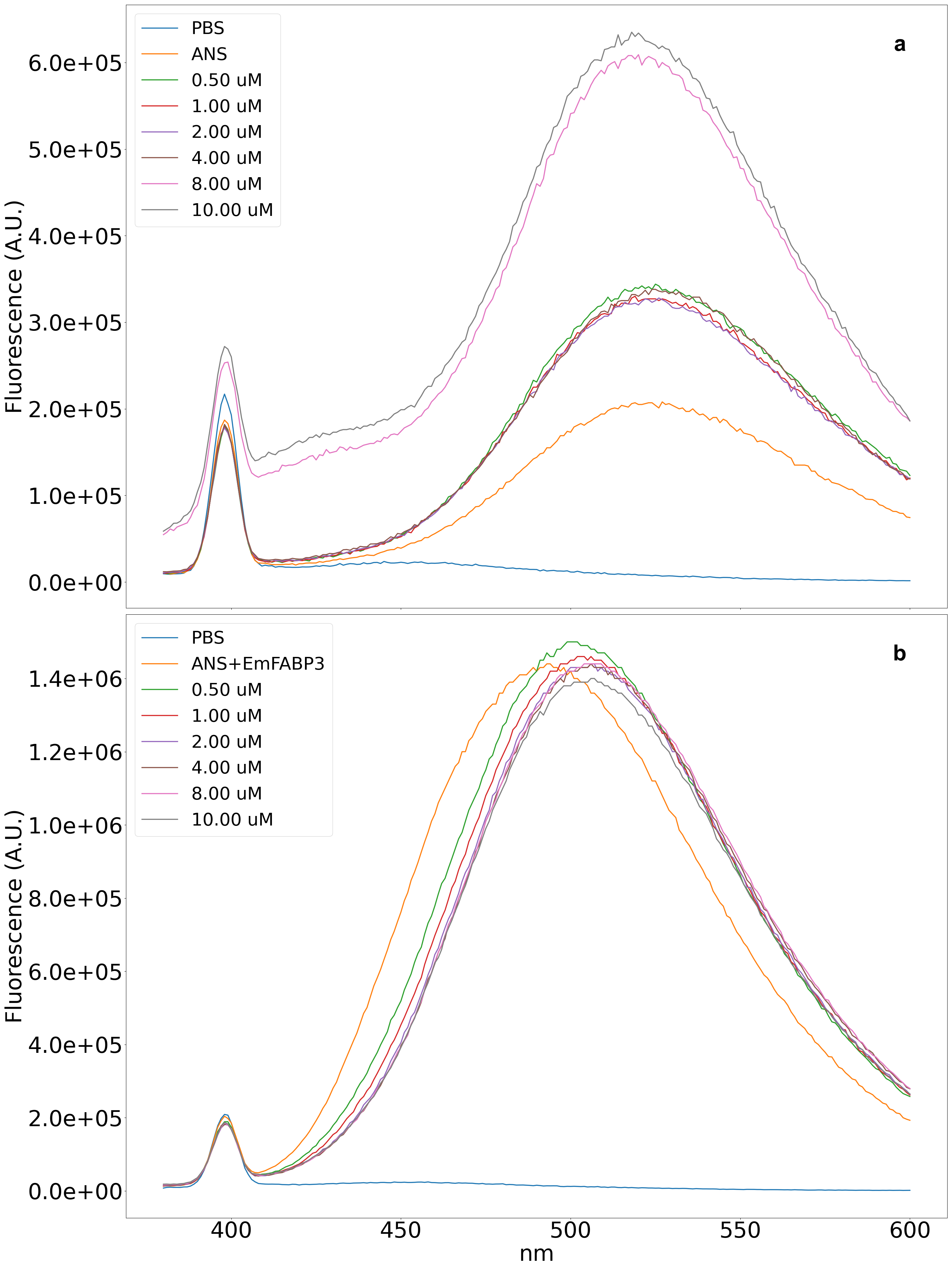

### FigureS16.png

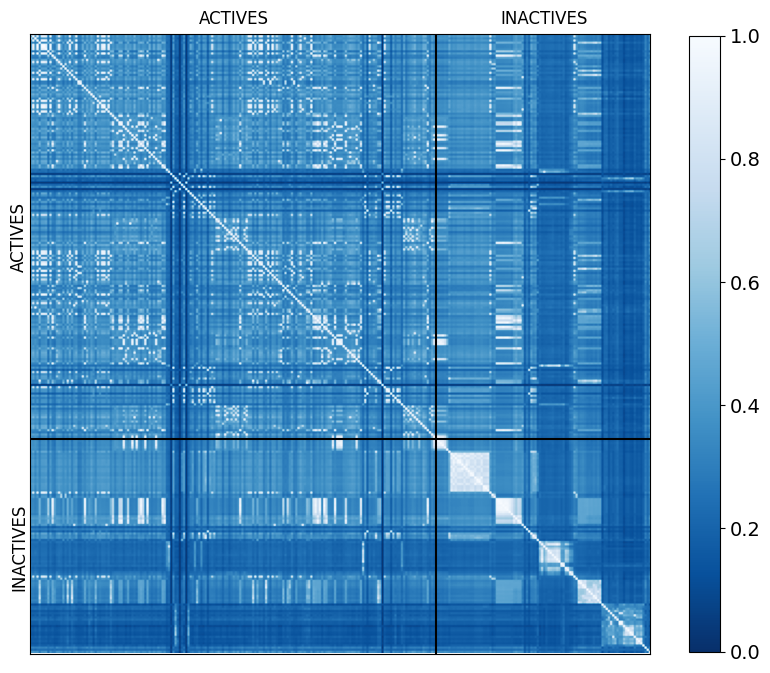
